# Mapping RNA-capsid interactions and RNA secondary structure within authentic virus particles using next-generation sequencing

**DOI:** 10.1101/720425

**Authors:** Yiyang Zhou, Andrew Routh

## Abstract

To characterize RNA-capsid binding sites genome-wide within mature RNA virus particles, we have developed a Next-Generation Sequencing (NGS) platform: Photo-Activatable Ribonucleoside Cross-Linking (PAR-CL). In PAR-CL, 4-thiouracil is incorporated into the encapsidated genomes of authentic virus particles and subsequently UV-crosslinked to adjacent capsid proteins. We demonstrate that PAR-CL can readily and reliably identify capsid binding sites in genomic viral RNA by detecting crosslink-specific uridine to cytidine transitions in NGS data. Using Flock House virus (FHV) as a model system, we identified highly consistent and significant PAR-CL signals across virus RNA genome indicating a clear tropism of the encapsidated RNA genome. Certain interaction sites correlate to previously identified FHV RNA motifs. We additionally performed dimethyl sulfate mutational profiling with sequencing (DMS-MaPseq) to generate a high-resolution profile of single-stranded genomic RNA inside viral particles. Combining PAR-CL and DMS-MaPseq reveals that the predominant RNA-capsid sites favor double-stranded RNA regions. We disrupted secondary structures associated with PAR-CL sites using synonymous mutations, resulting in varied effects to virus replication, propagation, and packaging. Certain mutations showed substantial deficiency in virus replication, suggesting these RNA-capsid sites are multifunctional. These provide further evidence to support that FHV packaging and replication are highly coordinated and inter-dependent events.

**Importance:** Icosahedral RNA viruses must package their genetic cargo into the restrictive and tight confines of the protected virions. High resolution structures of RNA viruses have been solved by Cryo-EM and crystallography, but the encapsidated RNA often eluded visualization due to the icosahedral averaging imposed during image reconstruction. Asymmetrical reconstructions of some icosahedral RNA virus particles have revealed that the encapsidated RNAs conform to specific structures, which may be related to programmed assembly pathway or an energy-minima for RNA folding during or after encapsidation. Despite these advances, determining whether encapsidated RNA genomes conform to a single structure and determining what regions of the viral RNA genome interact with the inner surface of the capsid shell remains challenging. Furthermore, it remains to be determined whether there exists a single RNA structure with conserved topology in RNA virus particles or an ensemble of genomic RNA structures. This is important as resolving these features will inform the elusive structures of the asymmetrically encapsidated genomic material and how virus particles are assembled.

## Introduction

Flock House virus (FHV) is a non-enveloped, single-stranded positive-sense RNA (+ssRNA) virus from the family *Nodaviridae*. The small bipartite genome comprising RNA 1 (3.1kb) and RNA 2 (1.4kb) is packaged into a 34 nm non-enveloped T=3 icosahedral virion. Only two non-structural proteins are produced by FHV: the RNA-dependent RNA polymerase (RdRp) and sub-genomic RNA encoded protein called B2. The B2 protein was discovered as the virus’s approach to evade the invertebrate anti-viral RNA silencing machinery (1, 2), which thereafter led to the discovery of similar mechanisms in plant cells (3). FHV is perhaps the best studied *alphanodavirus* and provides a powerful model system by virtue of its small genome size (4.5kb), genetic tractability and ability to infect Drosophila and mosquito cells in culture and whole flies (reviewed in (4, 5)). More recently, FHV has been adapted into medical field. FHV-related vaccine developments utilized either the viral particle as antibody-display system (6), or the viral RNA as trans-encapsidated chimeric viral vaccine platform (7–9).

Both authentic virions of FHV and the related Pariacoto virus have been reconstructed by cryo-EM and X-ray crystallography to reveal highly ordered dodecahedral cages of RNAs (10, 11). The X-ray structure of FHV virion showed electron density at the icosahedral 2-fold axis, which was modelled as an ordered RNA duplex of approximate 20 nucleotides (12). This would account for 1800nts (more than one third) of the viral genome, implicating a highly-ordered and specific set of interactions between the viral protein capsid and the encapsidated genome. Interestingly, and recombinantly expressed virus-like particles (VLPs) of FHV also exhibit a similar dodecahedral RNA cage despite packaging predominantly cellular RNAs indicating that viral capsid may either impose structure upon the encapsidated RNA or select for natively structured host RNAs such as ribosomal RNAs (13, 14). However, as these structures are obtained with icosahedral averaging, we still do not know what regions or sequences of viral genomic RNA comprise the RNA cage. Furthermore, it remains to be determined whether there exists a single RNA structure with conserved topology in FHV virions, or rather an ensemble of related genomic RNA structures.

The FHV encapsidation process also remains largely unknown. One molecule of each RNA 1 and 2 is specifically encapsidated into virus particles (15), while subgenomic RNA 3 is excluded (16). Several components of the capsid protein such as the arginine–rich motif and the C-terminal FEGFGF motif have been demonstrated to be essential determinants of packaging specificity of RNA 1, RNA 2, or both (17–19). It was also speculated that FHV packaging process may be in close association with viral replication and/or translational events (20–23). In the virus genome, one stem-loop structure in RNA 2 proximal to 5’ end was demonstrated to be required for RNA 2 packaging (24). However, it remains unclear whether there are similar packaging sites on RNA 1 or 2, and how these sites interact and thus recruit capsid protein to fulfill virus encapsidation.

Next-generation sequencing (NGS) in combination with crosslinking techniques provides a high-throughput approach to study transcriptome-wide RNA-protein interactions (reviewed in (25)). A number of new technologies have successfully described interactions between RNA-binding proteins (RBPs) and different types of RNAs, including nascent transcripts, mRNAs, microRNAs and ribosomal RNAs. Among these, PAR-CLIP (Photoactivatable Ribonucleoside-Enhanced Crosslinking and Immunoprecipitation) (26) utilizes a 365 nm UVA-activatable ribonucleoside analog 4-thiouridine (4SU) to effectively crosslink RNA to bound proteins. The enriched crosslinked RNAs result in a highly specific U to C mutation during NGS library preparation (27–29), granting the ability to rapidly identify RBP and microRNA target sites on a transcriptome scale (26).

In an analogous fashion to PAR-CLIP, here we applied the same principles to study the interaction of FHV genomic viral RNA in the context of assembled authentic virions. Unlike the complex cellular micro-environment, authentic virions represent a highly simplified enclosure with few well-defined components (viral RNA and capsid). Therefore, we are able to screen for specific in virion RNA-capsid interaction events without interference from other cellular components. Furthermore, since viruses can be readily separated from other cellular components, we avoided the need of immunoprecipitation for RNA recovery, and thus largely simplify PAR-CLIP methodology. This method is hence named *‘PAR-CL’*.

Using FHV as a model system, PAR-CL methodology was validated by determining that the increased U to C (U-C) mutation rate was highly specific to crosslink between viral RNA and capsid. We noticed that the intensity of PAR-CL signals was subjected to the dose of 4SU and time of incubation. Triplicated FHV PAR-CL experiments revealed significant and highly consistent PAR-CL signals across genome, which implicated a clear tropism of RNA cage inside capsid shell. The multiple clusters of PAR-CL sites suggest that FHV encapsidation may require multiple synergetic packaging sites. DMS-MaPseq (dimethyl sulfate mutational profiling with sequencing) was used to chemically probe single-stranded FHV genomic RNA in virions. We thus constructed a whole genome DMS-MaPseq-imposed RNA secondary structure map for FHV. We noticed RNA-capsid interaction sites favored double stranded RNA regions. Synonymous mutations were designed to disrupt predicted PAR-CL sties in dsRNA regions, which resulted in varied effects to virus RNA replication, propagation, and virulence. Mutations over certain PAR-CL sites showed evidential deficiency in RNA replication, suggesting these sites serve a multifunctional role in both virus packaging and replication. This provides further evidence to support that FHV packaging and replication are highly synchronized and inter-dependent events.

## Results

### Photoactivatable-ribonucleoside-enhanced crosslinking (PAR-CL) in virus particles

PAR-CLIP (Photoactivatable-Ribonucleoside-Enhanced Crosslinking and Immunoprecipitation) is a well-established method for identification of RNA-protein binding sites and can provide nucleotide-resolution through analysis of specific uridine to cytosine transitions that occur at the site of RNA-crosslinking during cDNA synthesis (26, 30, 31). Here, we simplified the technique by applying a similar approach to purified authentic virions of RNA virus, thereby removing the necessity of immunoprecipitation, and hence deriving the name “PAR-CL”. A schematic of the process is illustrated in **Figure 1**. In PAR-CL, photoactivable nucleotide 4-thiouridine (4SU) was provided to cells in culture prior to infection with Flock House virus (FHV). 4SU is rapidly taken by cultured cells without significant cytotoxic effect (26, 32). Upon infection, 4SU is randomly incorporated into newly synthesized viral RNA, which is subsequently packaged into authentic virus particles of FHV. Virus particles were isolated using established methods for virus purification by ultracentrifugation (13). Next, purified virus particles were subjected to UV 365 nm irradiation, yielding crosslinks between the thio-group of 4SU in the viral genomic RNA and amino acid residues in the protein capsid shell only if they are in close proximity (32, 33). Virus particles were then disrupted by proteinase K treatment and a pool of viral RNAs with varied crosslinking sites was obtained. We then generated canonical random-primed RNAseq libraries using ClickSeq (34–36) for Illumina SE150 read sequencing on a MiSeq platform.

**Figure 1.**
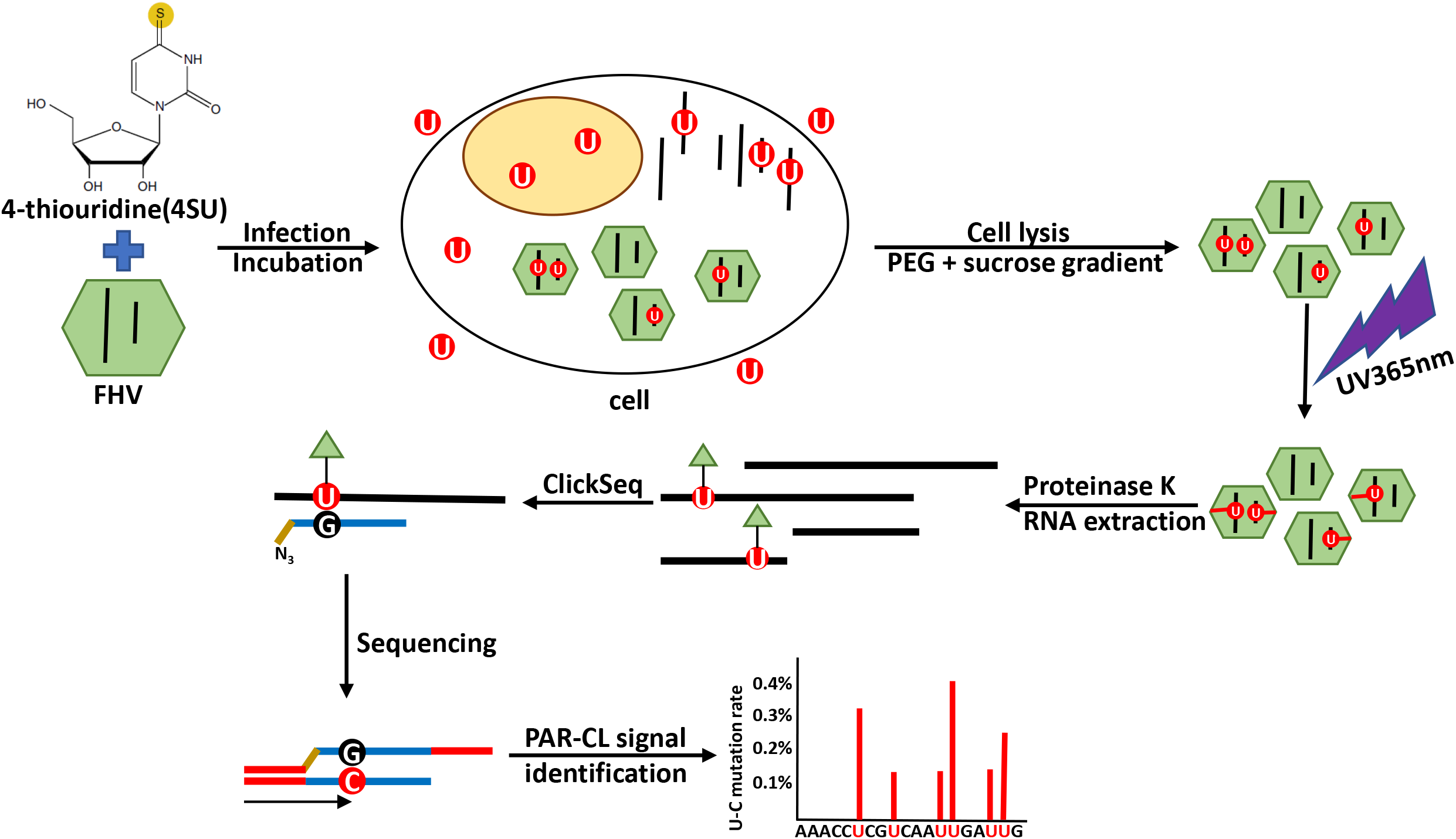
PAR-CL methodology. 4-thiouridine (4SU) were supplemented to S2 cells, during Flock House virus (FHV) infection. After incubation, purified viruses are irradiated with 365nm UV to induce crosslink. After proteinase K digestion, crosslinked RNAs are purified and subjected to ClickSeq with Azido-NTPs. In viral genome, the crosslinked sites are characterized with elevated U to C mutation rates.

The raw sequencing reads were trimmed and quality filtered using *cutadapt* (37) and the *FASTX toolkit* (http://hannonlab.cshl.edu/fastx_toolkit/index.html). Reads were aligned to the FHV genome with *Bowtie (v1.0.1)* (38). The read coverage at each genomic position and the frequency of each nucleotides found in the mapped reads was enumerated using *samtools* and the mutation rates at each genomic position was calculated.

To validate the PAR-CL methodology, we first sought to determine if there was a substantial increase in U-C mutation rate as a specific consequence of 4SU-capsid crosslink. We performed a series experiments in which wild-type (wt) FHV without 4SU (4SU−) or 4SU-containing FHV (4SU+) were treated with (UV+) or without (UV−) UV irradiation. As illustrated for FHV RNA 1 in **Figure 2a**, we plot the measured U-to-C (U-C) mutation rate across the genome and calculate the fold change at each U position. A small number of positions, such as nt. 1259 on RNA 1, showed high U-C mutation rates in both conditions. This possibly reflects the selection of a minority variant during virus passaging. Other than this, we did not notice an increased U-C mutation rate for UV-irradiated FHV in the absence of 4SU (4SU-/UV+). This indicated as expected that UV irradiation alone was not sufficient to induce novel U-C mutations. Similarly, we measured the influence of 4SU substitution in FHV genomic RNA without UV exposure (4SU+/UV−) (**Figure 2b**). We also did not notice an increase in U-C mutation rate. We only observed increased U-C mutations when 4SU and UV irradiation both were present (4SU+/UV+) (**Figure 2c**). This confirms the elevated U-C mutation rate is indeed a specific result of 4SU− induced crosslinking. The FHV RNA 2 data of these experiments is shown in **Supplemental Figure S1**.

**Figure 2.**
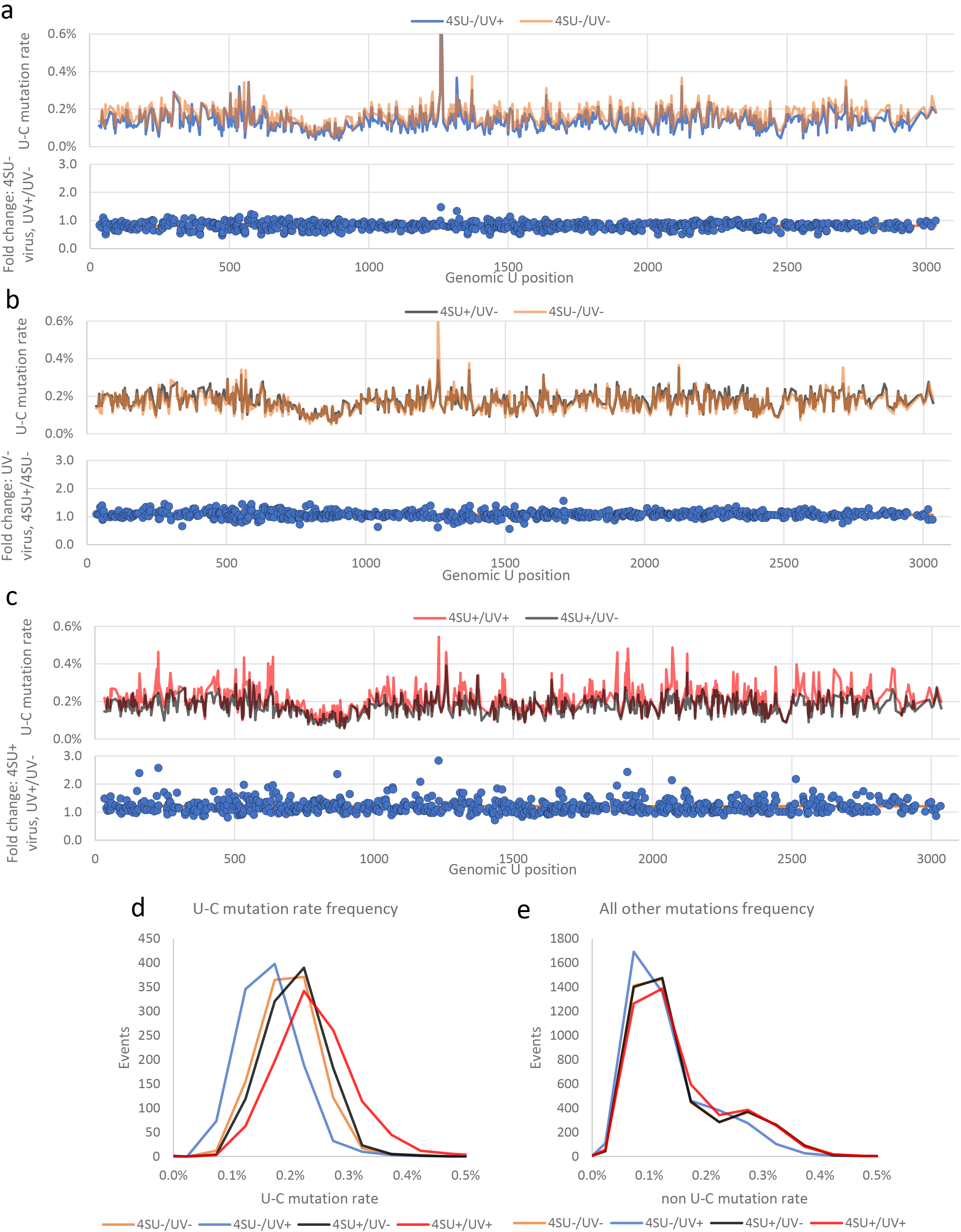
U-C mutation rate elevation is specific to 4SU crosslink. Using FHV RNA 1 as an example, several control experiments were conducted to ensure specificity of PAR-CL signals. **(a)** effects of UV exposure to wt FHV (4SU−) were compared. We did not observe significant U-C mutation rate elevation. **(b)** effects of 4SU incorporation were compared to FHV without UV exposure (UV−). We did not observe significant U-C mutation rate elevation. **(c)** only when 4SU− containing FHV was irradiated with UV, we observed a significant increase in U-C mutation rates. **(a-c)** orange line represents the average fold change in each experiment. **(d)** the distribution of U-C mutation frequency: only when both 4SU and UV both presented, did we notice a shift towards higher U-C mutation rates. **(e)** crosslink did not impact the rates of other mutations (A,G,C-mutations, or U-A/U-G mutations).

Histograms of the U-C mutation rate frequencies at all genomic U positions are shown in **Figure 2d**. This demonstrated that under 4SU+/UV+ condition, more U positions exhibited high U-C mutation rate (≥0.3%) than controls (4SU−/UV− and 4SU+/UV−). Interestingly, we noticed reduced U-C mutation rates when wt FHV was exposed to UV (4SU−/UV+), for an unknown reason. We also sought to determine if 4SU incorporation and UV exposure would induce any non-specific (non-U-C) mutations. A histogram of the frequencies of all non-U-C mutations (A,C,G mutations and U-A, U-G mutations) over all genomic positions is shown in **Figure 2e**. Importantly, 4SU+/UV+ FHV did not show any significant change in non-U-C mutations. We therefore conclude that the increased U-C mutation rate is specific to 4SU-induced crosslinking.

### Magnitude of PAR-CL signals was associated with 4SU dose and incubation time

Our PAR-CL method requires no immunoprecipitation to recover and enrich for crosslinked RNAs. However, this permits wild type uridine and/or uncrosslinked 4-thiouridine to persist in the RNA pool which may dilute the PAR-CL signal. To investigate optimum conditions for PAR-CL, we conducted three parallel experiments (**Figure 3**). S2 cells were infected with FHV and incubated with 4SU at 100μM (4SU16h) or 150μM (4SU1.5X). Viruses were harvested from infected cells 16 h post-infection. In a third experiment, we extended the incubation time to 40 h, and applied 4SU in a “prime-boost” manner, to reach a final concentration of 200μM (4SU40h). In each experiment, U-C mutation profile of 4SU+/UV+ virus was compared with correspondent 4SU+/UV− virus to yield PAR-CL signals (fold change). Results for FHV RNA 1 are shown in **Figure 3a** and RNA 2 in **Supplemental Figure S2**. We noticed that the concentration of 4SU and the incubation time of FHV/4SU had an impact on PAR-CL signal intensities over a number of genomic U positions.

**Figure 3.**
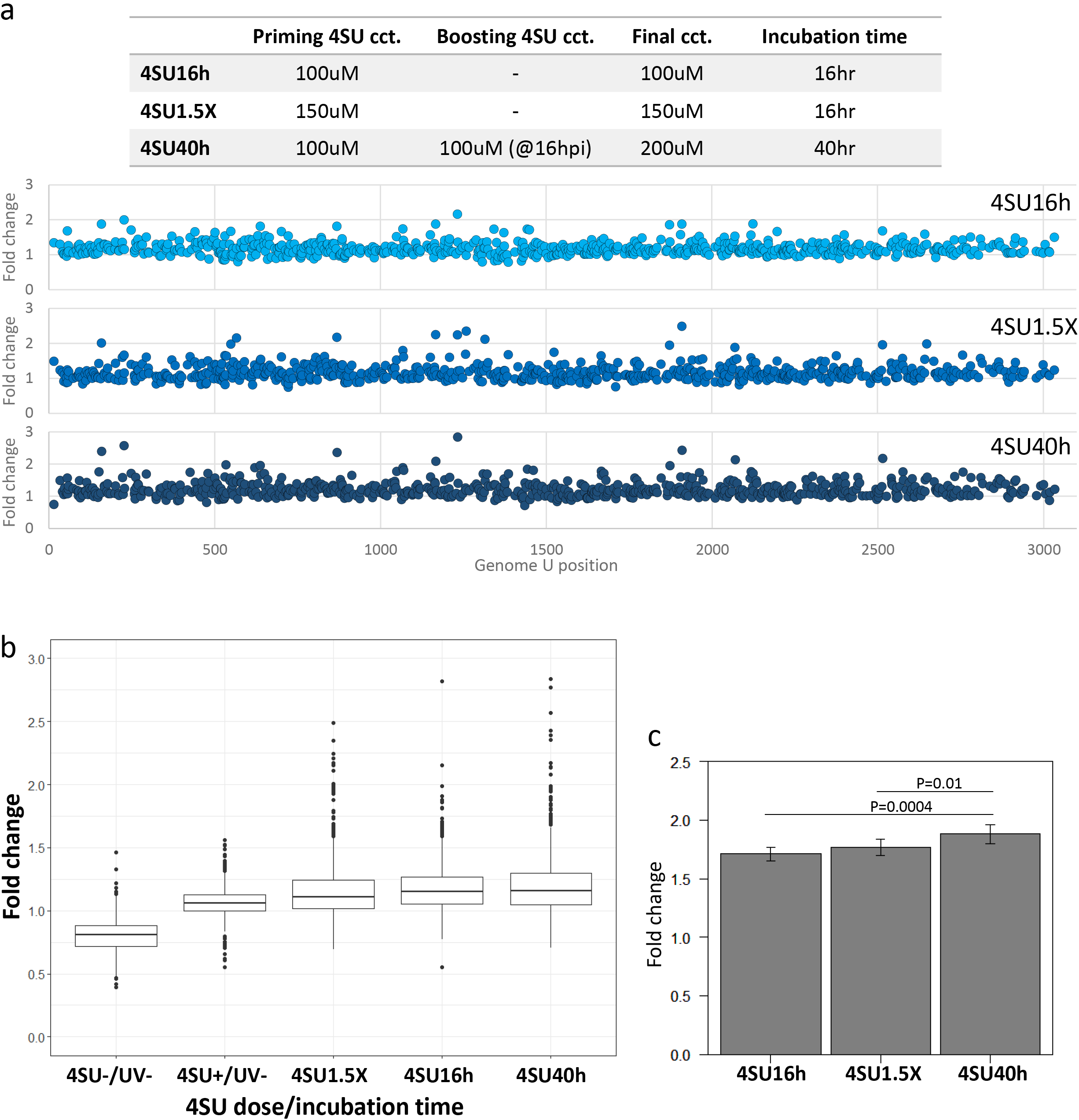
PAR-CL signal intensities correlated with 4SU dose and incubation time. Using FHV RNA 1 as an example, **(a)** three experimental conditions were tested for impact to PAR-CL signals. We observed that the intensities of PAR-CL signals (i.e. fold change of U-C mutation rate) were related to the concentration of 4SU in cell culture and the time of incubation. **(b)** with or without crosslink, the average PAR-CL signals among all 4SU-containing FHVs is similar. However, the outliers of crosslinking groups showed a significant higher PAR-CL signals than that of control (4SU+,UV-), and the magnitude of outliers correlated with 4SU concentration and incubation time. **(c)** We sampled top 5% PAR-CL signals from three experiment groups and determined that optimal PAR-CL signal was achieved under 4SU40h condition, which was applied to all further PAR-CL experiments.

The same results were observed when we plotted the frequency of PAR-CL signals for these three experiments, as well as two controls (**Figure 3b**). This shows that while the mean PAR-CL signals and the distribution under all three conditions (4SU16h, 4SU1.5X, and 4SU40h) and a 4SU+/UV-control were all comparable, the magnitude of outliers showed correlation to experimental conditions (4SU40h > 4SU1.5X > 4SU16h). This indicates that only certain 4SU substitution sites are available for crosslink and therefore sensitive to the varied 4SU concentrations/incubation times. Again, for an unknown reason, the 4SU-, UV+ control (**Figure 3b**) showed a slightly lesser than 1-fold change PAR-CL signal. Importantly, this does not interfere with our interpretation of PAR-CL signals in crosslinking samples.

We sampled the top 5% of PAR-CL signals in each conditioned experiment (**Figure 3c**) and concluded that, the 4SU40h group showed significantly higher PAR-CL signals than the rest. For this reason, the 4SU40h experimental condition was applied to all further PAR-CL experiments, unless otherwise mentioned. Despite the varied PAR-CL signal intensities under different experimental conditions, the PAR-CL signals presented good Pearson’s correlation coefficient (≥0.6) between these parallel experiments (**Supplemental Figure S3**), which indicates high reproducibility of PAR-CL experiments.

### Consistent PAR-CL signals indicates structural tropism of encapsidated viral RNAs

We applied the 4SU40h condition in three parallel experiments: three separate S2 cell cultures were incubated with virus and 4SU, and individually purified viruses were exposed to UV and thereafter proceeded to sequencing. Similar to before, PAR-CL experiments were conducted in pairs, with each PAR-CL dataset comprised of one crosslinked sample (4SU+/UV+) and control with the same sample but without UV irradiation (4SU+/UV−). To ensure reliable mutation rate calculation, we selected for U positions with coverage of at least 10,000 reads. This allowed us to detect reliable mutation profiles over U34 – U3034 on RNA 1, and U9 – U1337 on RNA 2. The PAR-CL signals of these three experiments were compared on each U position on viral RNA genomes (**Figure 4a, b**). We observed good Pearson’s correlation coefficient (≥0.6) between these replicates (**Supplemental Figure S4**). To validate the consistent PAR-CL signals, the signals over every U position were box-plotted over the triplicates (**Supplemental Figure S5**). This allows us to readily measure the mean signal strength and signal variation over the triplicates. In order to distinguish reliable crosslinking sites and avoid potential false positives, we removed any PAR-CL signals in crosslinked sample by applying a conservative background threshold filter, retaining only the highest 5% of PAR-CL signals in the uncrosslinked control sample. (**Figure 4c, d**, **Supplemental data 3**). Among the most consistent PAR-CL sites (passed background threshold in all three replicates), we identified 20 sites in RNA 1 and 8 sites in RNA 2 that showed the highest average PAR-CL signals (**Figure 4e**, **Supplemental data 3**). T-test revealed most of these sites have significantly (P<0.05) higher PAR-CL signal than average. As these same sites consistently displayed significant PAR-CL signals over parallel replicates, this indicates a set of consistent RNA-capsid interactions in authentic FHV virions, which further indicates a structural tropism of FHV RNA in association with the topology of virus capsid shell.

**Figure 4.**
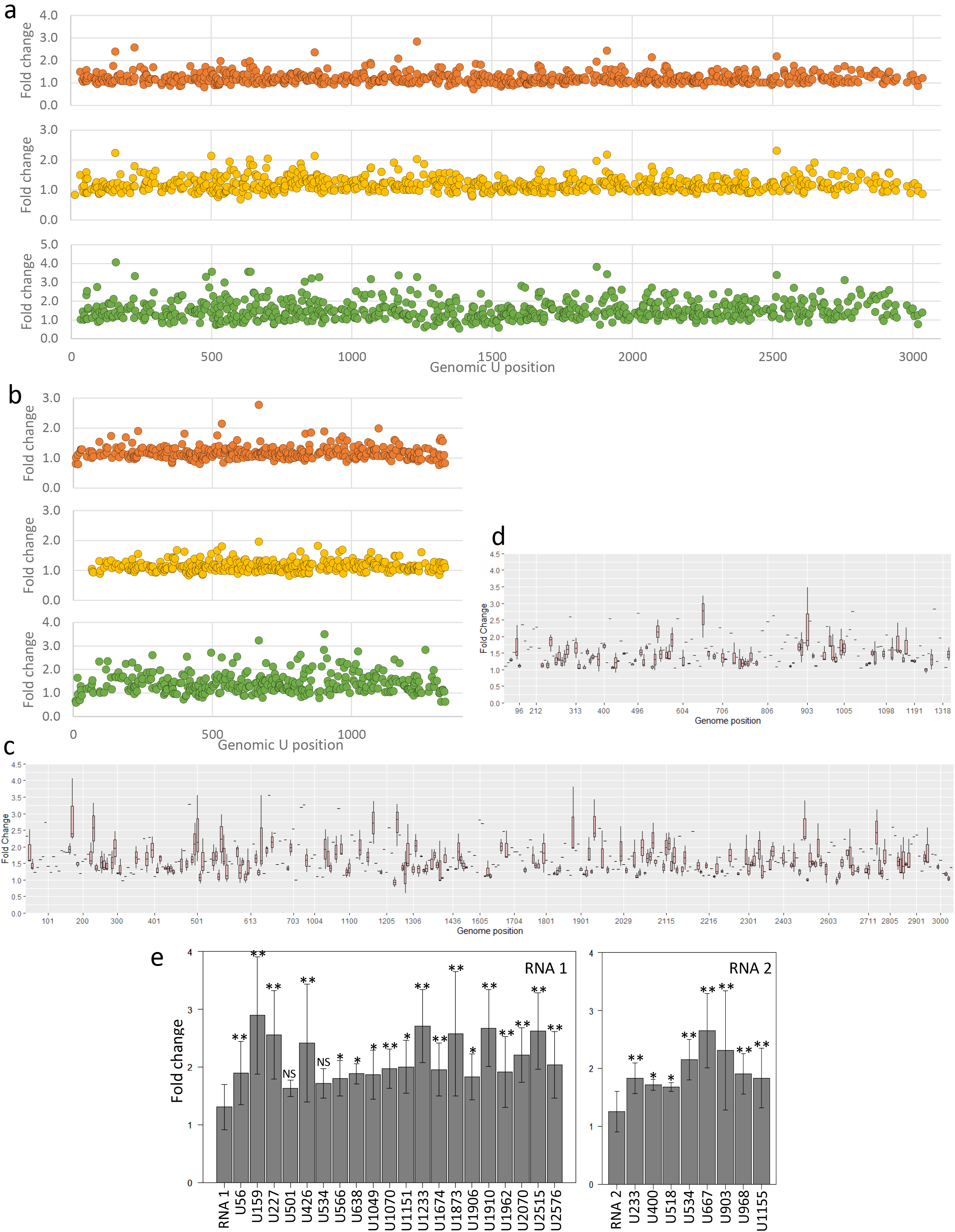
Consistent PAR-CL sites revealed clear RNA tropism. **(a, b)** PAR-CL signals of FHV RNA 1 and 2 respectively, in triplicates. **(c, d)** Triplicated PAR-CL signals of FHV RNA 1 and 2 were box-plotted. We removed any PAR-CL signal failed to pass the background threshold. A number of sites on both RNA 1 and RNA 2 showed consistently significant PAR-CL signals, indicating reliable crosslinking sites between RNA and protein. These consistent PAR-CL sites also suggest a strong tropism of FHV RNA cage inside virion. X-axis is not continuous. **(e)** Among these consistent PAR-CL sites, most of them showed significantly higher PAR-CL signals than the average. *P<0.05; **P<0.01; NS=not significant.

### Probing FHV in virion RNA secondary structures with DMS-MaPseq

We sought to understand if there is any sequence motif among the PAR-CL sites. Significant (>2σ) PAR-CL sites (28 sites from RNA 1 and 15 from RNA 2) and their flanking sequences were analyzed with Discriminative Regular Expression Motif Elicitation (DREME, (39)) for possible sequence motif identification (**Supplemental Figure S6**). However, no common motif was identified. This led us to hypothesize that the mechanism of RNA recognition by FHV capsids may be related to similar RNA structures rather than sequences. To reliably predict the RNA secondary structures of PAR-CL signals, we sought to determine the secondary structure of FHV RNA in authentic virions.

Dimethyl sulfate mutational profiling with sequencing (DMS-MaPseq) provides a reliable and high throughput method to probe RNA secondary structures *in vivo* (40–42). The resulting constraints provided improvement to thermodynamic map and free energy-based secondary structure prediction. We performed DMS-MaPseq using the TGIRT^TM^-III enzyme but in combination with ClickSeq to generate RNAseq libraries (“TGIRT-ClickSeq”) (**Figure 5a**), demonstrating that TGIRT^TM^-III enzyme is compatible with ClickSeq. DMS-MaPseq induces RNA modifications to unpaired adenines and cysteines (and guanine to a lesser level (43)) across the viral genome. Therefore, as expected, in comparison to untreated control virus (DMS−) DMS-treated FHV (DMS+) has a higher average mutation rate over genomic A/C positions (**Figure 5b**). Similarly, we plotted the frequency of mutation rates over A/C or G/U positions and only noticed a significant higher mutation rate frequency over A/C positions (**Figure 5c**). We analyzed A or C positions with at least 10k read coverage, which corresponds to nt. 14 -3043 on RNA 1 and nt. 11 - 1378 on RNA 2. Similar to PAR-CL data, DMS-MaPseq signal represents the mutation rate fold change between DMS-treated virus and untreated control virus, on all genomic A or C positions. Likewise, we removed potential false-positive signals by applying a background noise threshold, retaining only the genomic sites with mutation rate higher than this threshold. The resulting DMS-MaPseq profile of FHV (**Figure 5d** and **e**) showed clear signals up to 100-fold change over both RNA 1 and 2. The un-refined DMS-MaPseq profiles with background noise, and mutation rate comparison between DMS-treated and untreated viruses are shown in **Supplemental Figure S7**.

**Figure 5.**
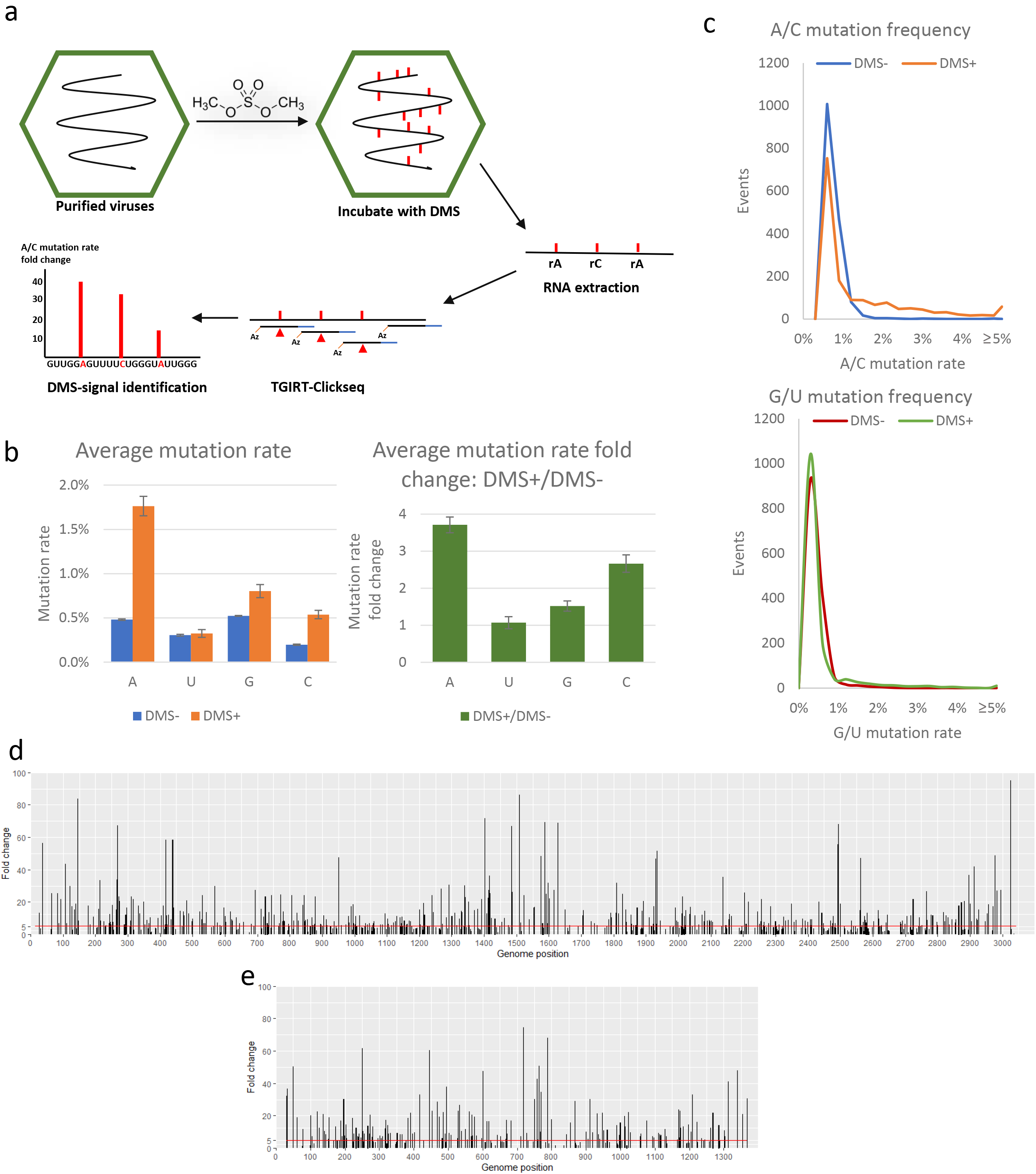
FHV DMS-MaPseq. **(a)** In order to precisely predict the secondary structure of FHV RNA, we used DMS (dimethyl sulfide) to induce in virion methylation of unpaired adenines and cystines. The extracted RNAs were subjected to ClickSeq library preparation with TGIRTTM-III enzyme, which invokes mutations over methylated bases. DMS-MaPseq signal represents the mutation rate fold change over A/C positions. Red markers on RNA represent methylated ribonucleotides; red triangles on cDNA represent DMS-induced mutations. **(b)** of A/C mutations were detected as a result of DMS treatment. **(c)** DMS treated virus exhibited higher mutation rate frequencies over A/C positions. **(e,f)** DMS-MaPseq map of FHV RNA 1 and RNA 2, respectively. Background noise was removed. Red line represents the average DMS-MaPseq signal.

### DMS-MaPseq resolved FHV RNA secondary structures reveals that PAR-CL sites favor double stranded structures and are highly clustered

We incorporated the DMS-MaPseq data into free energy based thermodynamic prediction, by introducing a series of “soft” constraints. Only the most significant (>2σ) DMS-MaPseq sites (60 sites in RNA 1 and 30 sites in RNA 2) were forced as unpaired constraints in RNAstructure Web Server(44) with “Fold” algorithm (44, 45). Regardless of their DMS-MaPseq signals, the remaining genome positions were left without any constraints, to allow maximum prediction flexibility. We thereby constructed a DMS-MaPseq-imposed secondary structure map of complete FHV RNA genome (snapshots in **Figure 6**, full-scaled maps of RNA 1 and RNA 2 were also provided in **Supplemental data 1** and **2**). Despite the low number of introduced constraints, we were able to greatly improve the thermodynamic mapping of FHV RNAs. With the 60 RNA 1 constraints, 37% (1145/3107) of nucleotides underwent refolding compared to the unconstrained model, yielding different paired/unpaired patterns. Similarly, with the 30 RNA 2 constraints, 20% (273/1400) nucleotides underwent refolding. The dot-bracket maps comparing the differences between unconstrained and constrained folding can be found in **Supplemental Figure S8**.

**Figure 6.**
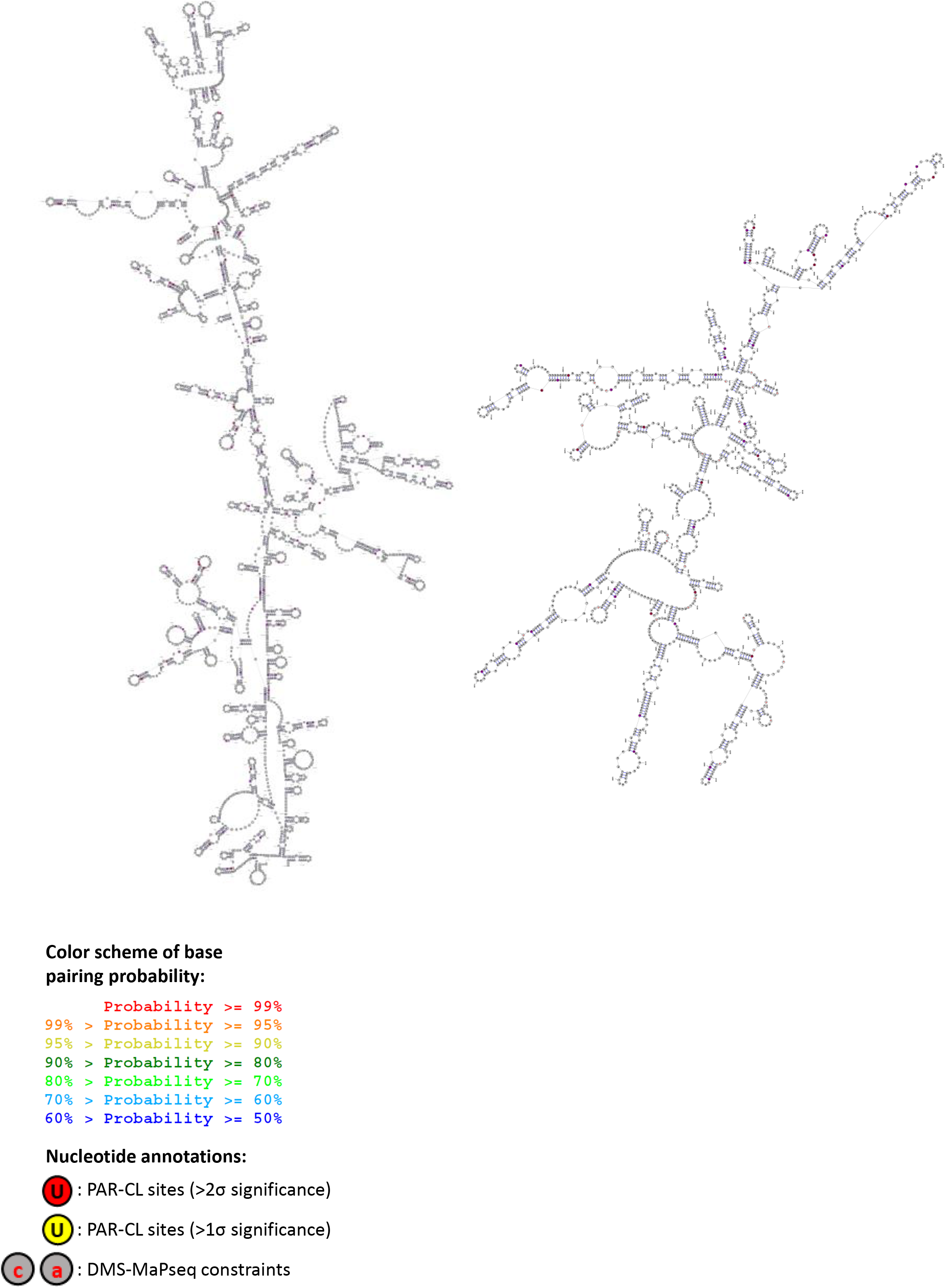
DMS-MaPseq-corrected secondary structure map of complete FHV RNA genome. Snapshots of RNA 1 (left) and RNA 2 (right) are shown. Full scaled maps can be found in Supplementary data 1 and 2. PAR-CL signal sites of different significance were color annotated. The introduced DMS-MaPseq constraints were highlighted by lower case “a” or “c” in red color. The base pairing probabilities of representative PAR-CL sites are color schemed on the base pairing bonds. Some examples are shown in **Table 1**.

In combination with PAR-CL data, we noticed that the significant PAR-CL sites heavily favored double-stranded base-pairing. In RNA 1, among the 28 most significant (>2σ) PAR-CL sites, 22 are located in double-stranded regions, whereas 3 sites were 1 nt. adjacent to double-stranded stems. In the much shorter RNA 2, 8/15 of most significant PAR-CL sites are located in dsRNA stems, whereas 3 are 1 nt. adjacent. In **Table 1**, we illustrate the detailed structures of 16 PAR-CL sites (11 on RNA 1 and 5 on RNA 2) that presented with highest consistency and average PAR-CL signals (**Figure 4a-d**).

**Table 1.**
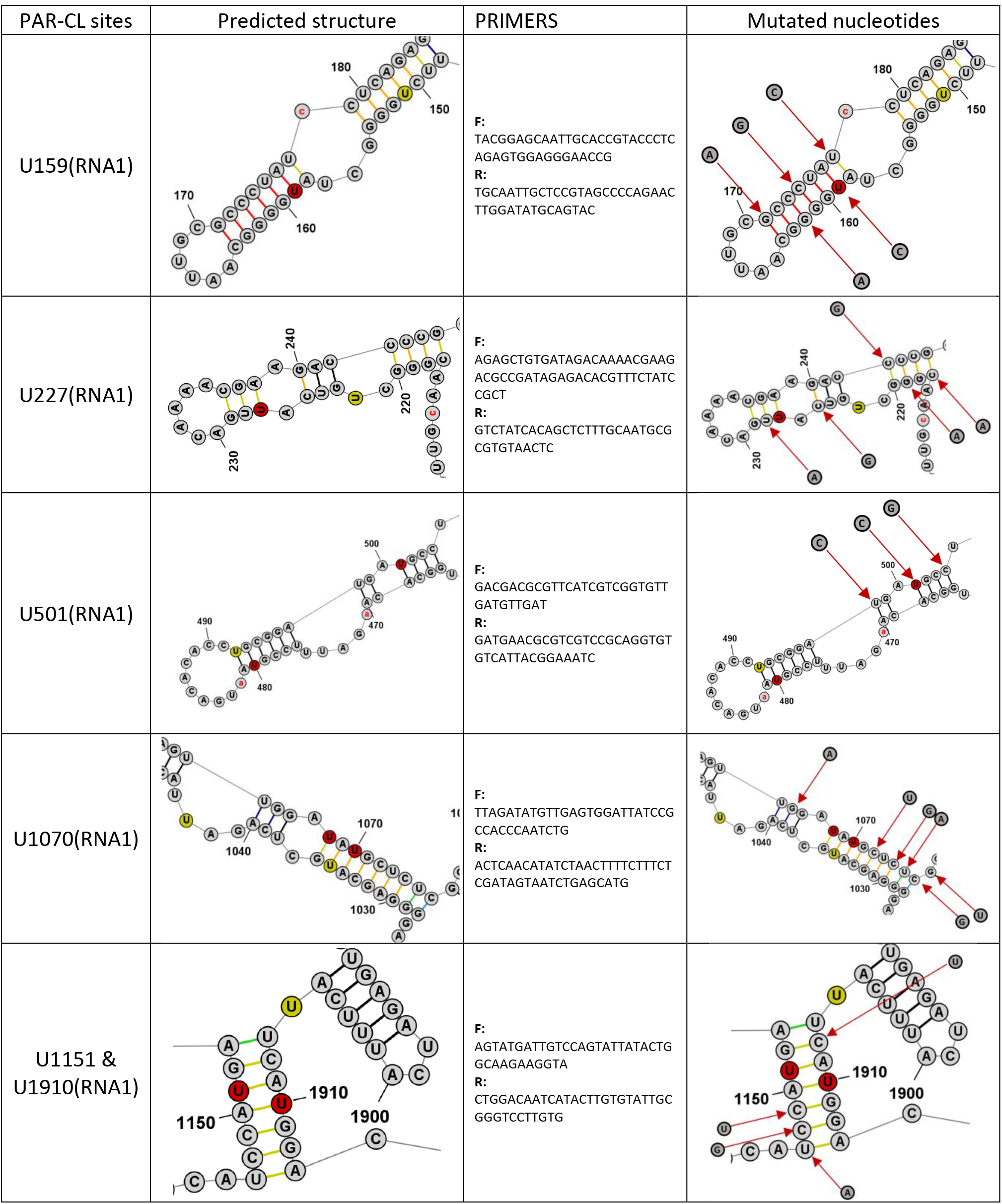
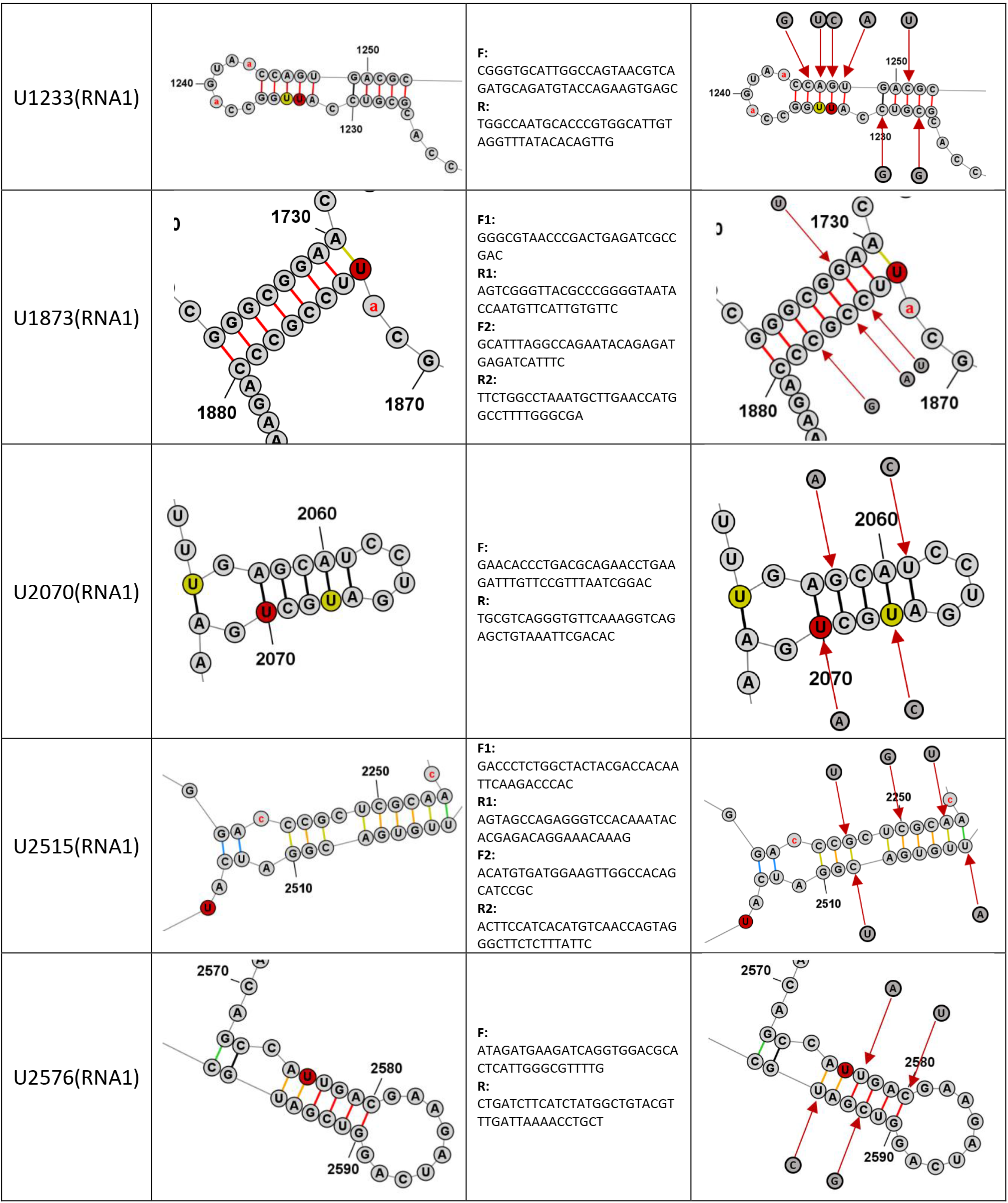
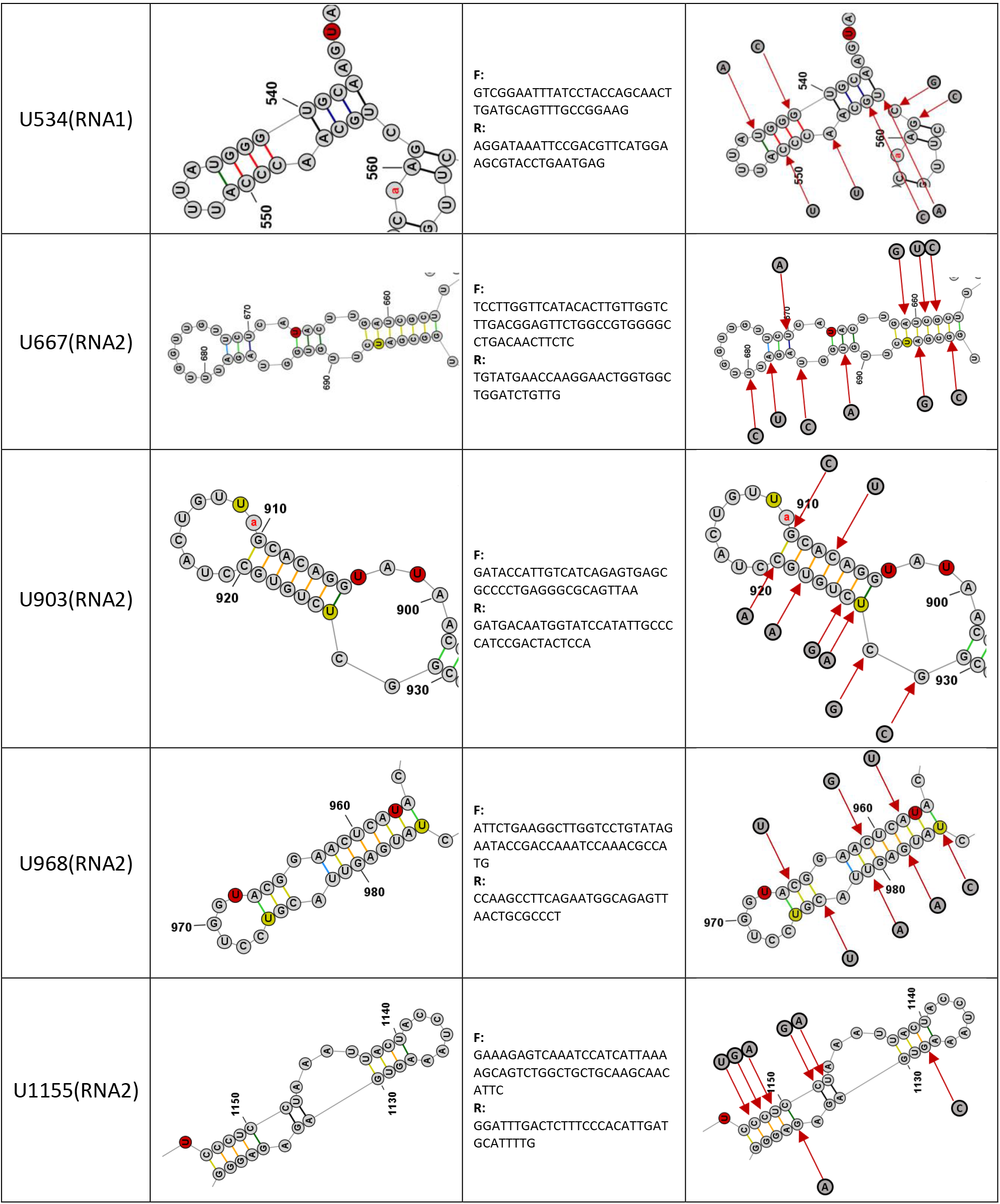
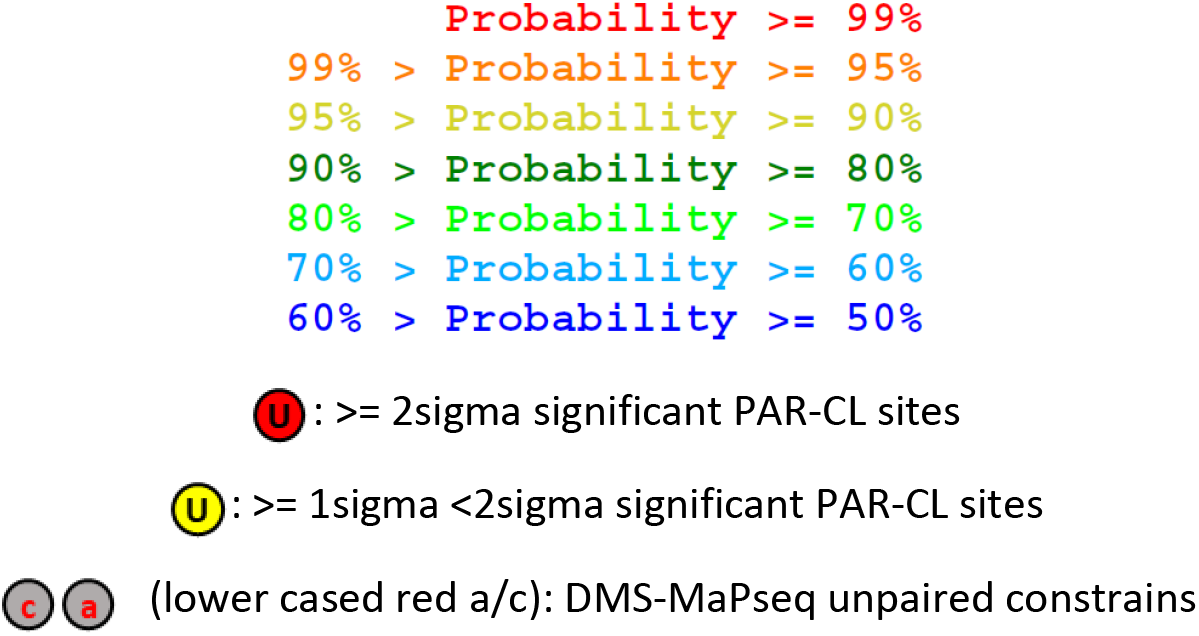

We also noticed that the distribution of PAR-CL signals was uneven and highly clustered. Numerous PAR-CL stems showed more than one PAR-CL sites with >1σ significance (**Supplemental data 1, 2**, some examples were listed in **Table 1**). We calculated the average shortest distance between adjacent PAR-CL sites. On RNA 1 (3107 nt.), among 721 uridine sites, 102 showed >1σ significant PAR-CL signal. The average shortest distance between these PAR-CL sites is 7.4 nucleotides, which is substantially shorter than the average shortest distance of 102 random uridines (30.47 nucleotides). On RNA 2 (1400 nt. genome with 351 uridines), the average shortest distance among 45 PAR-CL sites (>1σ significance) was 8.8 nt., which is also shorter than the average shortest distance of 45 random uridines (31.11 nt.).

Notably, by combining PAR-CL data and DMS-MaPseq-imposed RNA structure, we are able to characterize a stem loop site which is structurally near identical to a previously predicted stem loop (nts. 168-249) on RNA 2 (24) (**Supplemental Figure S9**). This stem loop site, as well as the flanking sequence (nt. 210-249) has been determined to be essential for RNA 2 encapsidation. We identified three PAR-CL signals within this region, consistent with role of this stem loop site in RNA 2 packaging.

### Structurally-disrupted PAR-CL sites impact FHV lifecycle and fitness

To determine whether the identified PAR-CL sites have a biological function, we selected 11 PAR-CL sites from RNA 1 and 5 PAR-CL sites from RNA 2 as our candidate sites (**Table 1**). Referring to the DMS-MaPseq-corrected FHV RNA structure maps (**Figure 6**), we introduced synonymous mutations to disrupt the double-stranded RNA regions of the PAR-CL sites (or the nearest stem of certain PAR-CL sites, i.e. U2515 on RNA1, U534, U903, U968, and U1155 on RNA2). The predicted structure of these PAR-CL sites, primers, and replaced nucleotides are listed in **Table 1**. Plasmids containing these point mutations were transfected into S2 cells. Each transfection consisted either a mutated RNA1 and wild-type RNA2, or a mutated RNA2 and wild-type RNA1 (**Figure 7a**). After transfection, induction and incubation, cell viability of each transfected mutant was determined with alamarBlue assay (**Figure 7b**). Almost all mutant virus transfections showed reduced cytopathic effect compared to transfection with wild-type FHV RNA. Notably RNA 1 mutants U159, and U1233 resulted in little to no detrimental effect to S2 cells.

**Figure 7.**
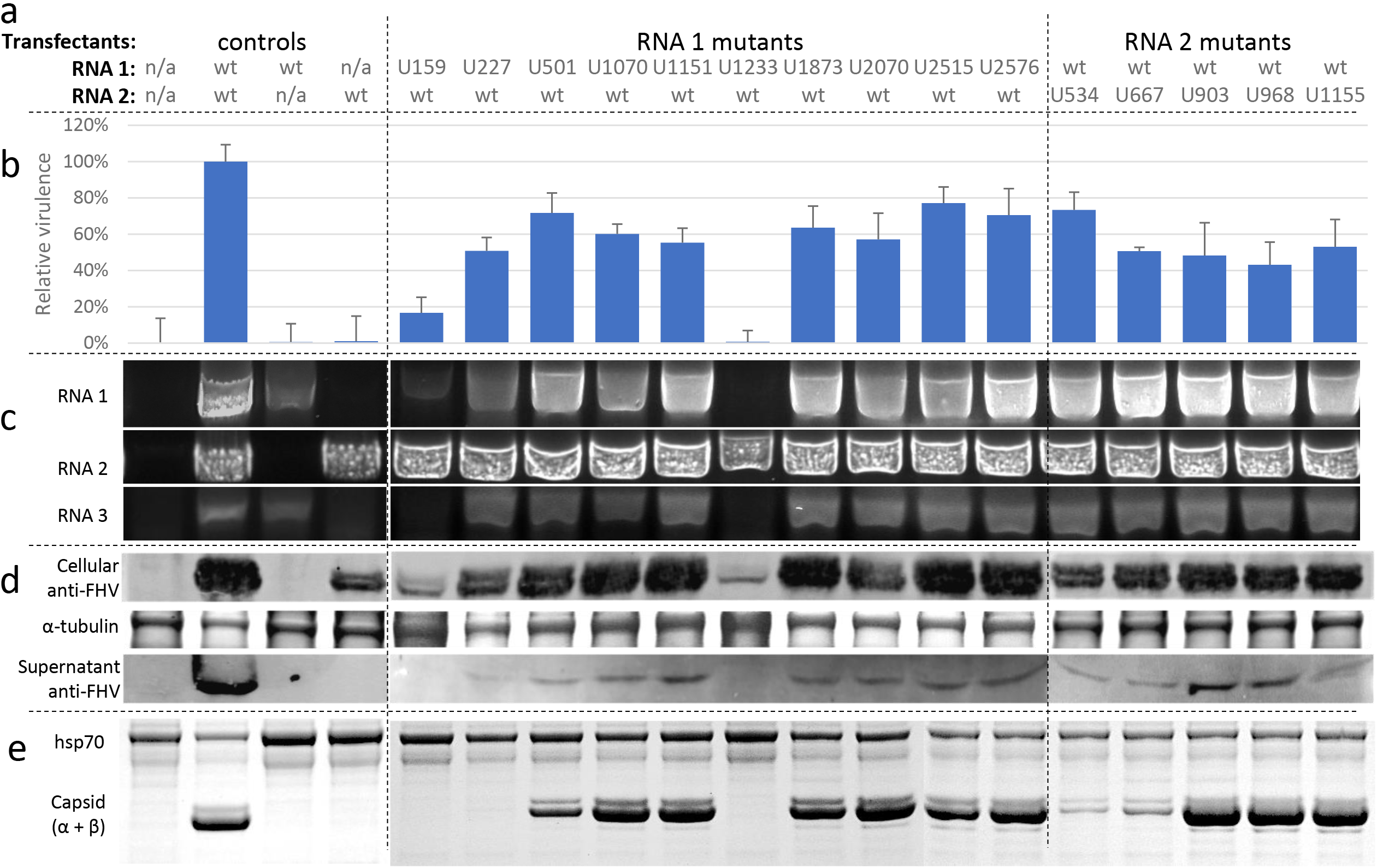
FHV PAR-CL mutants characterization. **(a)** Plasmids containing FHV RNA 1 or RNA 2 with mutated PAR-CL structures were transfected to S2 cells to yield p0 mutant viruses. **(b)** Relative virulence of p0 mutant viruses was determined with alamarBlue assay to measure cell viability after transfection. **(c)** 200ng of total cellular RNA of each transfection was analyzed with RT-PCR to measure the accumulation of FHV RNAs. **(d)** Cellular and supernatant FHV capsid productions were detected with anti-FHV antibodies. Coomassie-stained α-tubulin as loading control for cellular assay. **(e)** p1 viruses were purified and filtered with 100 K molecular weight filter, mutant virus production was verified with SDS-PAGE gel. Heat shock protein 70 (hsp70) shown as a loading control.

Total cellular RNA was extracted from transfected cells and in-column DNase digestion was conducted to remove remaining plasmids. From each transfection, 200ng of purified RNA was used as template for RT-PCR to detect FHV RNA (**Figure 7c**). We noticed that accumulation of FHV RNA 2 was unaffected by any PAR-CL mutants, while RNA 1 accumulation varied drastically among RNA 1 mutants. Notably, RNA 1 mutant U1233 yielded undetectable levels of RNA 1 and RNA 3, while RNA 2 production was less affected. RNA 1 mutant U159 also produced marginal amount of RNA 1 and RNA 3, and U227 produced substantially less RNA 1 than that of control or RNA 2 mutants. The replication deficiency of these three mutants agreed with our findings of their low virulence (**Figure 7b**). Interestingly, these three sites are found within or adjacent to previously described FHV RNA regulatory regions (46, 47) **(Figure 8)**. The importance of these three sites in both RNA-capsid interaction shown here and RNA replication regulation indicates that the same motifs in the RNA genome are involved in multiple stages of the viral life-cycle, consistent with the notion that replication and RNA genome packaging are tightly coupled processes (20, 21).

**Figure 8.**
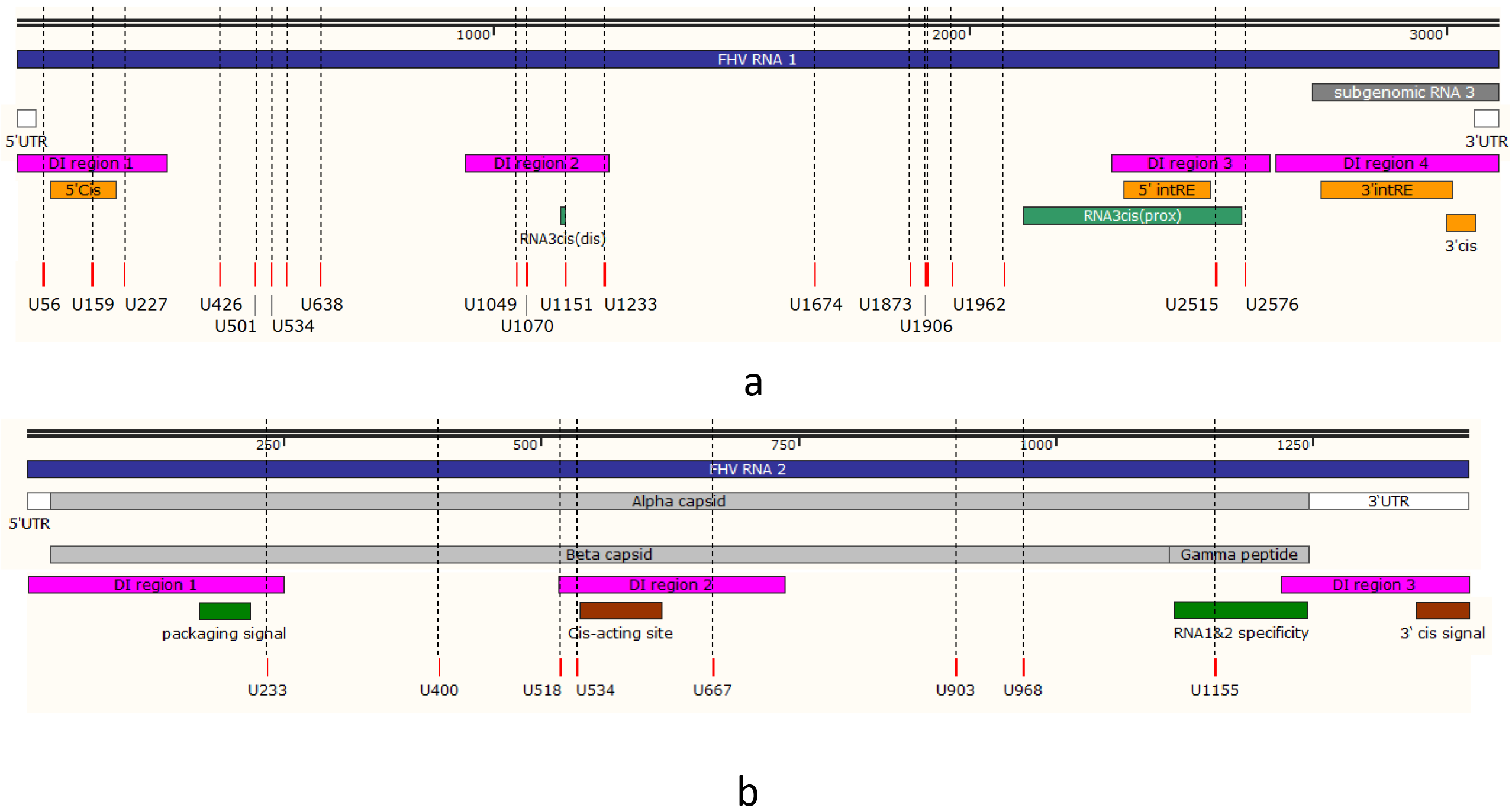
FHV significant PAR-CL signals and known motifs. **(a)** On RNA 1, four highly conserved defective interfering RNA regions (DI regions 1-4)(34) were shown in pink; four RNA 1 replication regulatory elements were shown in orange (5’Cis (46), 5’intRE(47), 3’intRE(47), and (67)); two RNA 3 regulatory elements and a putative RNA 3 subgenomic promoter region were shown in green (RNA3cis(dis) (47), RNA3cis(prox)(47)); the candidate PAR-CL sites (listed in Figure 4e) were shown as red bars on bottom. **(b)** On RNA 2, three conserved defection interfering RNA regions were shown in pink (DI regions 1-3 (34)); a mid-genome cis-acting replicational element(59) and a 3’ cis-acting regulatory element(67) were shown in brown; a RNA 2 packaging signal (24) and a capsid site essential for RNA 1&2 specificity (18) are shown in green.

To confirm capsid production, we separated cells and supernatant from the transfected cells. Western blots with anti-FHV were used to detect capsid proteins in both cellular components and supernatants (**Figure 7d**). In the cellular fraction, we readily detected both mature (alpha peptide) and autoproteolytically cleaved capsid protein (beta peptide) in all mutants. However, reduced capsid yields were found in U159 and U1233 mutants, possibly due to the observed RNA 1 replication deficiency. In supernatant fractions, the U159 and U1233 mutants resulted in undetectable level of capsid protein, while U227 resulted in detectable but very marginal amount of capsid production. This confirmed that the mutations at these three PAR-CL sites have significant impact on virus production in S2 cells.

To expand mutant viruses, we further inoculated naïve S2 cells with equal amount of transfected p0 cell mix. From the inoculated P1 cell culture, we observed different degrees of cytopathic effect (CPE) under microscope (**Supplemental Figure S10**), which was correlated to earlier findings. P1 mutant viruses were nuclease treated, PEG precipitated, and purified with PES membrane protein concentrator to remove potentially unassembled capsid subunits. The presence of virus particles was confirmed with SDS-PAGE (**Figure 7e**), and virus yield was calculated by densitometry. Similar to before, we failed to detect virus production of U159 and U1233 mutants, while U227 mutant resulted a marginal virus production which can only be detected by western blot but not with SDS-PAGE. This result also agreed with our western blot analysis (**Figure 7d**).

We further tested P1 mutant virus relative virulence by infecting cells with mutants at MOI = 1 (**Supplemental Figure S11**). Most mutant viruses still resulted in varied but inferior virulence, in comparison to wild type virus.

## Discussion

In this study, we demonstrated that PAR-CL can be used as a reliable and convenient method to screen for capsid-interacting sites on viral RNA genomes. PAR-CL data analysis features low background noise and thus, highly distinguishable signals. Therefore, PAR-CL signals are highly specific and representative of consistent crosslinking events between virus RNA and capsid. We showed that under the optimized experimental condition (4SU40h), FHV PAR-CL replicates revealed consistent crosslinking sites, indicating clear structural tropism of the asymmetrically packaged genomic RNA inside particles.

We also demonstrated that by combining a functional screening method PAR-CL and a structural probing method DMS-MaPseq, we can relate virus RNA-capsid interactions with RNA secondary structures, and *vice versa*. DMS-MaPseq was used to construct the whole genome RNA secondary structure maps of FHV, with which we observed that PAR-CL signals were highly clustered and favored double-stranded RNA stems. Synonymous mutations were designed to disrupt the double stranded structures in candidate PAR-CL sites. These structural mutants resulted in varied effects with most displaying reduced cytotoxic effect after transfection, while certain mutants were detrimental to RNA 1 replication. Together, we demonstrated that in combination with DMS-MaPseq, PAR-CL method can be used *de novo* to identify RNA regions that are important in virus packaging, virulence, and/or replication.

### PAR-CL methodology

Photoactivatable nucleoside analogs were successfully utilized in the past to enhance crosslinking efficiency and hence, providing approaches to study RNA-RNA and RNA-protein interactions (reviewed in (48)). Thionucleobases such as 4-thiouracil (4SU) and 6-thioguanosine (6SG)) allows for highly effective crosslinking at 330-365 nm excitation spectrum (49), as well as advantages such as minimum nucleoside structure perturbation (48, 50), lower cytotoxicity (26, 32, 33, 51), and less photochemistry and/or photodamage (48, 50). Importantly, the 4SU/6SG incorporated RNA can lead to specific base mismatches during reverse transcription (U-C, and G-T)(27–29), which enables high-throughput screening as indications of crosslinking. This is best illustrated with PAR-CLIP (PhotoActivatable-Ribonucleoside-enhanced CrossLinking and ImmunoPrecipitation) technology (26), which allows for pinpointing crosslinking sites at nucleotide resolution. PAR-CLIP has been successfully applied in the past to identify crosslinking sites of Argonaute 2, embryonic lethal abnormal vision (ELAV) protein and pumilio homologue 2 (PUM2), insulin-like growth factor proteins (26, 52).

As its primary purpose, PAR-CLIP was designed to screen entire transcriptome for RNA sequences binding to RBP-of-interest. Typically (26, 30, 31), PAR-CLIP was conducted by incubating cell cultures with 4SU, followed by 365 nm UVA irradiation, cell lysis, RNase T1 digestion, immunoprecipitation of RBP-of-interest, second RNase T1 digestion, de-phosphorylation, radiolabeling, SDS-PAGE and electro-elution, proteinase K digestion, and RNA extraction. The recovered crosslinked RNA then is used as a template for cDNA library preparation and deep sequencing. A natural prerequisite is large amounts of starting materials (between 100-400 × 10^6^ cells (30)).

The unique aspect of our simplified PAR-CL (PhotoActivatable-Ribonucleoside-enhanced CrossLinking) method is that we applied the similar PAR-CLIP principles to an RNA virus (FHV), which can be easily separated from cellular components. Crosslinks within purified virus particles allow us to: (1) eliminate the need for immunoprecipitation to recover crosslinked RNA; (2) look for specific in virion interactions between viral RNA genomes and viral capsid proteins; (3) study a reductionist and highly controlled microenvironment. The greatly simplified PAR-CL methodology, in combination with ClickSeq library construction technology (36), granted the ability to conduct an experiment with as little as 2 μg of purified FHV particles. A single T25 flask of S2 cells can generate ample amount of pure 4SU-containing viruses to conduct multiple PAR-CL experiments.

In our PAR-CL method, the final pool of purified viral RNA can comprise large number of wild type uridines, or uncrosslinked 4SUs. As a consequence, the signal of any randomly generated, non-specific crosslinking event will be largely diluted into background level. Only the consistent crosslinking sites present due to homogeneity in RNA-capsid interactions within a viral population can readily provide distinguishable PAR-CL signals from background. Therefore, in contrast to the canonical PAR-CLIP approach where only cross-linked RNA fragments are sequenced, we are also able to identify regions of the viral genome where there is no reproducible PAR-CL signal, either due to a lack of RNA-capsid interactions or heterogeneous interactions. This is best illustrated in **Figure 3b**, where the background noise levels are largely unchanged, with or without crosslinking.

In both PAR-CLIP and PAR-CL, there are intrinsic limitations of 4SU-induced crosslinking. Firstly, crosslinking is only limited to U positions. Any potential interaction between protein and other nucleotides is undiscoverable. Next, 4SU crosslinking with protein is affected by reactivity of amino acid side chains (27, 29), with aromatic amino acids (phenylalanine, tyrosine, and tryptophan) being predominant targets but also lysine and cysteine (27). Consequentially, not all RNA-protein interactions can be depicted by PAR-CL or PAR-CLIP, and certain interactions may not result in crosslinking.

### FHV PAR-CL experiments and data analysis

Several approaches were used to ensure reliable interpretation of PAR-CL signals on FHV: 1) to ensure reliable interpretation of mutation rate, we limited our analysis to genomic positions with at least 10k coverage. For this reason, our FHV PAR-CL experiments reliably covered U34 – U3034 on RNA 1, and U9 – U1337 on RNA 2. However, it is possible that we omitted potential capsid interaction sites out of our analyzed range. 2) We previously noticed that certain point mutations may be selected by virus and could be associated with defective interfering RNA generation (34). In this study, we also noticed substantially increased mutation rates on certain genomic positions (such as U1259 on RNA1, as illustrated in **Figure 2a-c**). Thus, to eliminate virus intrinsic mutational events, we avoided to use U-C mutation rate as a measurement. Instead, we decided to use fold change of U-C mutation rate, between crosslinked virus and uncrosslinked virus control, as our PAR-CL signals. 3) Because our PAR-CL signal corresponds to the fold change of U-C mutation rates of two datasets, a substantial PAR-CL signal can be a consequence of three scenarios: a high U-C rate in crosslinked virus, a much lower U-C rate in uncrosslinked control, or both. To minimize the possibility of false positives, we introduced a background threshold. Only the PAR-CL signals above this threshold were taken into our further consideration, as they represent mutation rates distinguishable from background fluctuation range (illustrated in **Supplemental data 3**). Together, we believe these three quality control measurements provided stringent analysis to our PAR-CL data to reveal truly biologically relevant FHV RNA-capsid interaction sites.

### DMS-MaPseq and FHV secondary structure mappings

Several studies have proposed lowest free energy-based FHV local or whole genome secondary structure predictions, with the focus on viral RNA intracellular arrangement and replication regulations (24, 46, 47, 53). *In vivo* RNA chemical probing methods such as DMS and SHAPE allow for structure-specific chemical modifications to be screened by next generation sequencing techniques (40, 54, 55). Using DMS-MaPseq in authentic FHV virions, we are able to provide experimental validation of the RNA structures inside virus particles. With the same rationale of PAR-CL, we also applied stringent quality control measurements to ensure reliable interpretation of mutational profiles generated by DMS-MaPseq: A/C error rates were only analyzed over positions with at least 10k coverage (A14-A3043 on RNA 1, and C11-A1378 on RNA 2); fold change of A/C mutation rate was regarded as DMS-MaPseq signals instead of actual mutation rate; similar background noise threshold was also applied to prevent potential false positives. Canonically, DMS-MaPseq data is imposed upon thermodynamic prediction by enforcing unpaired constraints on any position with a signal above a given threshold (40). In this study, we adjusted this approach by only allowing the most significant (top 5%) DMS-MaPseq signals to be unpaired constraints. However, in this study, we constructed FHV secondary structural maps over RNA 1 and RNA 2 separately, omitting potential inter-RNA interactions.

Combining PAR-CL and DMS-MaPseq, we demonstrated that these two high-throughput mutational profile technologies can work synergistically to answer basic virology questions. We observed that the FHV RNA-capsid sites heavily favored double stranded RNA structures. This finding agreed with earlier predictions that the RNA duplexes scaffold the 2-fold axis of FHV capsid (12).

### Flock house virus PAR-CL sites and biological indications

It has been observed previously that the RNAs of FHV, as well as other Nodaviruses, form a highly ordered dodecahedral cage inside virus particles (10, 56). However, it was not clear whether the dodecahedral RNA cage had a fixed topology. From our PAR-CL data (**Figure 4a-d**), we can clearly identify highly consistent RNA-capsid interactions over certain genomic positions among multiple replicate experiments. This provides evidence that there is well-defined tropism between FHV RNA cage and capsid shell, at least at a these sites identified here. Among the most consistent and distinguished PAR-CL sites **(Figure 4c, d)**, we noticed that they exhibited a highly clustered pattern. The clustering effect is more pronounced, when taking RNA secondary structures into consideration (**Figure 6** and **Supplemental data 1, 2**).

The multiple RNA-capsid interaction sites spanning the whole FHV genome suggest the possibility that FHV encapsidation may require multiple packaging signals to assembly the entire virus genome. Flock House virus genome packaging process may be similar to the two-staged packaging mechanism of MS2 bacteriophage (57). Numerous synergetic high-affinity RNA-capsid interaction sites are required to recruit capsid subunits. These widely-distributed interaction sites facilitate capsid-capsid interactions, which reciprocally mediate RNA folding and tertiary compression of RNA genome. Subsequently, this can be followed by continuous recruitment of capsids on folded RNA to finalize encapsidation process.

Several PAR-CL sites also aligned with, or in close adjacent to, known RNA motifs (**Figure 8**). On RNA 1 (**Figure 8a**), we could not align any candidate PAR-CL signal to subgenomic RNA 3, which suggests the possibility that the exclusion of RNA 3 during packaging is due to lack of strong RNA-capsid interactions. Interestingly, two most significant PAR-CL sites on RNA 1, U159 and U1233, aligned with previously discovered replication regulatory elements: a 5’ cis element (nts. 68-205) that is essential for RNA 1 replication and mitochondria-targeting(46), and short distal subgenomic control cis-element (nts. 1229-1239) which mediates subgenomic RNA 3 replication (47). Furthermore, U2515 and U2576 were located in the subgenomic promoter region (47, 58) which are also adjacent to a RNA 1 internal cis-acting replication element (intRE, nts 2322-2501) (47). Similarly, on RNA 2 (**Figure 8b**), we noticed PAR-CL site U534 is adjacent to a RNA 2 cis-acting regulatory site (59), and U1155 which is within a site required for specific packaging of both RNAs (18). A previously predicted stem loop site (nts 168-249) on RNA 2 serves as a RNA 2 packaging signal (24). This is also the only established FHV RNA packaging signal to date. Our DMS-MaPseq map did predict near identical stem loop structure as previous proposed and we noticed three significant (>1σ) PAR-CL sites were clustered in this critical region (**Supplemental Figure S9**). Since these RNA-capsid interaction sites are correlated to RNA cellular replication/mitochondrial targeting sites, we suggest they might be multi-functional in virus life cycle, and there can be a strong synergy between protein A-mitochondria localization (9, 60, 61), RNA replication, and virus assembly.

It was speculated previously that FHV packaging and replication are coordinated events. When FHV and brome mosaic virus (BMV) were co-expressed in plant cells, assembled virions only packed their own respective viral RNAs (20). Intracellular protein-protein interactions between FHV replicase (protein A) and capsid were detected (21). It has also been shown that FHV can ensure genome assembly specificity only when capsids were translated from replicating viral RNAs (23). It was hence suggested that FHV encapsidation may be coupled with the RNA replication. Our FHV PAR-CL experiments directly implicated only one aspect of FHV biology: the RNA sites interacting with capsid proteins. However, upon further analysis and mutational assays, a number of PAR-CL sites clearly indicated their significance in FHV replication and regulation: U159 and U227 mutants showed severe deficiency of RNA 1 replication and virion production, while U1233 entirely abolished viral replication. This provides further evidence that FHV replication and packaging are not sequentially separated events, but rather a synchronized, highly inter-dependent processes. Furthermore, these RNA-capsid interactions are not only important in post-replicational/translational RNA packaging, but may also be essential for multiple aspects of virus early stage activities in host cells.

## Materials and Methods

### Cell culture and virus

*D. melanogaster* (S2) cells were regularly maintained and passaged with Schneider’s *Drosophila* Media (Gibco) containing 10% fetal bovine serum, 1 × Antibiotic-Antimycotic (Gibco), 1 × MEM non-essential amino acids solution, and 1 mM sodium pyruvate.

As described previously (34), wt Flock House virus (FHV) was generated by transfecting S2 cells with pMT plasmid vectors (Invitrogen) containing respective genomes (NC_004146 for RNA 1, and NC_004144 for RNA 2). Copper sulfate was used to induce the promoter 24 h post transfection, while viruses were allowed to accumulate until 3 days post induction to yield passage 0 (p0) virus/cell mixture. The p0 transfected cells and viruses were then used to inoculate naïve S2 cells in a T75 flask for 3 days to yield passage 1 viruses, which were purified and used as FHV inoculum in this study, unless otherwise mentioned.

All virus transfections, infections, and passages with S2 cell culture were maintained in 27°C incubator, unless otherwise mentioned.

To purify FHV, 1% Triton X-100 was added to the cell culture containing p1 viruses. Cell culture underwent one freeze-thaw cycle, and cell debris was removed with 3000 × g centrifugation. FHV in the supernatant was crudely purified with 4% polyethylene glycol (PEG) 8000 and centrifuged (6000 × g) to remove debris (8). This was followed by DNase I and RNase A overnight digestion, to remove any co-precipitated cellular DNA or RNA. Unless otherwise mentioned, viruses were further purified with a 10-40% sucrose gradient, and ultracentrifuge at 40,000 RPM for 1.5 h. Viruses were then concentrated with 100K MWCO polyethersulfone (PES) membrane protein concentrator (Pierce) and washed three times with 10mM Tris pH 7.4.

### PAR-CL and ClickSeq

S2 cells were maintained in T75 flask until 70% - 90% confluency. Cells were infected with purified Flock House virus (p1) at MOI = 1 (34, 62, 63). As an initial dose, 4-thiouridine (Sigma-Aldrich) was supplemented to the cell culture to 100μM as 1 × concentration with virus. An optional “boost” dose of 4-thiouridine can also be supplied 16 h post infection (**Figure 3a**). Cells and viruses were harvested at 16 or 40 h post infection (**Figure 3a**). Viruses were purified with methods described above.

The nuclease-treated and purified 4SU-containing viruses were placed uncovered over ice and irradiated with 0.15 J/cm^2^ (26, 30) of 365nm UV light (3UV-38, UVP). After crosslink, viruses were digested with 8U of proteinase K (NEB) at 37 °C for 30 min. Crosslinked RNAs were extracted and purified with RNA Clean & Concentrator (Zymo Research) to yield RNA template for 4SU+/UV+ sequencing library sample.

Unless otherwise mentioned, the same 4SU-containing virus without any UV irradiation were prepared in the same way to give RNA template for 4SU+/UV− control library.

Both the crosslinked and uncrosslinked viral RNA were used to construct the ClickSeq Illumina libraries per standard ClickSeq method, which is detailed previously (34–36). 250ng of RNA template was used in reverse transcription reaction with 1:35 Azido-NTPs:dNTPs ratio and SuperScript III reverse transcriptase (Invitrogen).

Equal molar of each indexed library was pooled and run on a HiSeq 1500 platform (Illumina), with single read rapid run flowcell for 1×150 reads and 7 nucleotides for the index.

### DMS-MaPseq and ClickSeq

Dimethyl sulfate (DMS) RNA methylation method was described previously(40, 42). In this study, nuclease-treated and purified FHV was supplemented with DMS to 5% final concentration. After 5 min incubation at 30°C, reaction was quenched on ice for 5 min with 2 volumes of 10mM Tris pH 7.4 and 30% 2-Mercaptoethanol (BME). RNA extraction was conducted with Quick-RNA Viral Kits (Zymo Research) with additional BME in the extraction buffer. The DMS-control sample comprises the same virus stock with identical treatments as above, but without DMS supplement.

Methylated FHV RNA and respective controls were proceeded with similar ClickSeq library construction method. One exception is the use of a high-fidelity and processive thermostable group II reverse transcriptase enzyme (TGIRT-III, InGex) during reverse transcription. 100U of TGIRT-III was mixed with 250ng of RNA template, 0.5mM of AzNTPs/dNTPs mixture (AzNTPs:dNTPs = 1:35), and the following reaction conditions: 5 mM Dithiothreitol (Invitrogen), 10 U RNaseOUT (Invitrogen), 50 mM Tri–HCl pH 8.0, 75 mM KCl, and 3 mM MgCl2. The reaction mix was incubated in room temperature for 10 min, followed by 57°C incubation for 1.5 hrs, and 75°C termination for 15 min. The terminated reaction was then digested with RNase H to remove RNA template. The purified cDNA was proceeded with click reaction with Illumina adapters and final PCR amplification with indexes.

Library pooling and Illumina sequencing platform are the same as above.

### Bioinformatics and data analysis

The Illumina sequencing data of both PAR-CL and DMS-MaPseq were subjected to the following bioinformatic pipelines: first the Illumina sequencing adapter sequence “AGATCGGAAGAGC” was trimmed with *cutadapt* (37) (command line parameters: -b AGATCGGAAGAGC -m 40); then, we used *FASTX toolkit* (http://hannonlab.cshl.edu/fastx_toolkit/index.html) to remove the remaining random nucleotides from the Illumina adapter sequence and random base-pairing as a result of azide-alkyne cycloaddition from cDNA fragments (command line parameters: fastx_trimmer -Q33 -f 7); a further quality filter was applied to remove any reads that contained more than 4% nucleotides with a PHRED score <20 (command line parameters: fastq_quality_filter -Q33 -q 20 -p 96). The remaining reads were aligned to FHV genomes (NC_004146, and NC_004144). Data generated from PAR-CL experiments were aligned for end-to-end matches with *Bowtie (v1.0.1)* (38) (command line parameters: -v 2 --best). Data generated from DMS-MaPseq experiment were aligned with *Bowtie2* (64) to allow longer and gapped alignments (command line parameters: --local). Using *SAMtools* (65), the aligned reads were binarily converted, merged, indexed, sorted, and mathematically noted.

For both PAR-CL and DMS-MaPseq, we excluded any nucleotide location with less than 10k coverage to ensure reliable mutation rate calculation. For PAR-CL, we calculated the mutation frequencies of each of the four nucleotides, as well as the U-C mutation rate at each genomic U position. For DMS-MaPseq, similar analysis was conducted but we focused on the overall mutation rates of A and C genomic positions. Between test group and respective control (4SU+/UV+ and 4SU+/UV− for PAR-CL, DMS+ and DMS− for DMS-MaPseq), we compared the mutation rate at the same genomic position, to yield the fold change map, as presentations of PAR-CL or DMS-MaPseq signals.

A background filter was applied to both PAR-CL triplicates **(Figure 4a-d)** and DMS-MaPseq data **(Figure 5d, e)**, to ensure reliable data analysis and avoid potential false positives. For PAR-CL, the background threshold is determined by bottom 95% of U-C mutation rate in the uncrosslinked control group (4SU+,UV−). In the correspondent crosslinked group (4SU+,UV+), we removed any datapoint with U-C mutation rate below this threshold, as it is indistinguishable from background fluctuation. An example of applying background threshold for PAR-CL data can be found in **Supplemental data 3**. For DMS-MaPseq, similar background threshold was determined as the bottom 95% of A/C mutation rate, in the DMS-untreated control group (DMS−). Only the datapoints passed the background threshold were used to compile the fold change maps of mutation rate changes.

The raw sequencing data for both PAR-CL and DMS-MaPseq experiments are available in the NCBI sequence read archive (SRA) with accession number: PRJNA554838.

### RNA secondary structure prediction

RNA secondary structure prediction was conducted with RNAstructure (44) with 310.15 K temperature and maximum loop size = 30. “Fold” (44, 45) and “Partition” (66) were used to prediction the structure of RNA and calculate the base pairing probability, respectively. The most significant DMS-MaPseq signal sites were applied as unpaired constraints in structure prediction. No other constraints applied to the rest genomic sites, regardless of the DMS-MaPseq signals, to ensure the flexibility of algorithm. The predicted structure file was then re-organized and certain nucleotides were highlighted for graphical purposes with StructureEditor (v.1.0) which is also provided by RNAstructure suite.

### Mutated virus with disrupted PAR-CL sites

Plasmids containing FHV genomes were used as PCR templates. Universal upstream primer (TGCATAATTCTCTTACTGTCATGCCATCCGTAAG) and downstream primer (TAAGAGAATTATGCAGTGCTGCCATAACCATG) were used in combination with mutation primers (**Table 1**) to generate overlapped PCR fragments (Phusion High-Fidelity DNA Polymerase, NEB), with disrupted RNA structure at each selected PAR-CL sites. These overlapped fragments were then cloned into competent cells with standard In-Fusion HD Cloning (TaKaRa) techniques. The plasmids containing mutated FHV RNA 1 or RNA 2 sequences were then sanger-sequenced and mutation sites were confirmed.

To generate mutated viruses, the plasmids containing PAR-CL site mutations were used to transfect S2 cells with above-stated methods. Each mutant transfection consisted of equal amount of one mutated RNA genome with disrupted PAR-CL site, and wt genome of the other RNA (**Figure 7a**). These p0 mutant viruses were allowed to propagate in cell culture until 3 days post induction. Similar to before, p1 mutant viruses were generated by inoculating naïve S2 cells with p0 cell culture/virus mix.

### Relative virulence of mutant viruses

The virulence of p0 PAR-CL mutant viruses was measured via transfecting S2 cells with plasmids containing mutant viral genomes. For this experiment, transfection was conducted in black 96-well plate, with each well seeded with 25k S2 cells. The transfection reagents and methods were similar to above, except for scaling down to adapt for 96 well plate. For each mutant virus, 100ng plasmids of each mutant genome and the other wild type genome were used. alamarBlue was incubated with cell culture for 4h, before detecting fluorescence with EnSpire plate reader (PerkinElmer) at 560 nm excitation and 590 nm emission. The relative fluorescence then normalized reverse-ratiometrically with mock transfection = 0% and FHV wt RNA transfection = 100%.

The relative virulence of p1 PAR-CL mutant viruses was measured via infecting S2 cells with purified p1 mutant viruses. The p1 mutant viruses were purified through sucrose cushion (30% sucrose 10mM Tris pH 7.4, ultracentrifuge 80k rpm for 1.5hrs), PEG precipitation (4% v/v PEG8000), DNase I and RNase A treatment, buffer exchanged and concentrated with PES membrane protein concentrator (100K MWCO). The concentration of purified p1 viruses was determined with SDS-PAGE and densitometry. 25k S2 cells were seeded in black 96-well plate and 0.12 ng (approximately equivalent to MOI = 1 (34, 62, 63)) of serial diluted p1 viruses was used to infect each well. Standard alamarBlue assay was conducted as before, at 24 h post infection, to measure cell viability. The relative fluorescence then normalized reverse-ratiometrically with mock infection = 0% and purified wt FHV infection = 100%.

### RT-PCR

Transfected p0 PAR-CL mutants were sampled for RT-PCR to detect RNA production. Total RNA was extracted from transfected cells and media with Direct-zol RNA kit (Zymo Research), and DNase I in-column digestion was conducted to remove plasmids. For each mutants and controls, 200 ng of total RNA was used as template for RT-PCR. RT reaction was conducted with SuperScript III reverse transcriptase (Invitrogen), per manufacture’s protocol. PCR was conducted with OneTaq® (NEB), per manufacture’s protocol. The entire RT-PCR reaction was loaded on agarose gel for electrophoresis.

### SDS-PAGE and western blot

After collecting p0 transfections, the cell/virus/supernatant mix was centrifuged at 1000 × g for 10 min. Supernatant fraction was removed and collected thereafter. The cell pellet was washed once with 1 × PBS and centrifuged as before. The washed cellular fraction was then resuspended in 1 × PBS and 1 × cOmplete (Roche). 150 μL of supernatant (of each sample) was supplemented with 1 × cOmplete and then reduced with vacuum centrifuge prior to SDS-PAGE.

All SDS-PAGE assays were conducted with Bolt 4-12% Bis-Tris Plus Gels (Invitrogen). Membrane transfer was conducted with iBlot 2 Dry Blotting System (Invitrogen) with standard protocol. Western blot was conducted with iBind Western Device (Invitrogen) with standard protocol. Rabbit Anti-FHV polyclonal antibody was given as a gift from Dr. Vijay Reddy from Scripps Research, which was labelled with Alexa Fluor 488 goat anti-rabbit IgG (Invitrogen). Prior to membrane transfer, part of SDS-PAGE gel was cut and stained with Coomassie brilliant blue R-250 to highlight α-tubulin (55 kDa) as a loading control.

## Acknowledgements

A.R. is supported by start-up funds from the University of Texas Medical Branch. We thank Dr. Vijay Reddy from Scripps Research for providing samples of the anti-FHV capsid antibody.

