## Supplemental figures for "Mapping RNA-capsid interactions and RNA secondary structure within authentic virus particles using next-generation sequencing"

### Slide 1
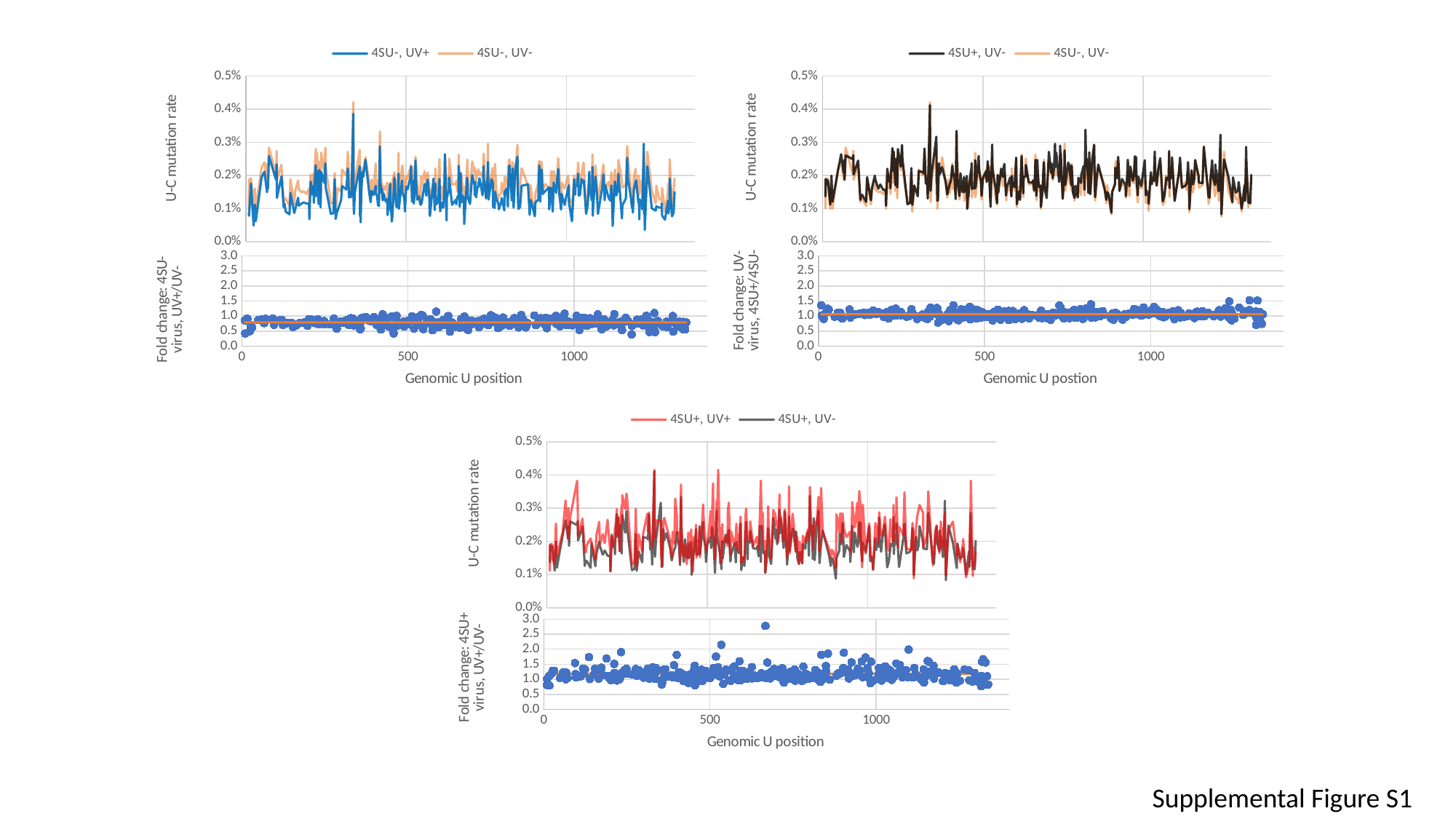

#### Chart
| Category | 4SU-, UV+ | 4SU-, UV- |
|---|---|---|
#### Chart
| Category | ave | 4A/4B |
|---|---|---|
#### Chart
| Category | 4SU+, UV- | 4SU-, UV- |
|---|---|---|
#### Chart
| Category | Ave | 3B/4B |
|---|---|---|
#### Chart
| Category | 4SU+, UV+ | 4SU+, UV- |
|---|---|---|
#### Chart
| Category | 3A/3B | ave |
|---|---|---|Supplemental Figure S1

### Slide 2
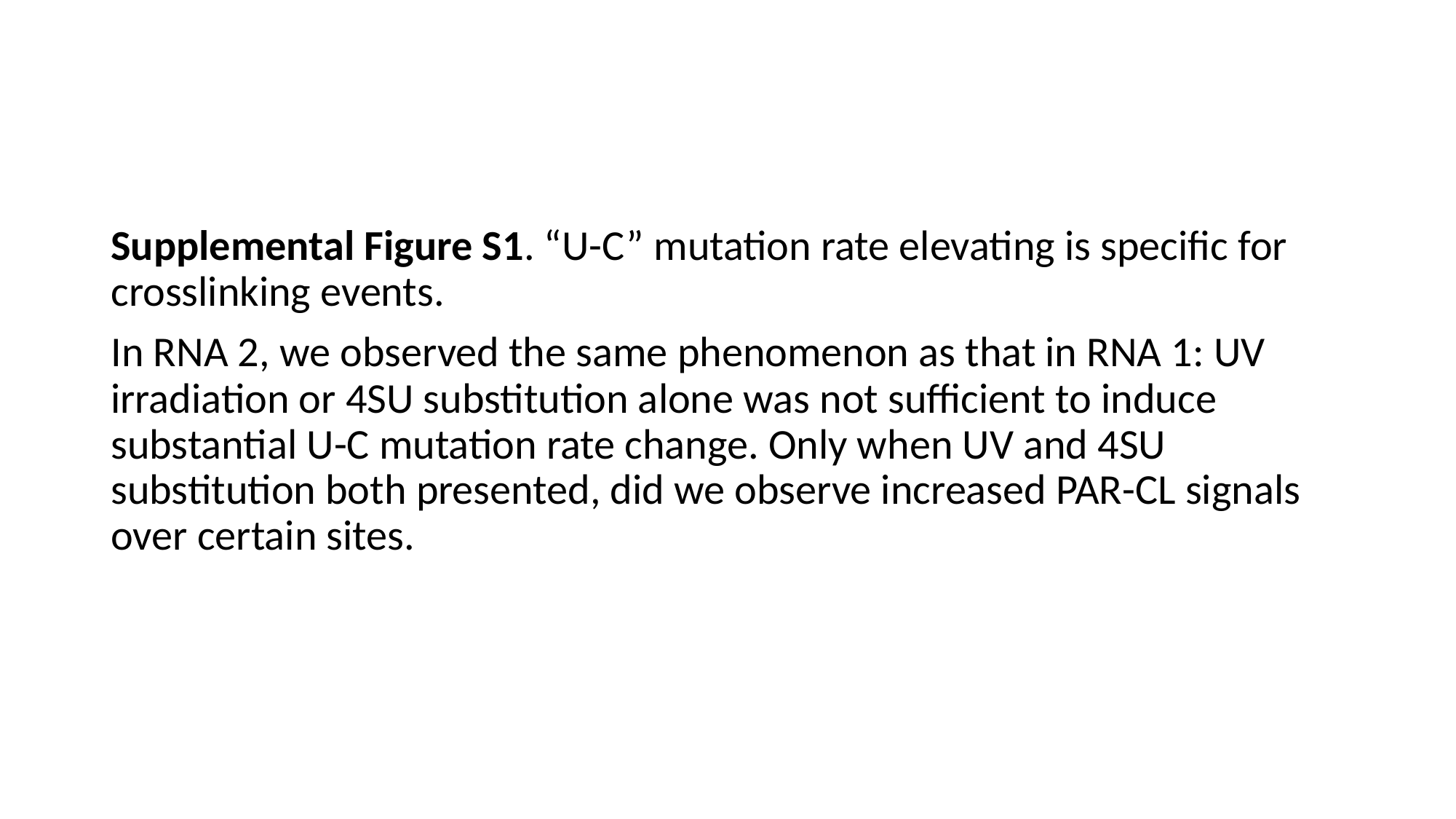

#
Supplemental Figure S1. “U-C” mutation rate elevating is specific for crosslinking events.
In RNA 2, we observed the same phenomenon as that in RNA 1: UV irradiation or 4SU substitution alone was not sufficient to induce substantial U-C mutation rate change. Only when UV and 4SU substitution both presented, did we observe increased PAR-CL signals over certain sites.

### Slide 3
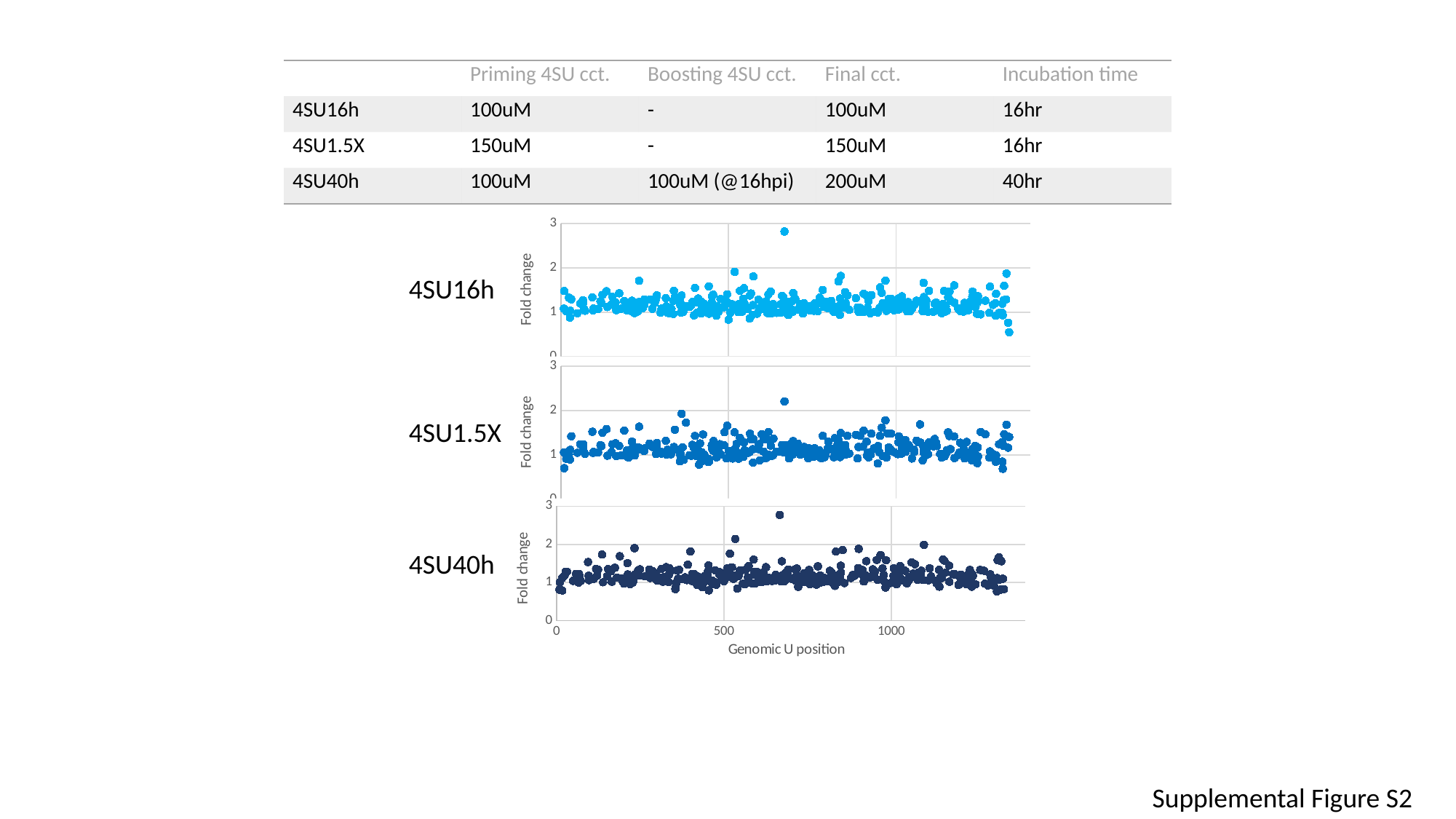

| | Priming 4SU cct. | Boosting 4SU cct. | Final cct. | Incubation time |
| --- | --- | --- | --- | --- |
| 4SU16h | 100uM | - | 100uM | 16hr |
| 4SU1.5X | 150uM | - | 150uM | 16hr |
| 4SU40h | 100uM | 100uM (@16hpi) | 200uM | 40hr |
#### Chart
| Category | 1A/1B |
|---|---|4SU16h
#### Chart
| Category | 2A/2B |
|---|---|4SU1.5X
#### Chart
| Category | 3A/3B |
|---|---|4SU40h
Supplemental Figure S2

### Slide 4
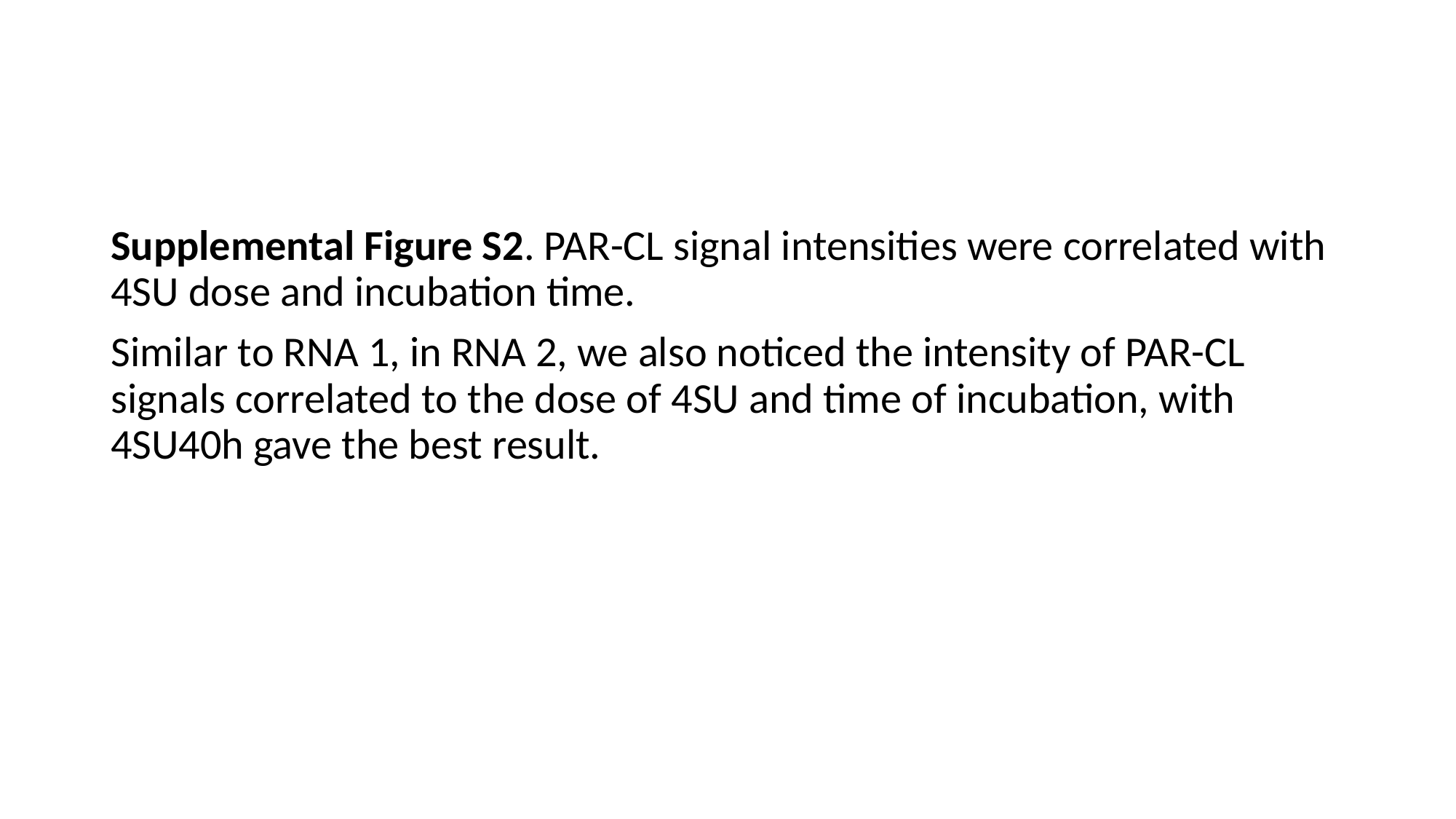

#
Supplemental Figure S2. PAR-CL signal intensities were correlated with 4SU dose and incubation time.
Similar to RNA 1, in RNA 2, we also noticed the intensity of PAR-CL signals correlated to the dose of 4SU and time of incubation, with 4SU40h gave the best result.

### Slide 5
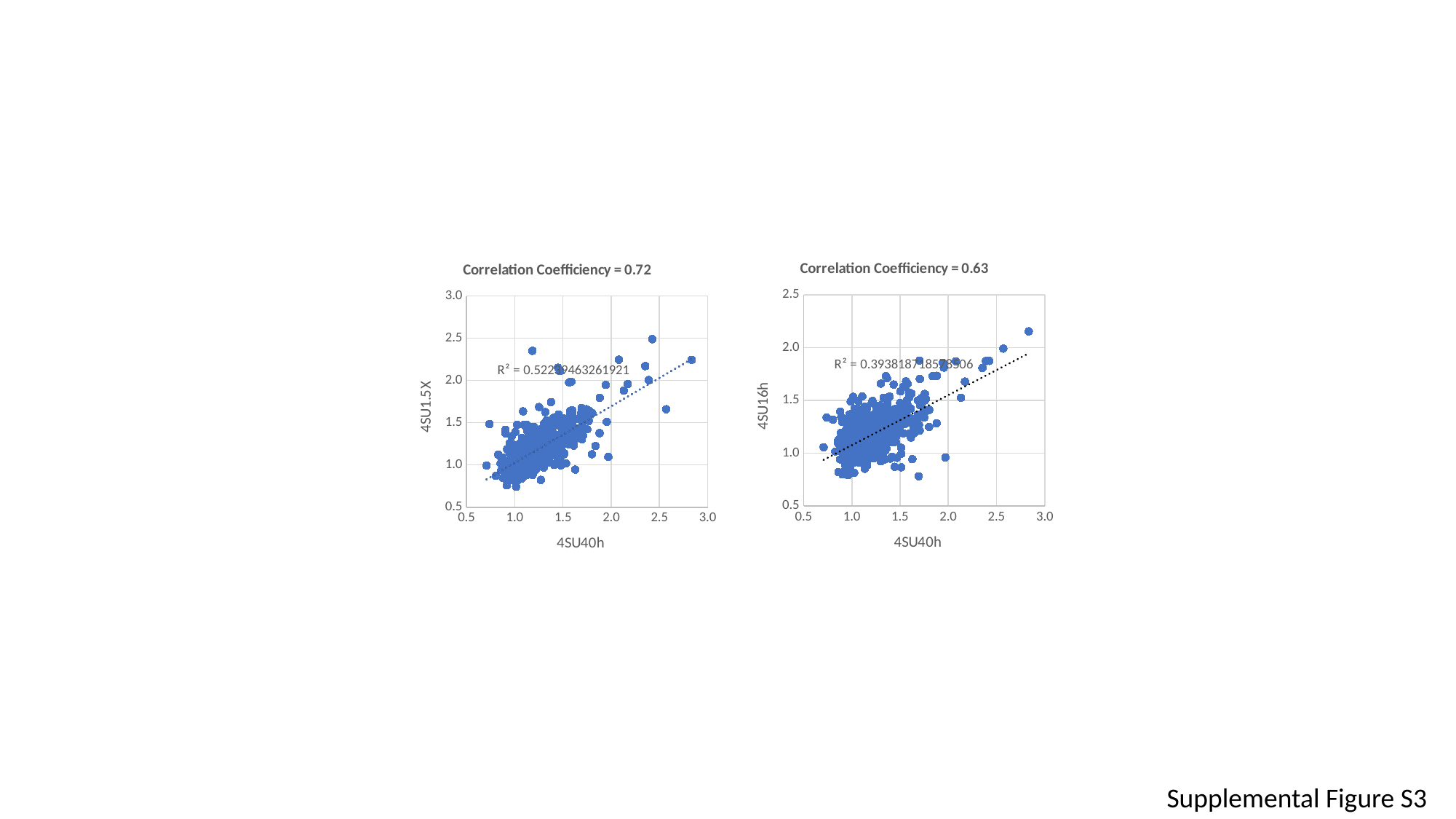

#### Chart: Correlation Coefficiency = 0.63
| Category | fold change: 1A/1B |
|---|---|
#### Chart: Correlation Coefficiency = 0.72
| Category | fold change:2A/2B |
|---|---|Supplemental Figure S3

### Slide 6
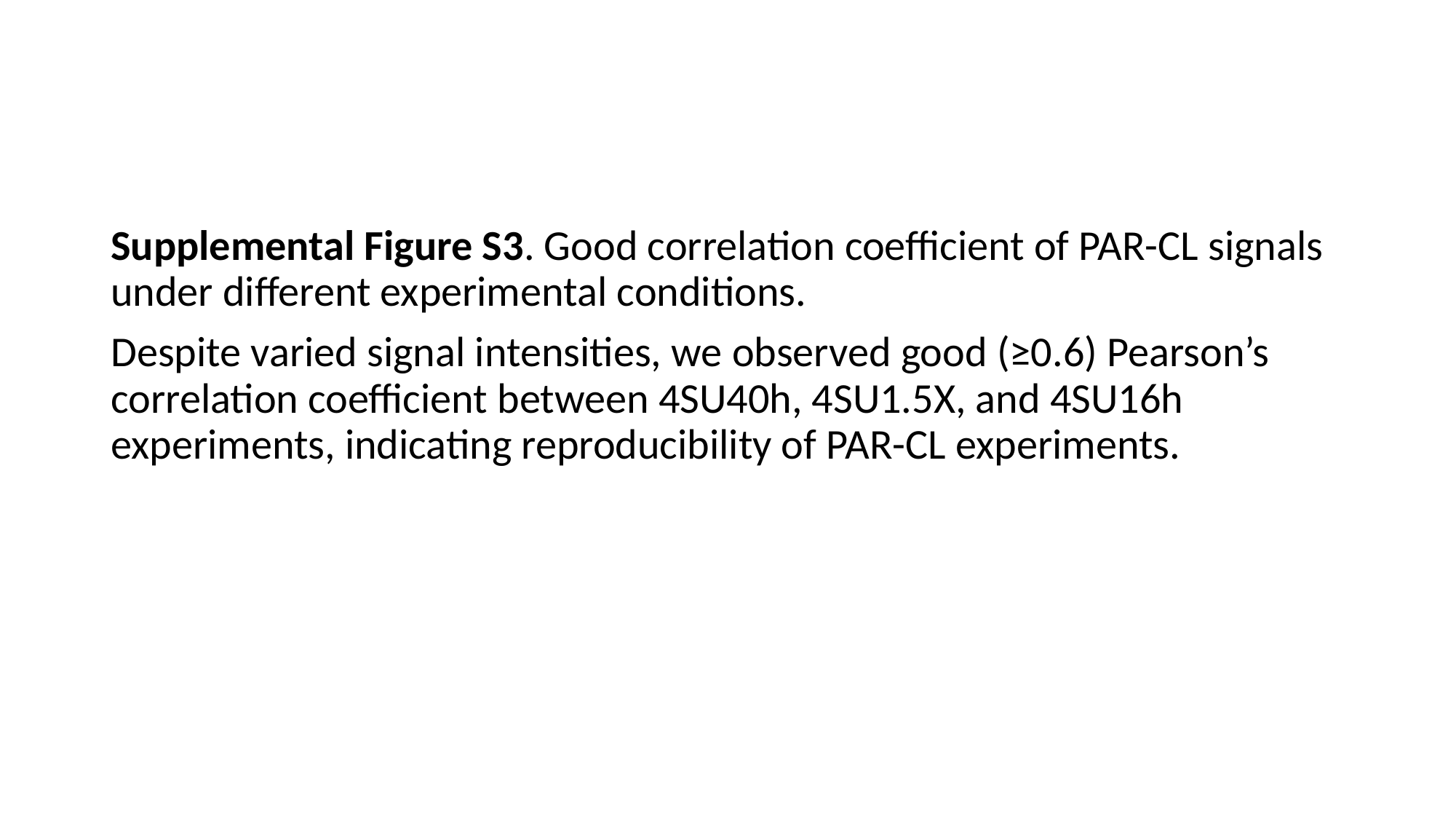

#
Supplemental Figure S3. Good correlation coefficient of PAR-CL signals under different experimental conditions.
Despite varied signal intensities, we observed good (≥0.6) Pearson’s correlation coefficient between 4SU40h, 4SU1.5X, and 4SU16h experiments, indicating reproducibility of PAR-CL experiments.

### Slide 7
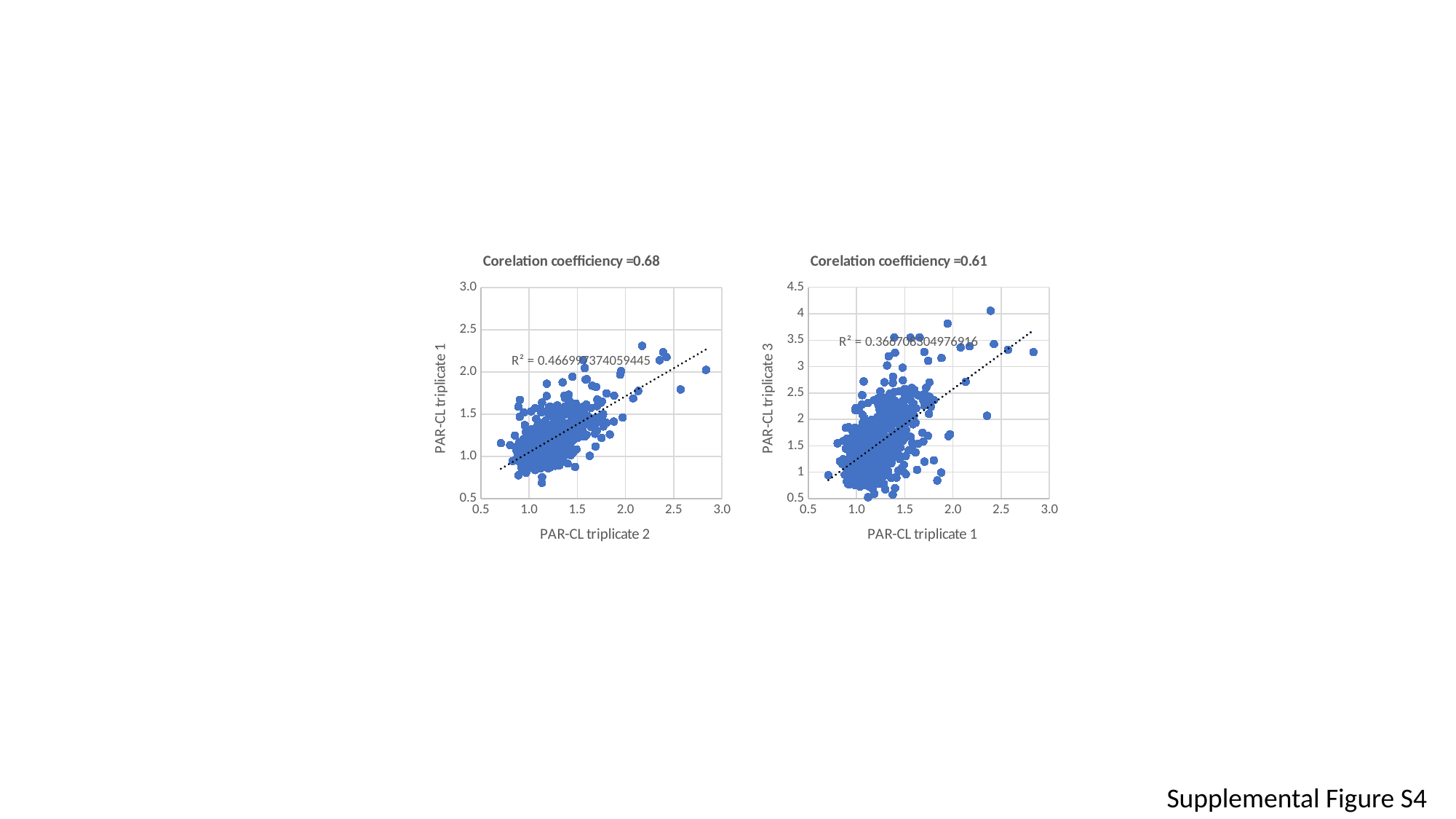

#### Chart: Corelation coefficiency =0.68
| Category | FCCD1 |
|---|---|
#### Chart: Corelation coefficiency =0.61
| Category | FCp3 |
|---|---|Supplemental Figure S4

### Slide 8
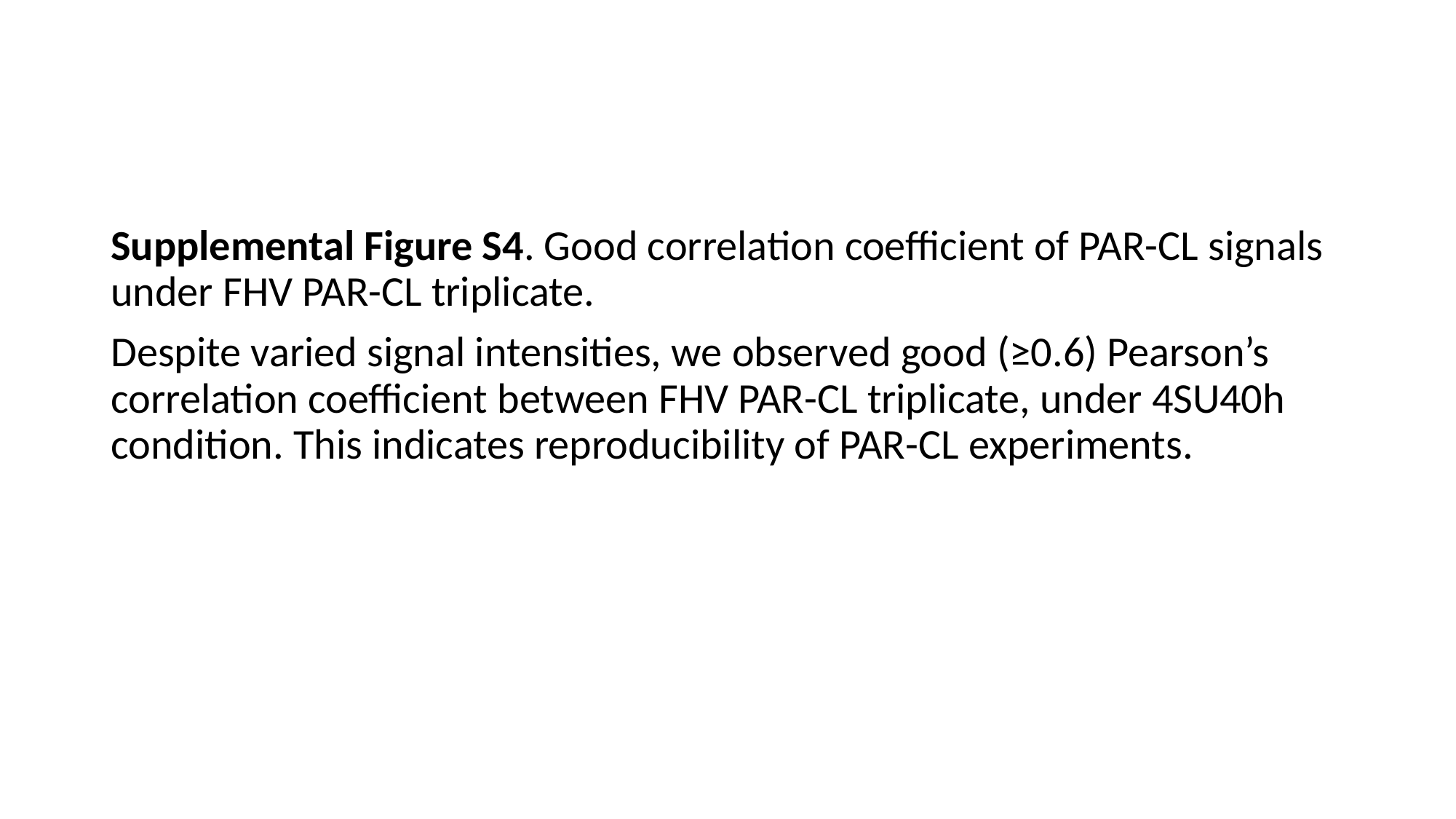

#
Supplemental Figure S4. Good correlation coefficient of PAR-CL signals under FHV PAR-CL triplicate.
Despite varied signal intensities, we observed good (≥0.6) Pearson’s correlation coefficient between FHV PAR-CL triplicate, under 4SU40h condition. This indicates reproducibility of PAR-CL experiments.

### Slide 9
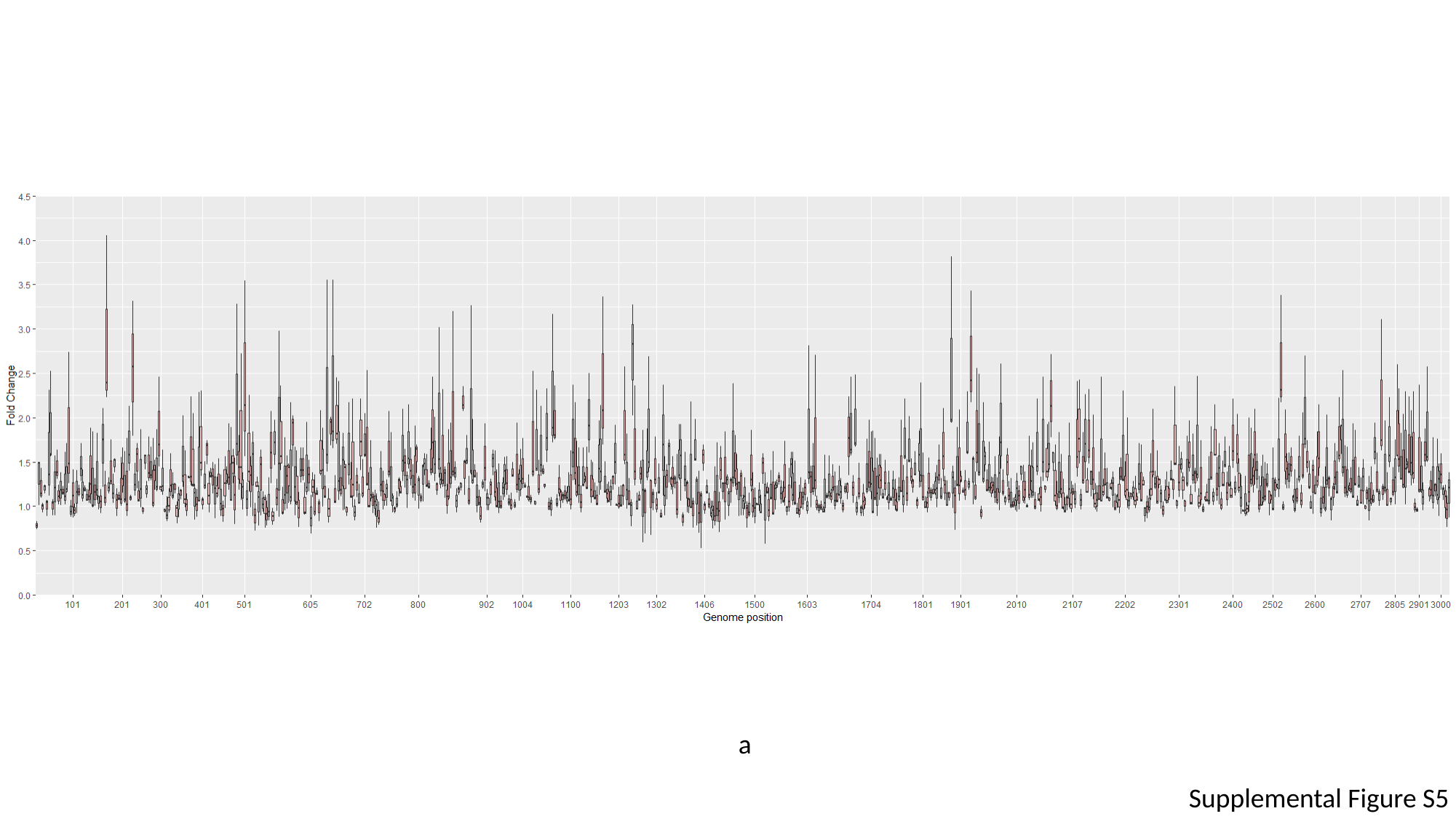

a
Supplemental Figure S5

### Slide 10
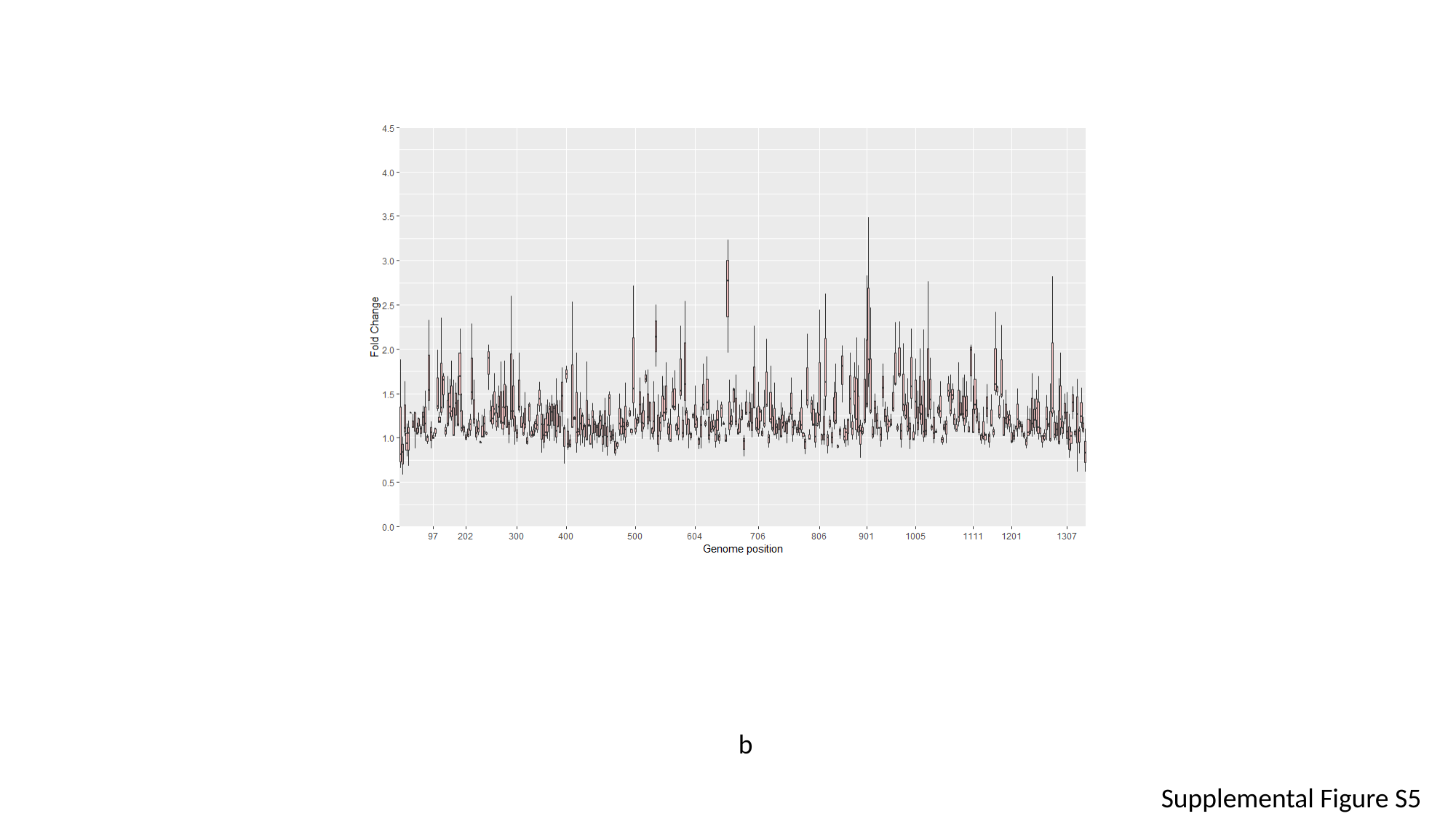

b
Supplemental Figure S5

### Slide 11
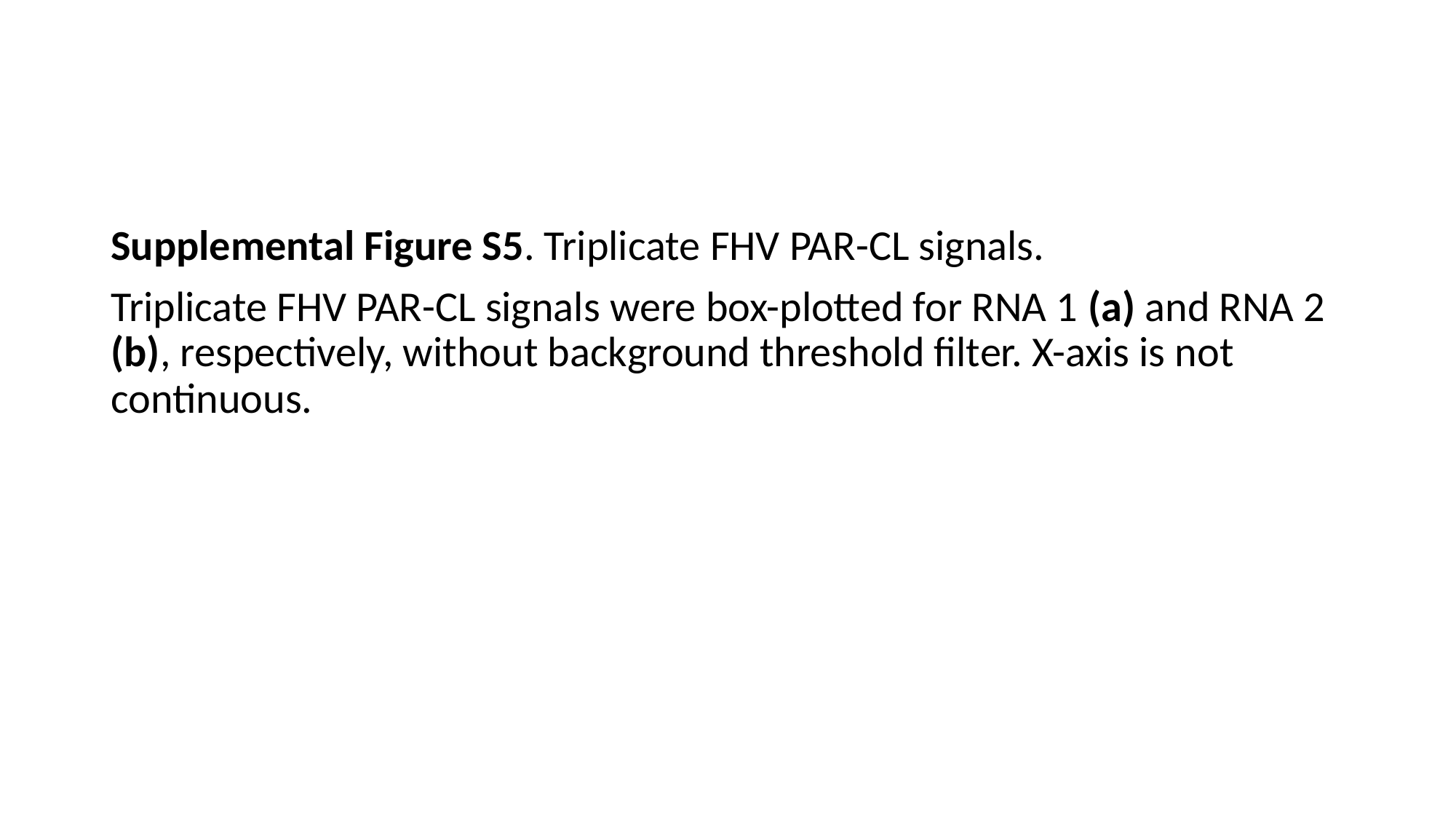

#
Supplemental Figure S5. Triplicate FHV PAR-CL signals.
Triplicate FHV PAR-CL signals were box-plotted for RNA 1 (a) and RNA 2 (b), respectively, without background threshold filter. X-axis is not continuous.

### Slide 12
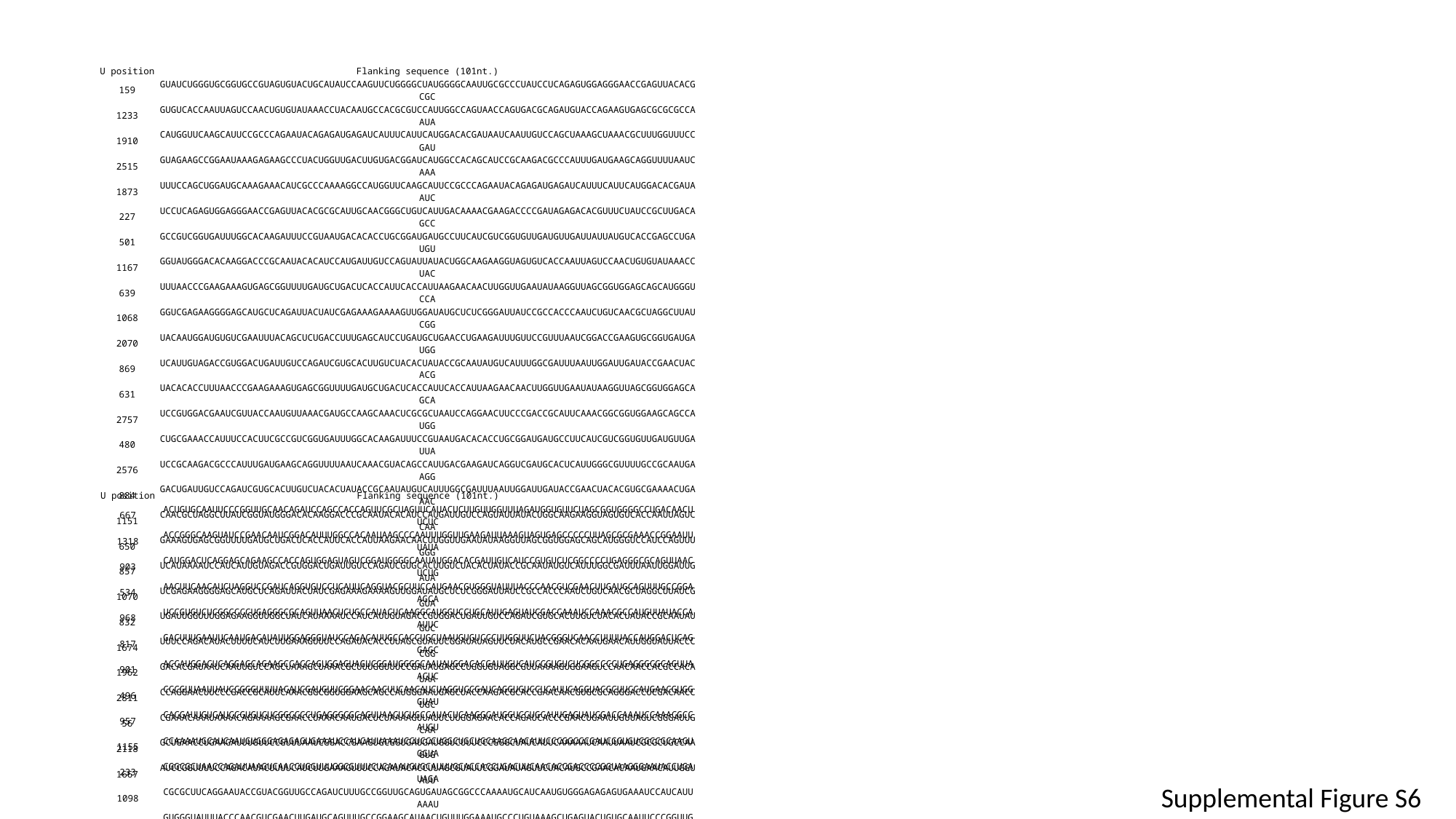

| U position | Flanking sequence (101nt.) |
| --- | --- |
| 159 | GUAUCUGGGUGCGGUGCCGUAGUGUACUGCAUAUCCAAGUUCUGGGGCUAUGGGGCAAUUGCGCCCUAUCCUCAGAGUGGAGGGAACCGAGUUACACGCGC |
| 1233 | GUGUCACCAAUUAGUCCAACUGUGUAUAAACCUACAAUGCCACGCGUCCAUUGGCCAGUAACCAGUGACGCAGAUGUACCAGAAGUGAGCGCGCGCCAAUA |
| 1910 | CAUGGUUCAAGCAUUCCGCCCAGAAUACAGAGAUGAGAUCAUUUCAUUCAUGGACACGAUAAUCAAUUGUCCAGCUAAAGCUAAACGCUUUGGUUUCCGAU |
| 2515 | GUAGAAGCCGGAAUAAAGAGAAGCCCUACUGGUUGACUUGUGACGGAUCAUGGCCACAGCAUCCGCAAGACGCCCAUUUGAUGAAGCAGGUUUUAAUCAAA |
| 1873 | UUUCCAGCUGGAUGCAAAGAAACAUCGCCCAAAAGGCCAUGGUUCAAGCAUUCCGCCCAGAAUACAGAGAUGAGAUCAUUUCAUUCAUGGACACGAUAAUC |
| 227 | UCCUCAGAGUGGAGGGAACCGAGUUACACGCGCAUUGCAACGGGCUGUCAUUGACAAAACGAAGACCCCGAUAGAGACACGUUUCUAUCCGCUUGACAGCC |
| 501 | GCCGUCGGUGAUUUGGCACAAGAUUUCCGUAAUGACACACCUGCGGAUGAUGCCUUCAUCGUCGGUGUUGAUGUUGAUUAUUAUGUCACCGAGCCUGAUGU |
| 1167 | GGUAUGGGACACAAGGACCCGCAAUACACAUCCAUGAUUGUCCAGUAUUAUACUGGCAAGAAGGUAGUGUCACCAAUUAGUCCAACUGUGUAUAAACCUAC |
| 639 | UUUAACCCGAAGAAAGUGAGCGGUUUUGAUGCUGACUCACCAUUCACCAUUAAGAACAACUUGGUUGAAUAUAAGGUUAGCGGUGGAGCAGCAUGGGUCCA |
| 1068 | GGUCGAGAAGGGGAGCAUGCUCAGAUUACUAUCGAGAAAGAAAAGUUGGAUAUGCUCUCGGGAUUAUCCGCCACCCAAUCUGUCAACGCUAGGCUUAUCGG |
| 2070 | UACAAUGGAUGUGUCGAAUUUACAGCUCUGACCUUUGAGCAUCCUGAUGCUGAACCUGAAGAUUUGUUCCGUUUAAUCGGACCGAAGUGCGGUGAUGAUGG |
| 869 | UCAUUGUAGACCGUGGACUGAUUGUCCAGAUCGUGCACUUGUCUACACUAUACCGCAAUAUGUCAUUUGGCGAUUUAAUUGGAUUGAUACCGAACUACACG |
| 631 | UACACACCUUUAACCCGAAGAAAGUGAGCGGUUUUGAUGCUGACUCACCAUUCACCAUUAAGAACAACUUGGUUGAAUAUAAGGUUAGCGGUGGAGCAGCA |
| 2757 | UCCGUGGACGAAUCGUUACCAAUGUUAAACGAUGCCAAGCAAACUCGCGCUAAUCCAGGAACUUCCCGACCGCAUUCAAACGGCGGUGGAAGCAGCCAUGG |
| 480 | CUGCGAAACCAUUUCCACUUCGCCGUCGGUGAUUUGGCACAAGAUUUCCGUAAUGACACACCUGCGGAUGAUGCCUUCAUCGUCGGUGUUGAUGUUGAUUA |
| 2576 | UCCGCAAGACGCCCAUUUGAUGAAGCAGGUUUUAAUCAAACGUACAGCCAUUGACGAAGAUCAGGUCGAUGCACUCAUUGGGCGUUUUGCCGCAAUGAAGG |
| 884 | GACUGAUUGUCCAGAUCGUGCACUUGUCUACACUAUACCGCAAUAUGUCAUUUGGCGAUUUAAUUGGAUUGAUACCGAACUACACGUGCGAAAACUGAAAC |
| 1151 | CAACGCUAGGCUUAUCGGUAUGGGACACAAGGACCCGCAAUACACAUCCAUGAUUGUCCAGUAUUAUACUGGCAAGAAGGUAGUGUCACCAAUUAGUCCAA |
| 650 | GAAAGUGAGCGGUUUUGAUGCUGACUCACCAUUCACCAUUAAGAACAACUUGGUUGAAUAUAAGGUUAGCGGUGGAGCAGCAUGGGUCCAUCCAGUUUGGG |
| 857 | UCAUAAAAUCCAUCAUUGUAGACCGUGGACUGAUUGUCCAGAUCGUGCACUUGUCUACACUAUACCGCAAUAUGUCAUUUGGCGAUUUAAUUGGAUUGAUA |
| 1070 | UCGAGAAGGGGAGCAUGCUCAGAUUACUAUCGAGAAAGAAAAGUUGGAUAUGCUCUCGGGAUUAUCCGCCACCCAAUCUGUCAACGCUAGGCUUAUCGGUA |
| 832 | UGAUUGGUUUGGAGAAGGUUGGCUAUCAUAAAAUCCAUCAUUGUAGACCGUGGACUGAUUGUCCAGAUCGUGCACUUGUCUACACUAUACCGCAAUAUGUC |
| 1674 | UUUCCAGACAUACUUUUCAUCUUGAAAGUUUCCAGAUACACCUUAGCGUAUUCGGAUAUAGUUCUACAUGCCGAACACAAUGAACAUUGGUAUUACCCCGG |
| 1962 | GACACGAUAAUCAAUUGUCCAGCUAAAGCUAAACGCUUUGGUUUCCGAUAUGAGCCUGGUGUAGGCGUUAAAAGUGGAAGUCCAACAACCACGCCACAUAA |
| 2811 | CCAGGAACUUCCCGACCGCAUUCAAACGGCGGUGGAAGCAGCCAUGGGAAUGAGCUACCAAGACGCACCGAACAACGUGCGCAGGGACCUCGACAACCUGC |
| 56 | CGAAACAAAUAAAACAGAAAAGCGAACCUAAACAAUGACUCUAAAAGUUAUUCUUGGAGAACACCAGAUCACCCGAACUGAAUUGUUAGUCGGGAUUGCAA |
| 2118 | GCUGAACCUGAAGAUUUGUUCCGUUUAAUCGGACCGAAGUGCGGUGAUGAUGGUCUUUCCCGGGCUAUCAUUCAAAAAUCAAUUAAUCGCGCUGCCAAGUG |
| 1667 | AUCCGGUUUUCCAGACAUACUUUUCAUCUUGAAAGUUUCCAGAUACACCUUAGCGUAUUCGGAUAUAGUUCUACAUGCCGAACACAAUGAACAUUGGUAUU |
| U position | Flanking sequence (101nt.) |
| --- | --- |
| 667 | ACUGUGCAAUUCCCGGUUGCAACAGAUCCAGCCACCAGUUCGCUAGUUCAUACUCUUGUUGGUUUAGAUGGUGUUCUAGCGGUGGGGCCUGACAACUUCUC |
| 1318 | ACCGGGCAAGUAUCCGAACAAUCGGACAUUUGGCCACAAUAAGCCCAAUUUGGUUGAAGAUUAAAGUAGUGAGCCCCCUUAGCGCGAAACCGGAAUUUAUA |
| 903 | CAUGGACUCAGGAGCAGAAGCCACCAGUGGAGUAGUCGGAUGGGGCAAUAUGGACACGAUUGUCAUCCGUGUCUCGGCCCCUGAGGGCGCAGUUAACUCUG |
| 534 | AACUUCAACAUCUAGGUCCGAUCAGGUGUCCUCAUUCAGGUACGCUUCCAUGAACGUGGGUAUUUACCCAACGUCGAACUUGAUGCAGUUUGCCGGAAGCA |
| 968 | UCCGUGUCUCGGCCCCUGAGGGCGCAGUUAACUCUGCCAUACUCAAGGCAUGGUCCUGCAUUGAGUAUCGACCAAAUCCAAACGCCAUGUUAUACCAAUUC |
| 817 | GACUUUGAAUUCAAUGACAUAUUGGAGGGUAUCCAGACAUUGCCACCUGCUAAUGUGUCCCUUGGUUCUACGGGUCAACCUUUUACCAUGGACUCAGGAGC |
| 901 | ACCAUGGACUCAGGAGCAGAAGCCACCAGUGGAGUAGUCGGAUGGGGCAAUAUGGACACGAUUGUCAUCCGUGUCUCGGCCCCUGAGGGCGCAGUUAACUC |
| 496 | CCCGUUAAUUAUCCGGGUUUUACAUCGAUGUUCGGAACAACUUCAACAUCUAGGUCCGAUCAGGUGUCCUCAUUCAGGUACGCUUCCAUGAACGUGGGUAU |
| 957 | CACGAUUGUCAUCCGUGUCUCGGCCCCUGAGGGCGCAGUUAACUCUGCCAUACUCAAGGCAUGGUCCUGCAUUGAGUAUCGACCAAAUCCAAACGCCAUGU |
| 1155 | CCAAAAUGCAUCAAUGUGGGAGAGAGUGAAAUCCAUCAUUAAAUCCUCCCUGGCUGCUGCAAGCAACAUUCCCGGCCCGAUCGGUGUCGCCGCAAGUGGUA |
| 233 | CGGCGCUAACCAGAUUAAGUCAACCUGGUUUGGCGUUUCUCAAAUGUGCAUUUGCACCACCUGACUUCAACACCGACCCCGGUAAGGGAAUACCUGAUAGA |
| 1098 | CGCGCUUCAGGAAUACCGUACGGUUGCCAGAUCUUUGCCGGUUGCAGUGAUAGCGGCCCAAAAUGCAUCAAUGUGGGAGAGAGUGAAAUCCAUCAUUAAAU |
| 589 | GUGGGUAUUUACCCAACGUCGAACUUGAUGCAGUUUGCCGGAAGCAUAACUGUUUGGAAAUGCCCUGUAAAGCUGAGUACUGUGCAAUUCCCGGUUGCAAC |
| 1266 | GUCAGCCCUUUUUGAAGGAUUUGGCUUUUAGAAGCAUCCGGACGCCAACCUAACCGGGCAAGUAUCCGAACAAUCGGACAUUUGGCCACAAUAAGCCCAAU |
| 835 | AUAUUGGAGGGUAUCCAGACAUUGCCACCUGCUAAUGUGUCCCUUGGUUCUACGGGUCAACCUUUUACCAUGGACUCAGGAGCAGAAGCCACCAGUGGAGU |
Supplemental Figure S6

### Slide 13
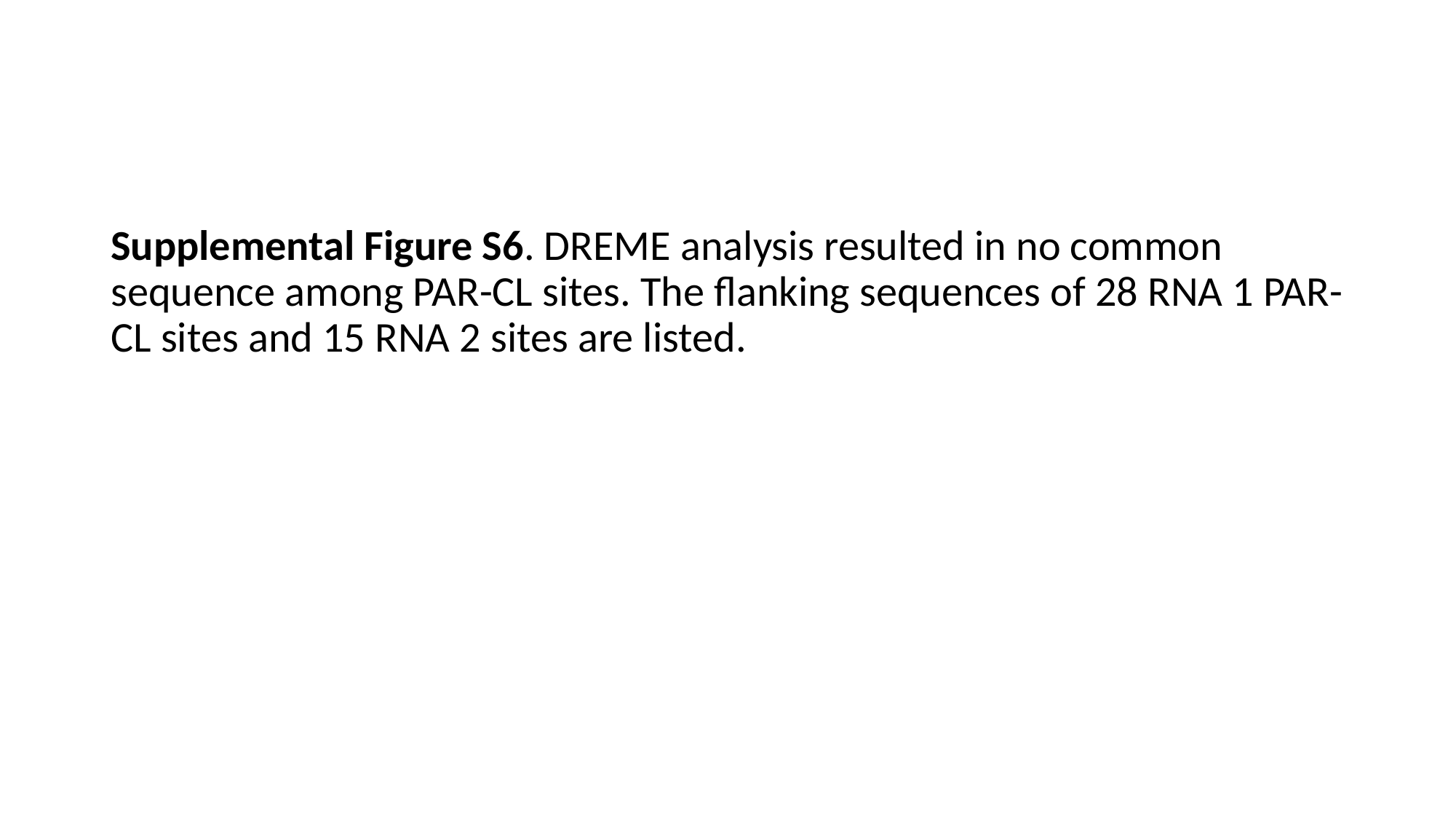

#
Supplemental Figure S6. DREME analysis resulted in no common sequence among PAR-CL sites. The flanking sequences of 28 RNA 1 PAR-CL sites and 15 RNA 2 sites are listed.

### Slide 14
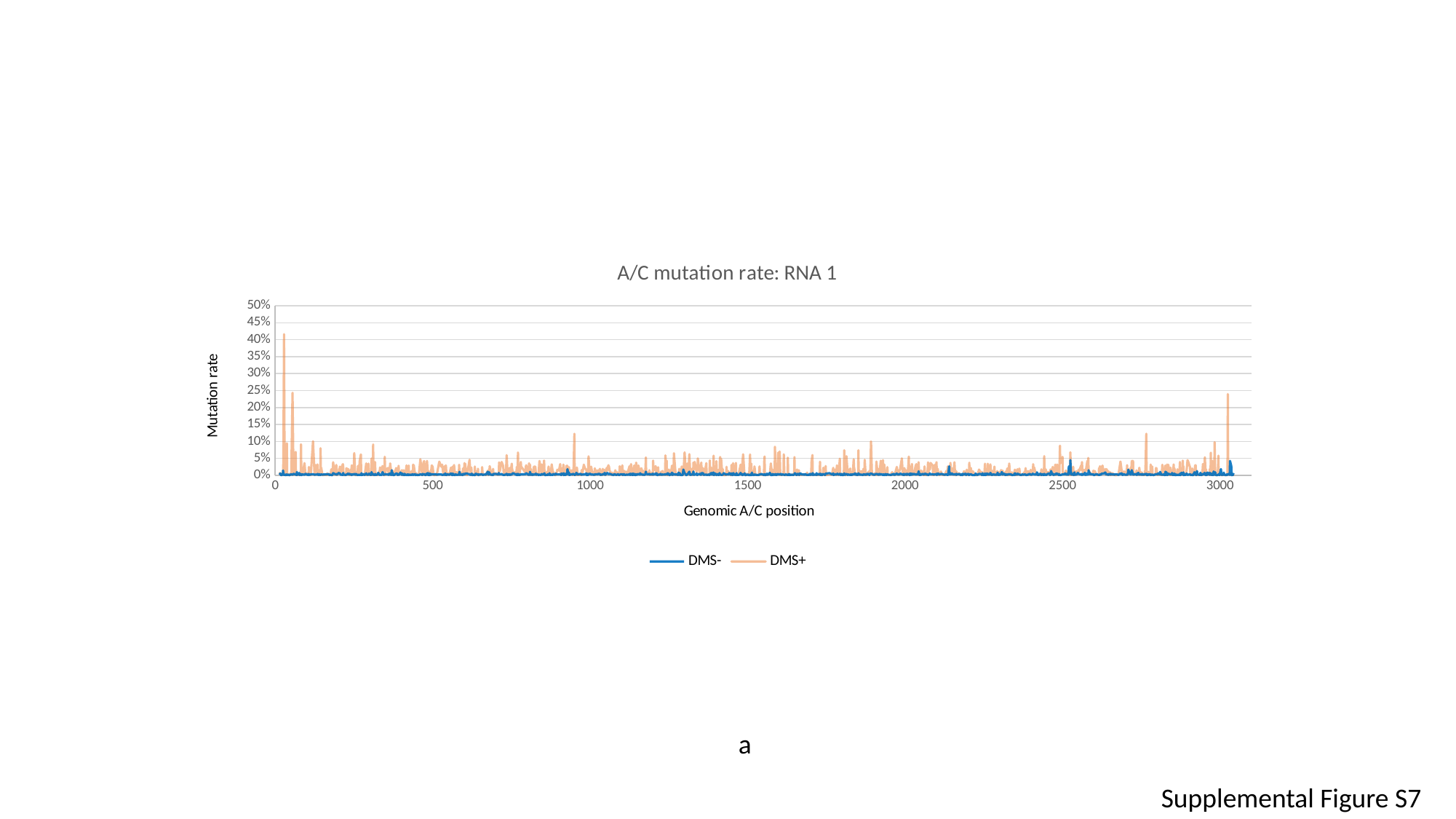

#### Chart: A/C mutation rate: RNA 1
| Category | DMS- | DMS+ |
|---|---|---|a
Supplemental Figure S7

### Slide 15
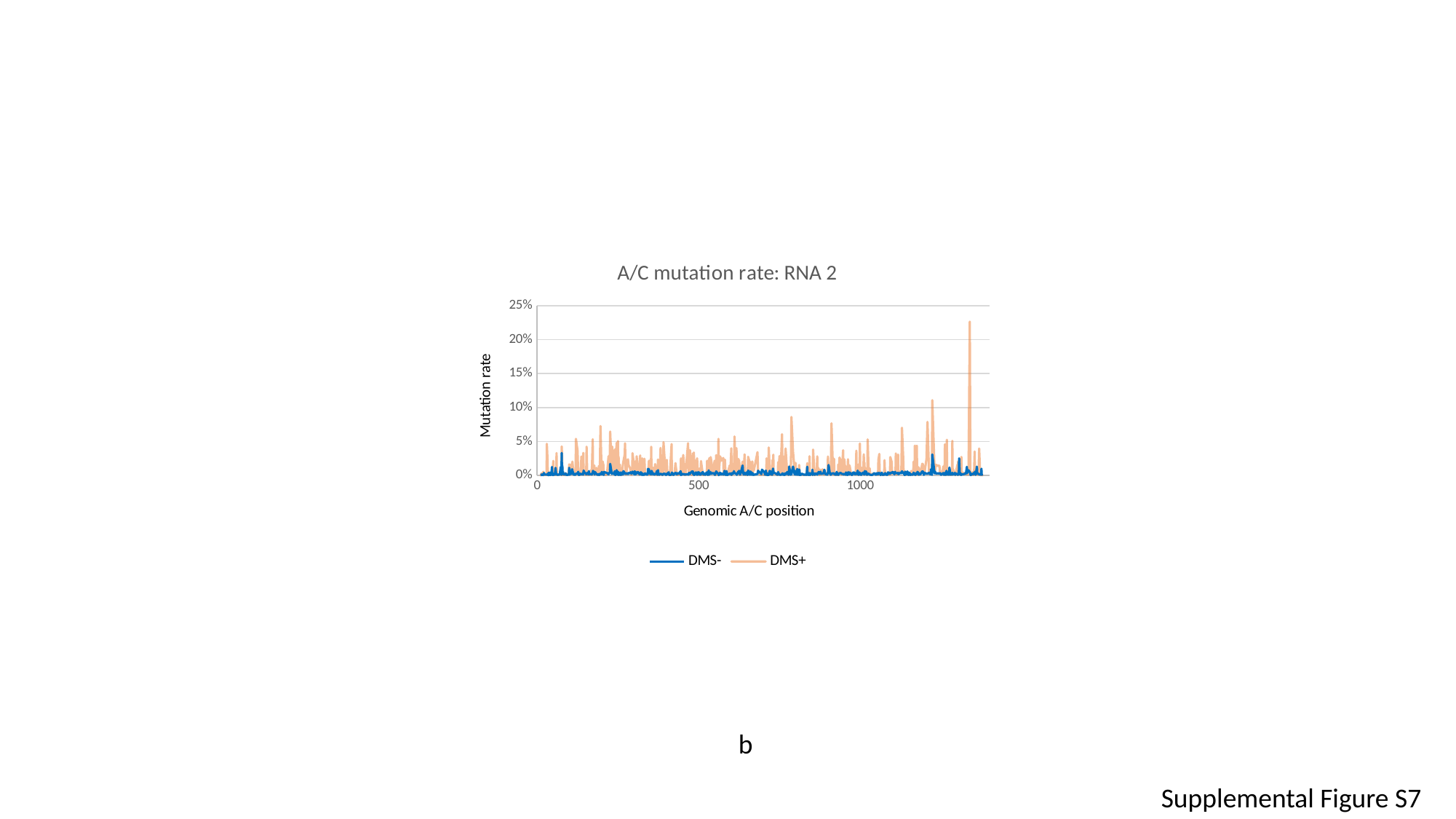

#### Chart: A/C mutation rate: RNA 2
| Category | DMS- | DMS+ |
|---|---|---|b
Supplemental Figure S7

### Slide 16
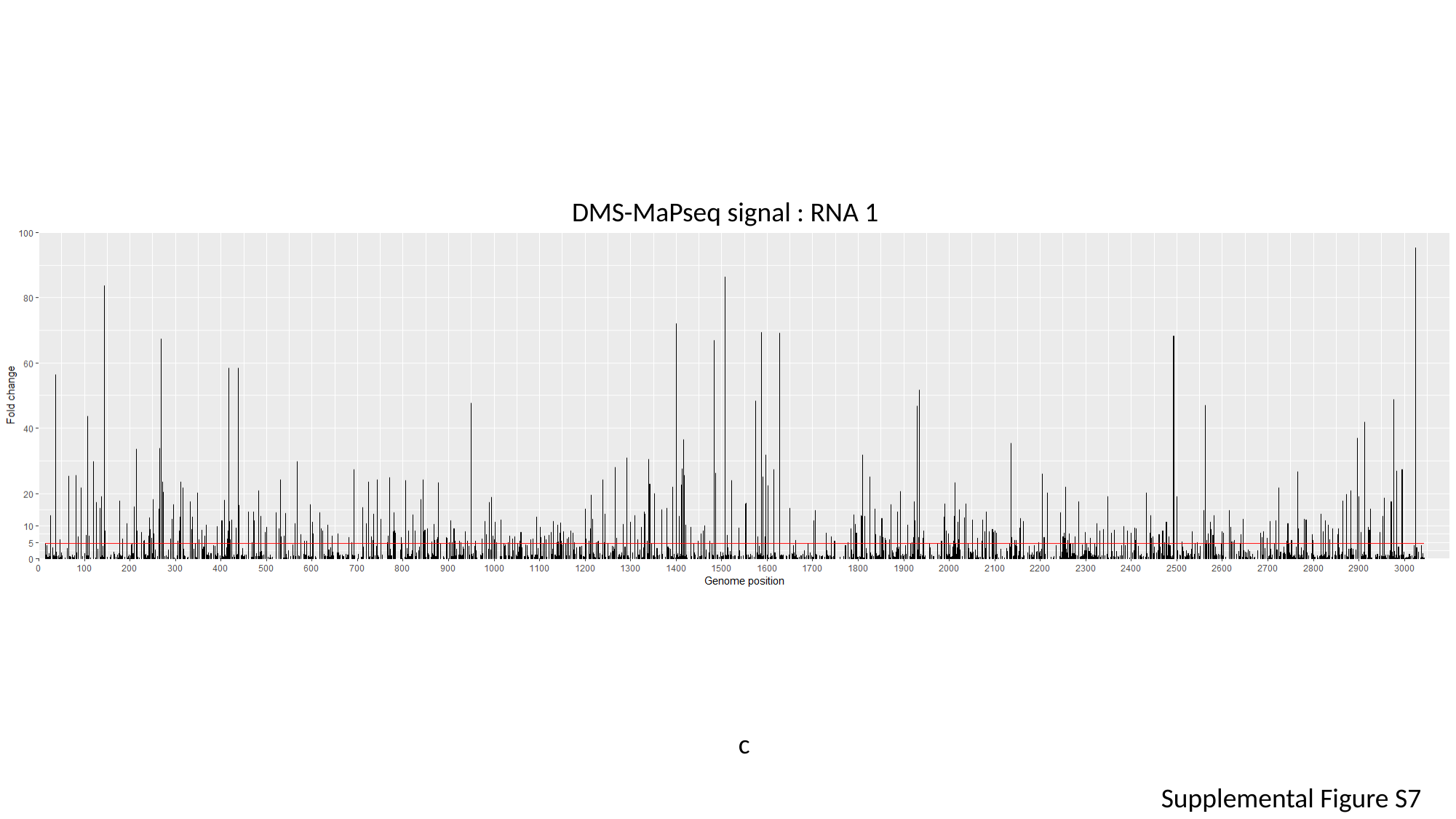

DMS-MaPseq signal : RNA 1
c
Supplemental Figure S7

### Slide 17
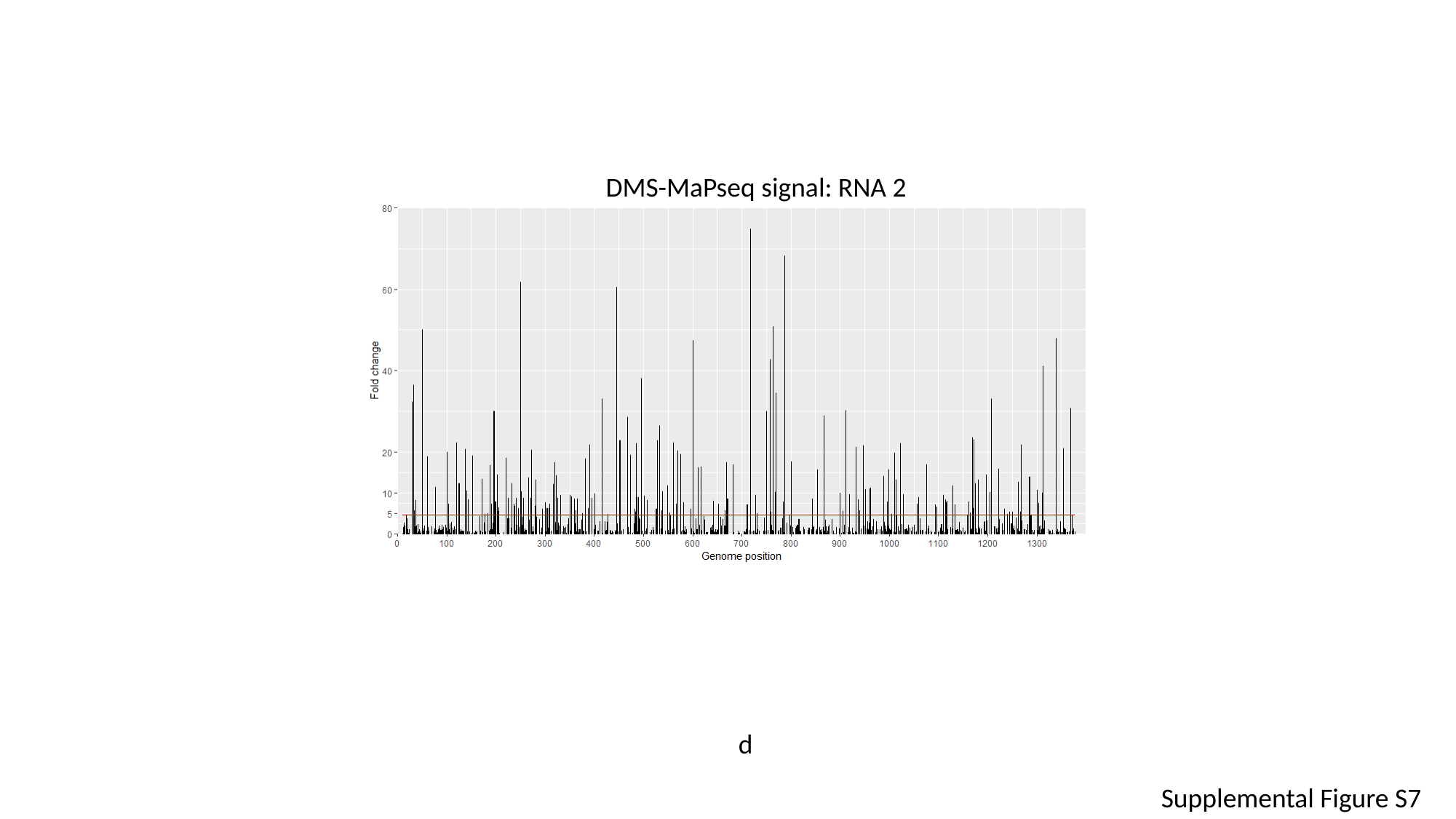

DMS-MaPseq signal: RNA 2
d
Supplemental Figure S7

### Slide 18
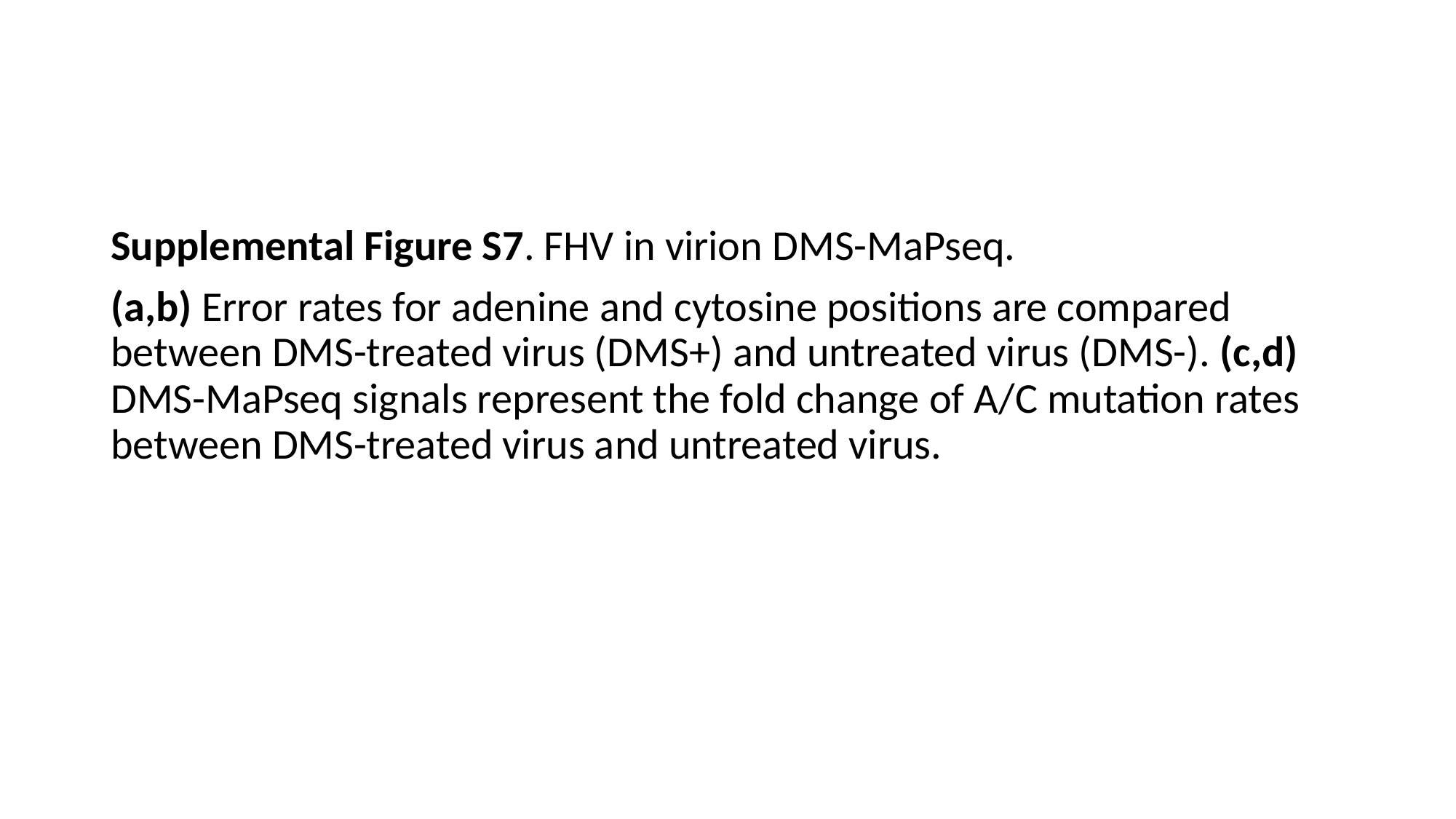

#
Supplemental Figure S7. FHV in virion DMS-MaPseq.
(a,b) Error rates for adenine and cytosine positions are compared between DMS-treated virus (DMS+) and untreated virus (DMS-). (c,d) DMS-MaPseq signals represent the fold change of A/C mutation rates between DMS-treated virus and untreated virus.

### Slide 19
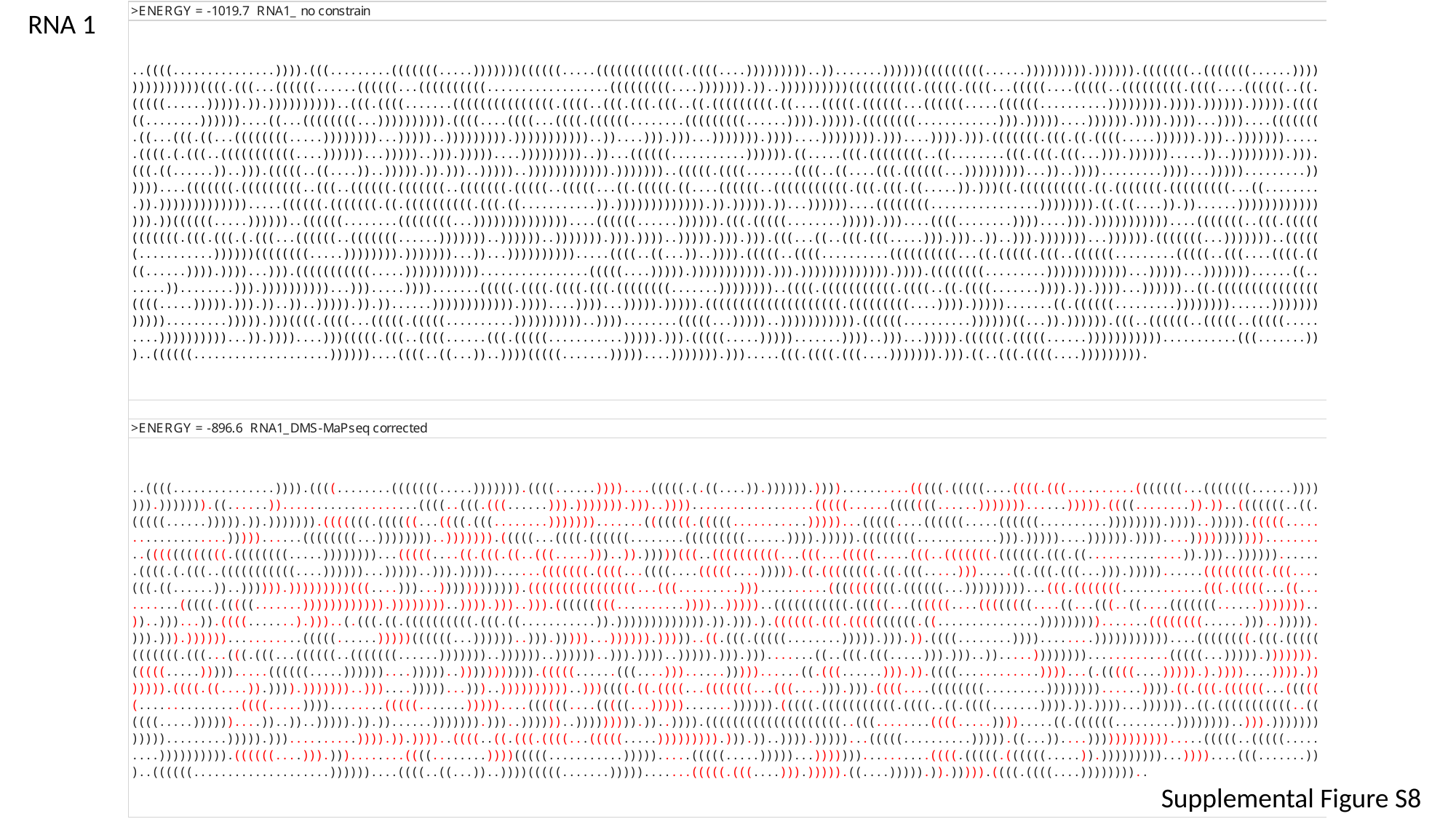

RNA 1
Supplemental Figure S8

### Slide 20
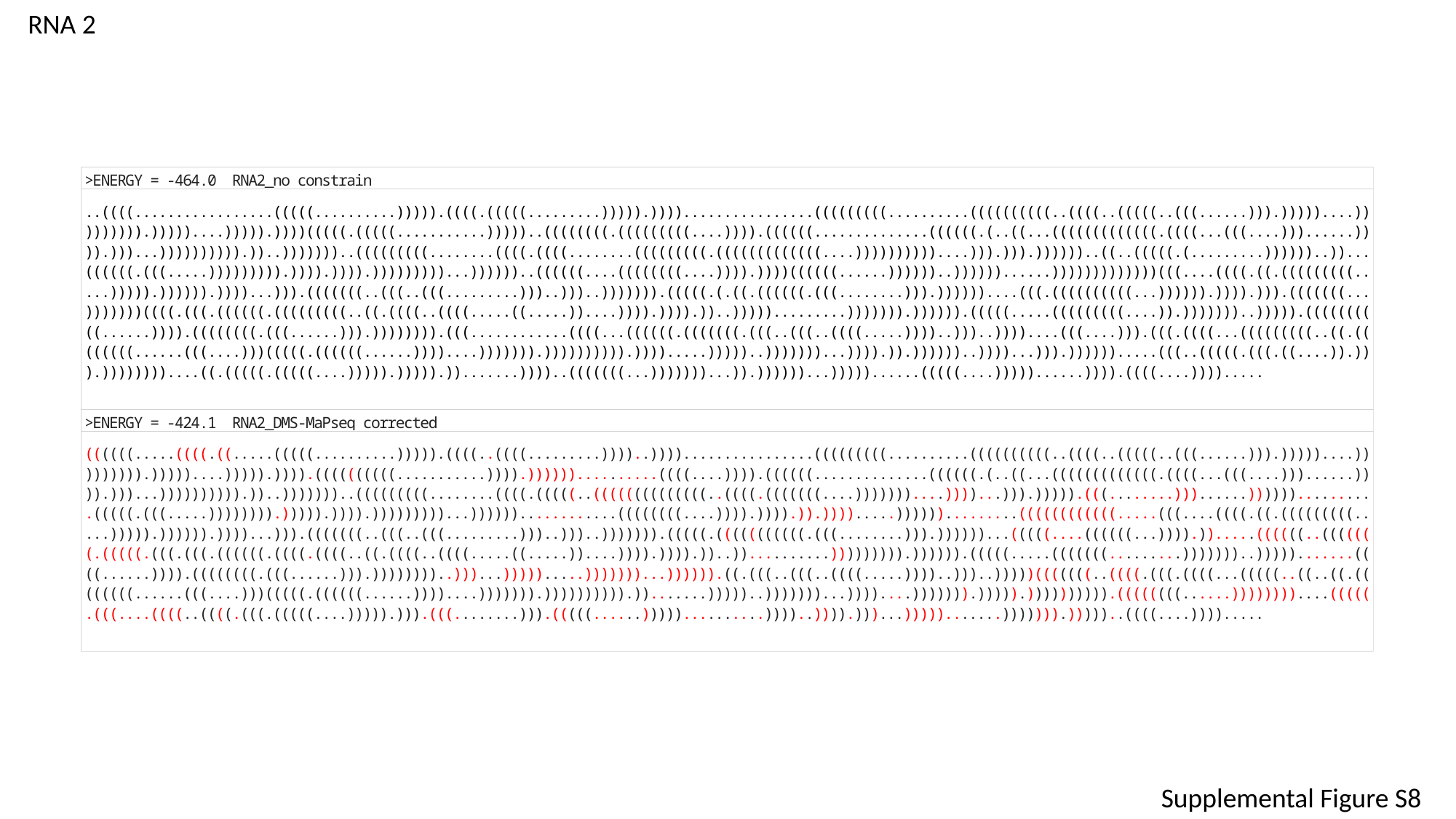

RNA 2
Supplemental Figure S8

### Slide 21
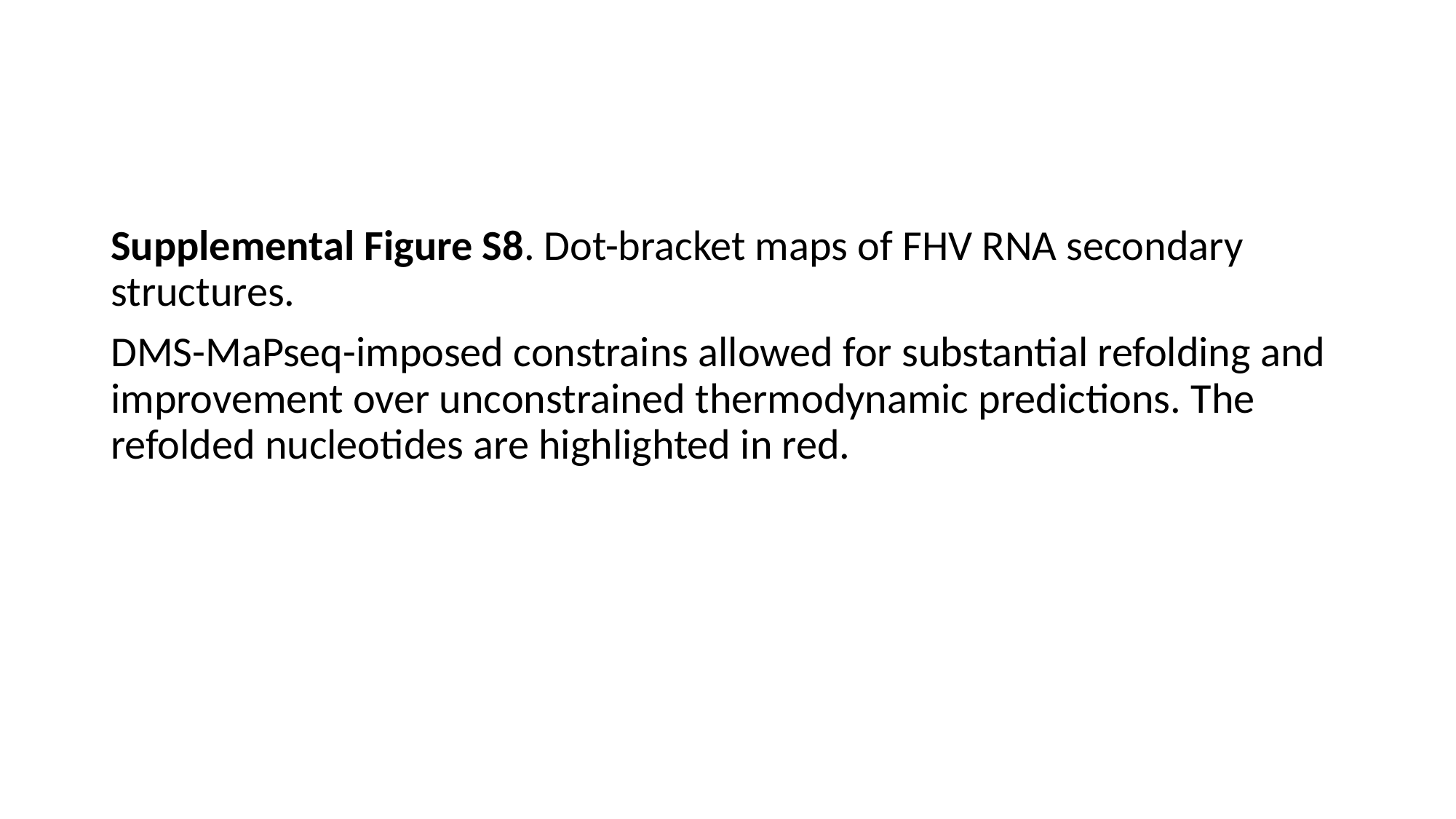

#
Supplemental Figure S8. Dot-bracket maps of FHV RNA secondary structures.
DMS-MaPseq-imposed constrains allowed for substantial refolding and improvement over unconstrained thermodynamic predictions. The refolded nucleotides are highlighted in red.

### Slide 22
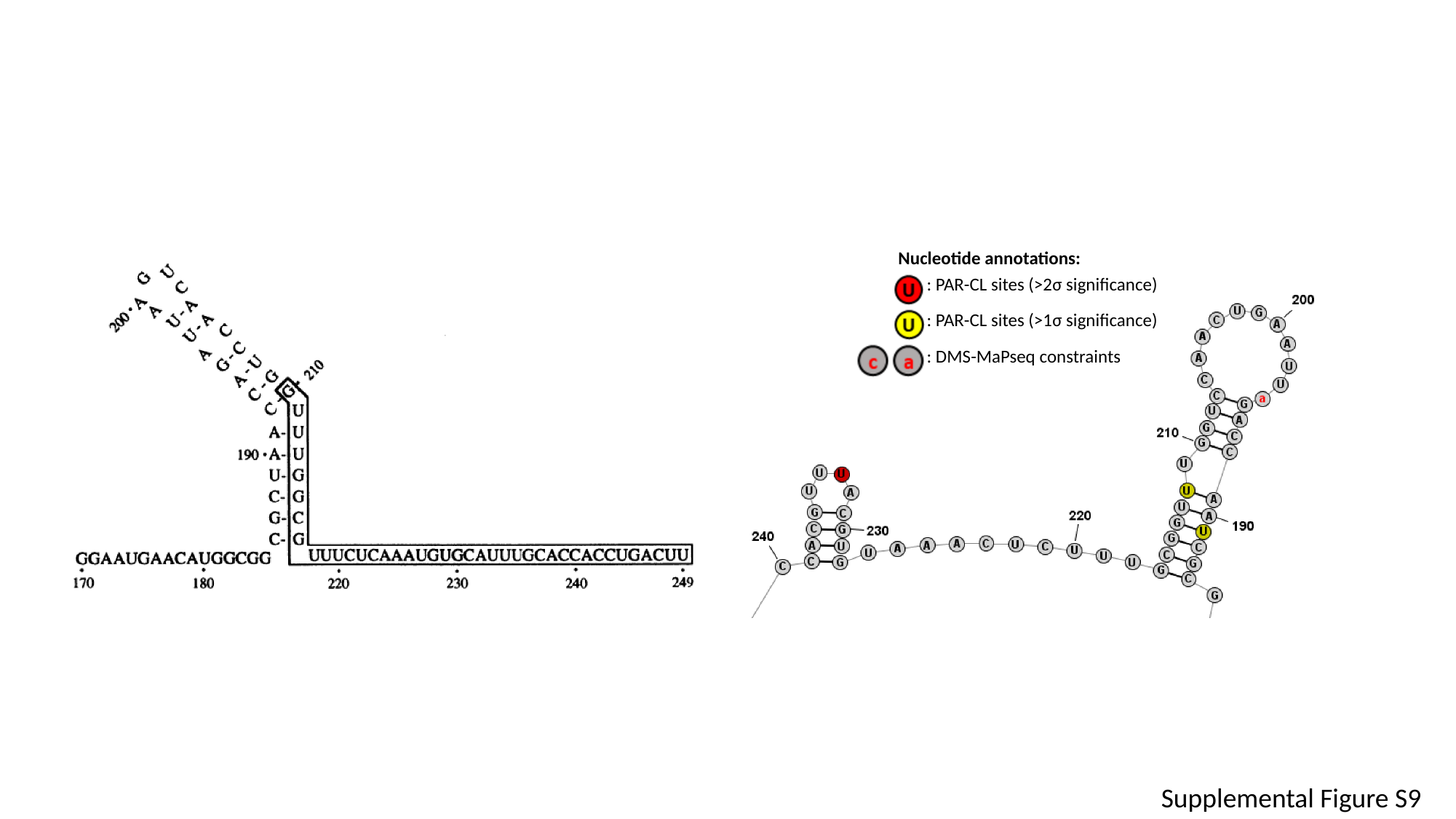

Nucleotide annotations:
| | : PAR-CL sites (>2σ significance) |
| --- | --- |
| | : PAR-CL sites (>1σ significance) |
| | : DMS-MaPseq constraints |
Supplemental Figure S9

### Slide 23
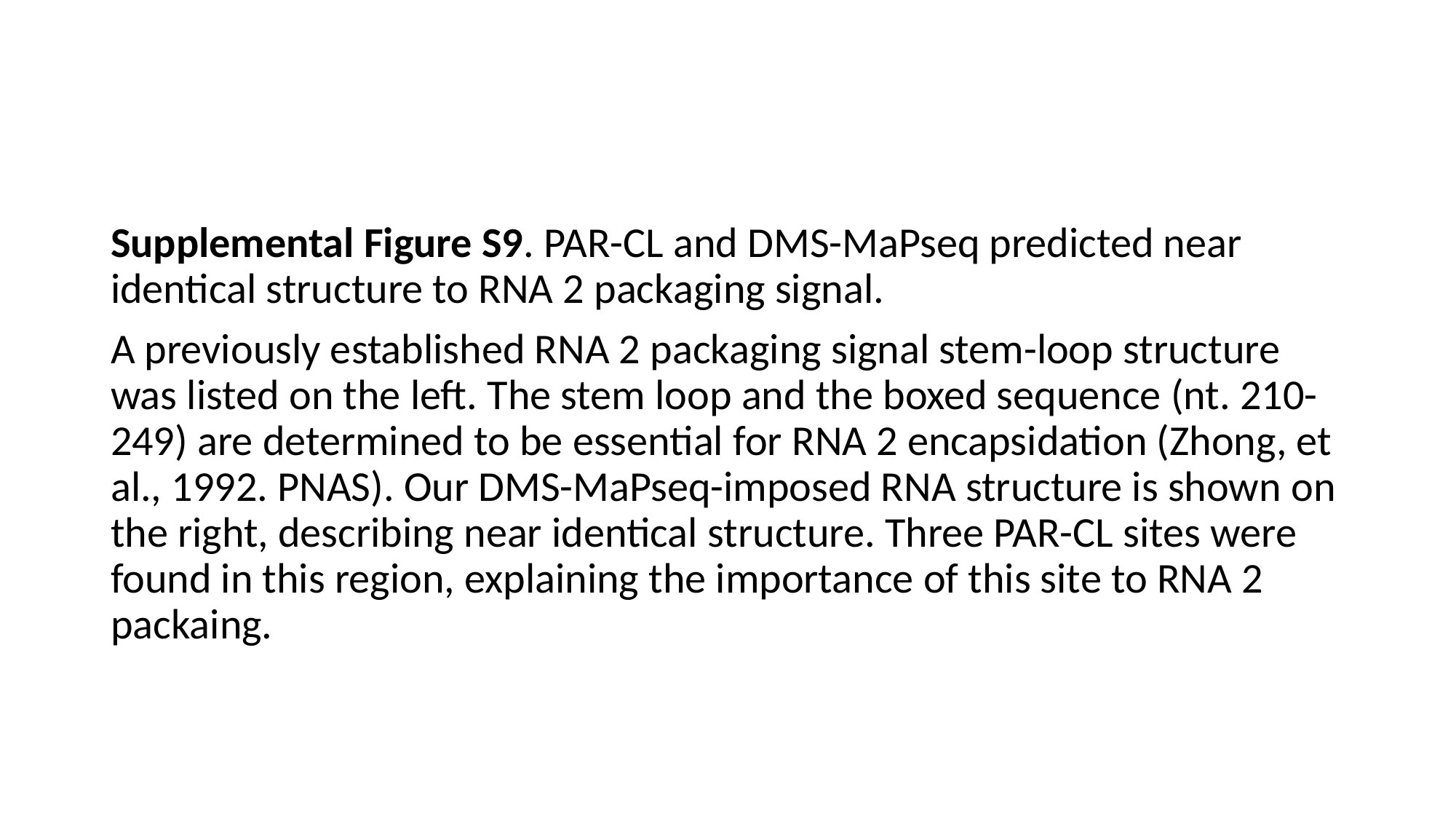

Supplemental Figure S9. PAR-CL and DMS-MaPseq predicted near identical structure to RNA 2 packaging signal.
A previously established RNA 2 packaging signal stem-loop structure was listed on the left. The stem loop and the boxed sequence (nt. 210-249) are determined to be essential for RNA 2 encapsidation (Zhong, et al., 1992. PNAS). Our DMS-MaPseq-imposed RNA structure is shown on the right, describing near identical structure. Three PAR-CL sites were found in this region, explaining the importance of this site to RNA 2 packaing.

### Slide 24
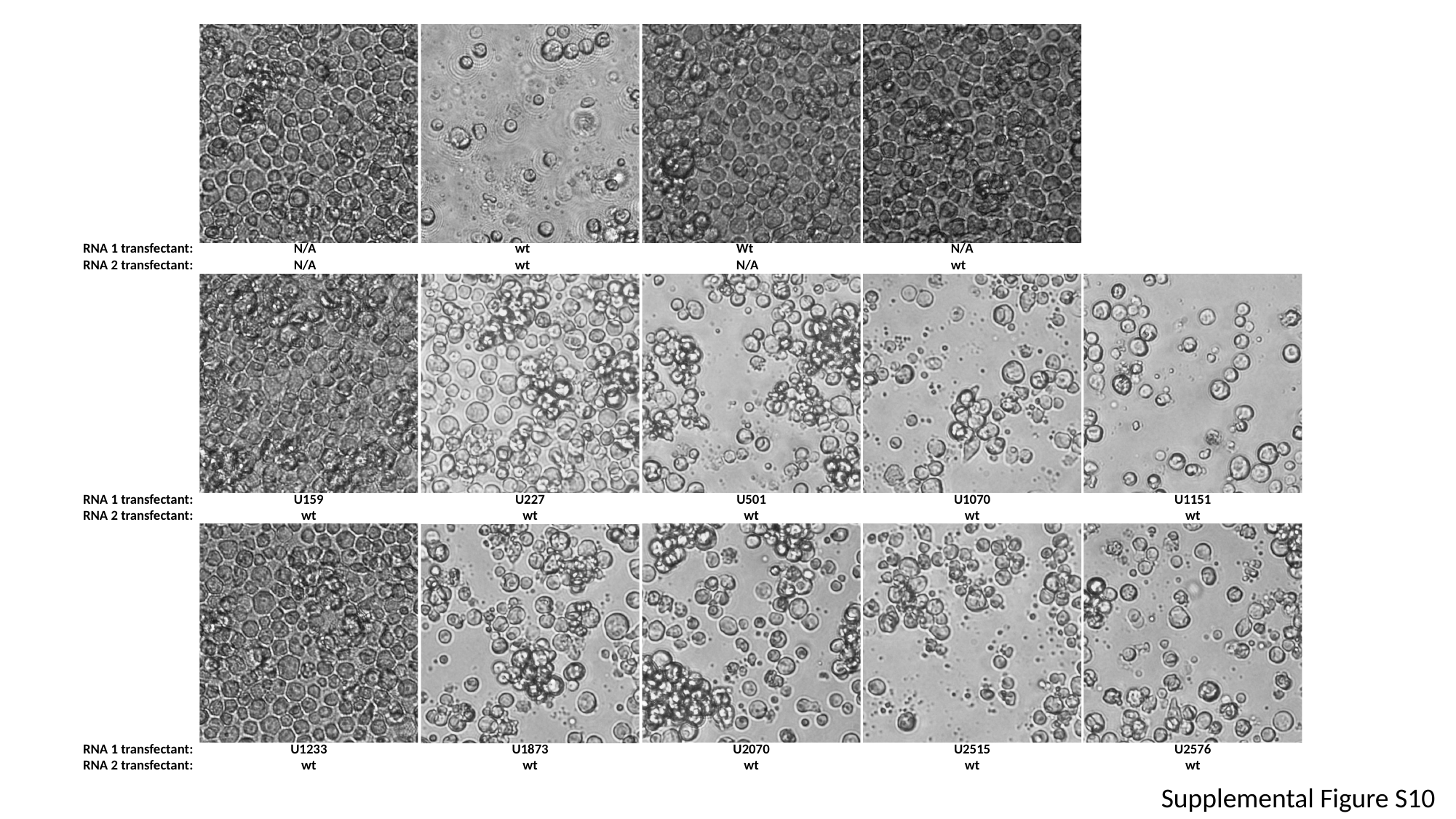

RNA 1 transfectant:
RNA 2 transfectant:
N/A
N/A
wt
wt
Wt
N/A
N/A
wt
RNA 1 transfectant:
RNA 2 transfectant:
U159
wt
U227
wt
U501
wt
U1070
wt
U1151
wt
RNA 1 transfectant:
RNA 2 transfectant:
U1233
wt
U1873
wt
U2070
wt
U2515
wt
U2576
wt
Supplemental Figure S10

### Slide 25
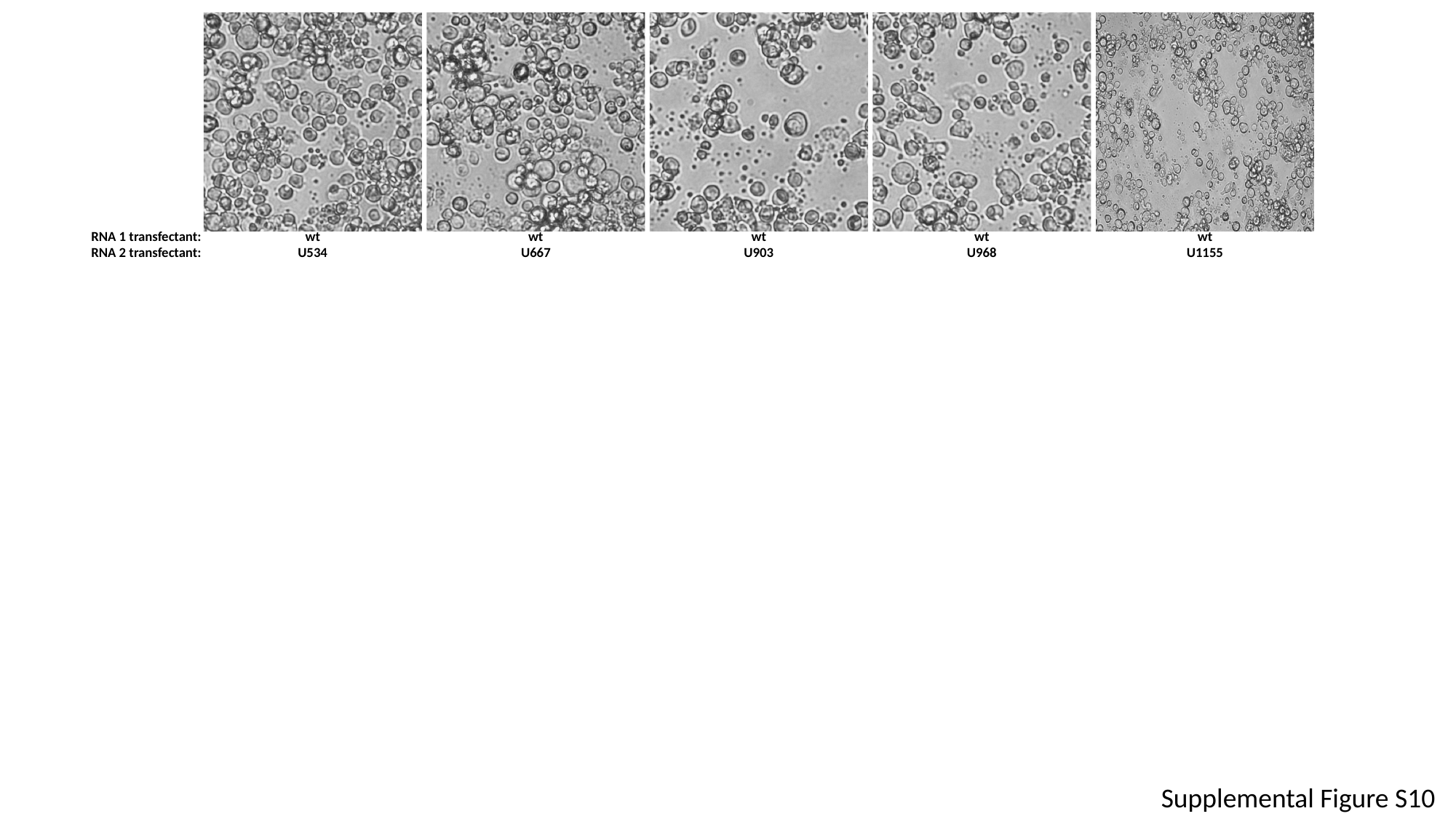

RNA 1 transfectant:
RNA 2 transfectant:
wt
U534
wt
U667
wt
U903
wt
U968
wt
U1155
Supplemental Figure S10

### Slide 26
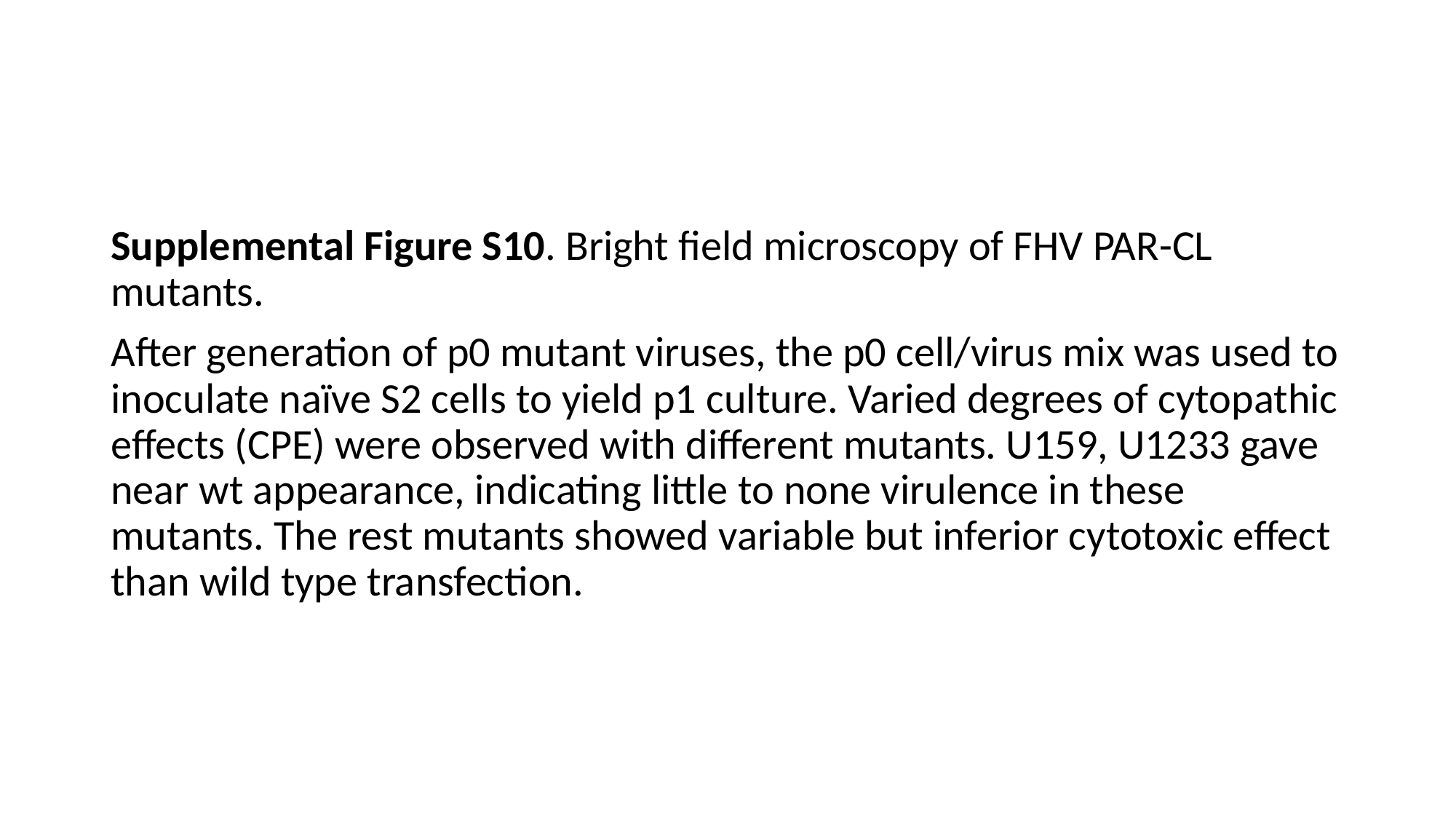

#
Supplemental Figure S10. Bright field microscopy of FHV PAR-CL mutants.
After generation of p0 mutant viruses, the p0 cell/virus mix was used to inoculate naïve S2 cells to yield p1 culture. Varied degrees of cytopathic effects (CPE) were observed with different mutants. U159, U1233 gave near wt appearance, indicating little to none virulence in these mutants. The rest mutants showed variable but inferior cytotoxic effect than wild type transfection.

### Slide 27
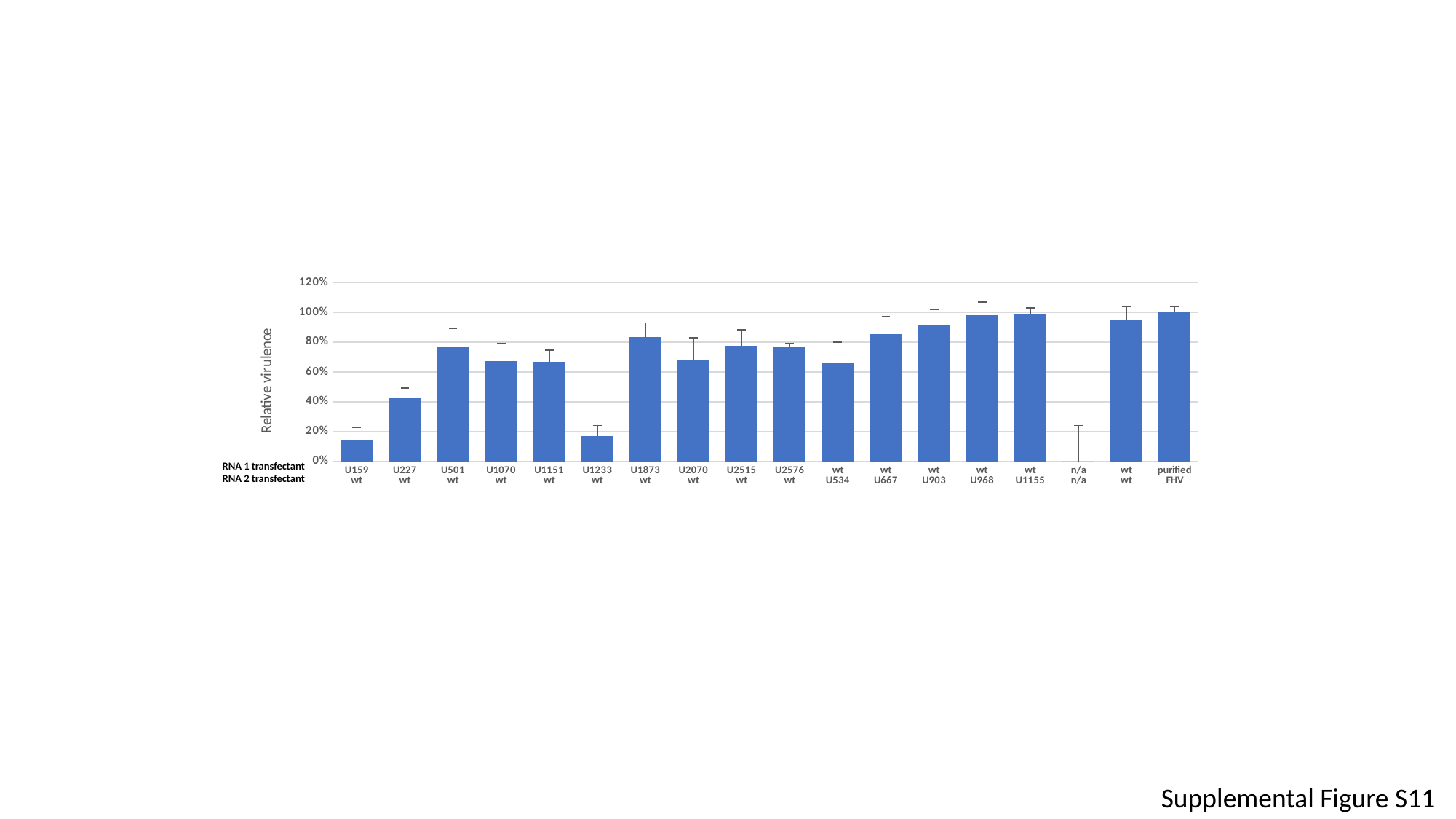

#### Chart
| Category |
|---|
| U159
wt | 0.14464968747084617 |
| U227
wt | 0.42219423453680377 |
| U501
wt | 0.7701278104300774 |
| U1070
wt | 0.6731504804552664 |
| U1151
wt | 0.6697440881114349 |
| U1233
wt | 0.1698852504897845 |
| U1873
wt | 0.832913518052057 |
| U2070
wt | 0.6851851851851852 |
| U2515
wt | 0.7751862649821833 |
| U2576
wt | 0.7639131843213476 |
| wt
U534 | 0.6593898684578786 |
| wt
U667 | 0.8520138214015764 |
| wt
U903 | 0.9194514413657991 |
| wt
U968 | 0.9789625897938241 |
| wt
U1155 | 0.9918835712286594 |
| n/a
n/a | 0.0 |
| wt
wt | 0.9496454213207414 |
| purified
FHV | 1.000069969213546 |RNA 1 transfectant
RNA 2 transfectant
Supplemental Figure S11

### Slide 28
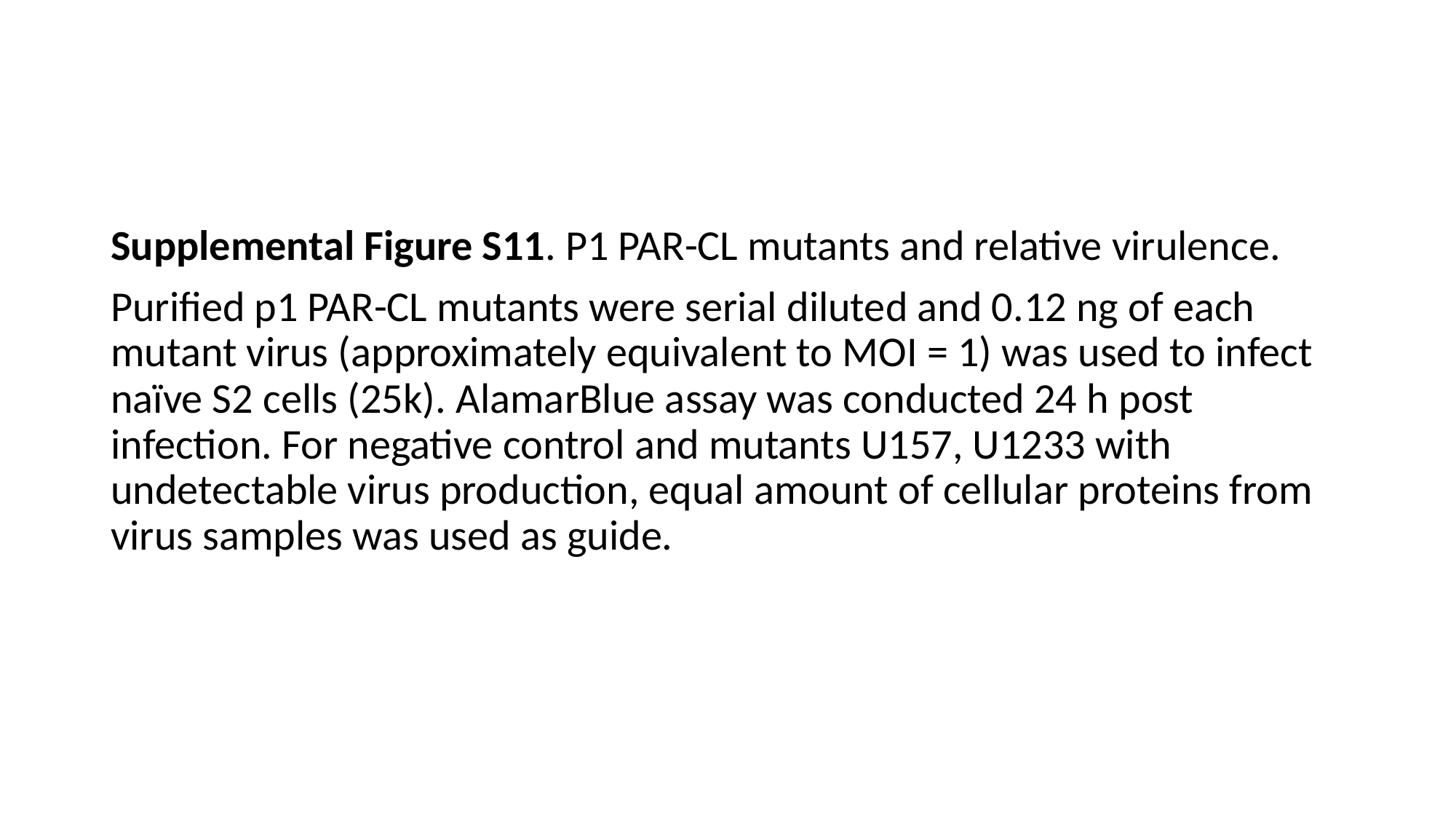

#
Supplemental Figure S11. P1 PAR-CL mutants and relative virulence.
Purified p1 PAR-CL mutants were serial diluted and 0.12 ng of each mutant virus (approximately equivalent to MOI = 1) was used to infect naïve S2 cells (25k). AlamarBlue assay was conducted 24 h post infection. For negative control and mutants U157, U1233 with undetectable virus production, equal amount of cellular proteins from virus samples was used as guide.
